# Standardized Parts for Activation of Small GTPase Signaling in Living Cells

**DOI:** 10.1101/2024.01.03.574079

**Authors:** Yuchen He, Benjamin M. Faulkner, Meaghan A. Roberti, Dana K. Bassford, Cliff I. Stains

## Abstract

Small GTPases comprise a superfamily of over 167 proteins in the human genome and are critical regulators of a variety of pathways including cell migration and proliferation. Despite the importance of these proteins in cell signaling, a standardized approach for controlling small GTPase activation within living cells is lacking. Herein, we report a split-protein-based approach to directly activate small GTPase signaling in living cells. Importantly, our fragmentation site can be applied across the small GTPase superfamily. We highlight the utility of these standardized parts by demonstrating the ability to directly modulate the activity of four different small GTPases with user-defined inputs, providing a plug and play system for direct activation of small GTPases in living cells.

## Main Text

Small GTPases have established roles in cell migration, morphogenesis, and proliferation, among several other functions (*1–3*). The activity of these proteins is regulated by a molecular switch that cycles between two distinct states: a GTP-bound active state or a GDP-bound inactive state (*4*). Cycling between these states is primarily regulated by guanine nucleotide exchange-factors (GEFs), GTPase-activating proteins (GAPs), and guanine nucleotide dissociation inhibitors (GDIs) (*5*). Exchange of GDP for GTP induces a conformational change in the small GTPase that increases its binding affinity to downstream effector proteins, leading to pathway activation (Fig. 1A). Single amino acid mutations can result in preferential binding to GTP (e.g. Q61L in Ras) or GDP (e.g. S17N in Ras) yielding constitutively active or dominant negative mutants, respectively, and are often associated with human disease (*6–8*). Overexpression of such mutants has been used to infer the role of small GTPases in signaling, but these methods can suffer from issues associated with long-term overexpression of signaling proteins, such as compensatory changes in signaling that obscure the function of the target protein (*9, 10*).

**Fig. 1.**
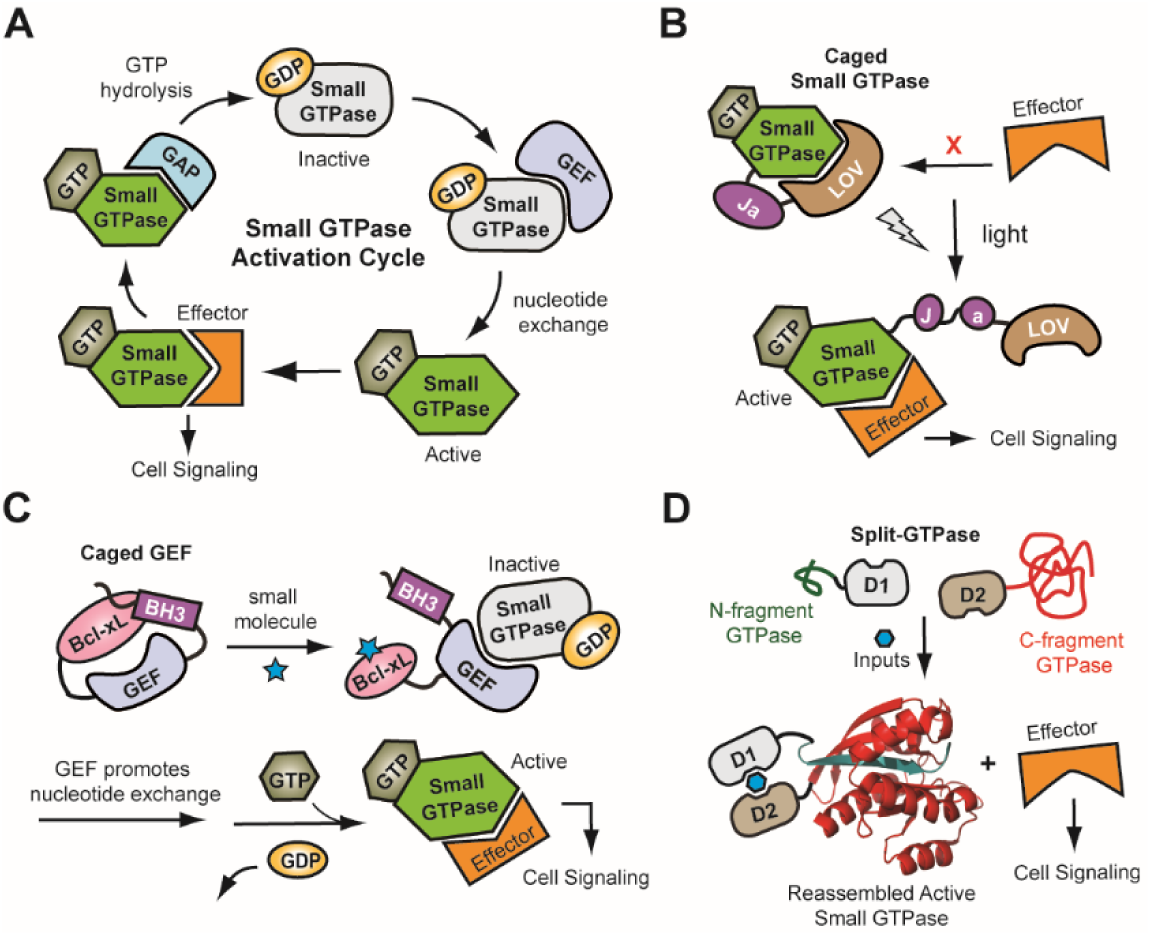
Protein engineering approaches to control small GTPase activity. (**A**) A general schematic of the small GTPase activation cycle is shown where GEF activity leads to GTP loading and GAP activity leads to hydrolysis of GTP to GDP. The GTP-bound small GTPase is capable of binding to effector proteins and promoting a signaling response. (**B**) An engineered LOV domain is used to sterically block binding of the small GTPase to an effector, rendering this interaction sensitive to light. (**C**) An engineered protein-protein interaction (PPI) is used to block GEF activity. Disruption of the PPI leads to activation of downstream small GTPases. (**D**) An increase in the local concentration of split-small GTPase fragments, using a user-defined input, leads to reassembly of an active protein.

Given the important roles of small GTPases in cellular signaling, longstanding efforts have been focused on developing small molecule modulators of these proteins for both fundamental biomedical research as well as therapeutic intervention. These efforts have largely been stymied due to the tight affinity of small GTPases for GTP (low picomolar) coupled with its relatively high concentration in cells (*11, 12*). Recent efforts have focused on covalent modification of residues adjacent to the GTP binding pocket and have achieved remarkable success (*13–15*). Indeed, compounds based on this approach have now entered the clinic (*16*). Nonetheless, this strategy relies on the presence of disease-related mutations that introduce covalently ligandable amino acids near the GTP binding site (e.g. KRas^G12C^ and KRas^G12S^) (*13, 14*). Thus, while powerful for developing inhibitors of these mutant proteins, the use of this approach to broadly study small GTPase function remains unclear. Moreover, small molecule inhibition strategies generally result in a loss of phenotype which can reduce the sensitivity of analytical assays used to detect the function of the target small GTPase. Consequently, protein engineering strategies have garnered increased attention for their potential to temporally control the activation of small GTPases.

Initial studies by Hahn and colleagues highlighted the capacity to control the activity of Rac1 by fusion of a LOV domain in close proximity to the effector binding site of a constitutively active Rac1 mutant, rendering it inactive (*17*). Subsequent irradiation with light causes destabilization of the LOV domain, allowing for binding to downstream effectors (Fig. 1B). The use of light to directly activate Rac1 activity enables high temporal and spatial resolution within living cells and organisms (*18, 19*). While powerful, this approach relies on noncovalent binding between the small GTPase and appended LOV domain to effectively inhibit activity in the dark state. Thus, the nature of this binding interface is critical for translation of this technology across the small GTPase family. Indeed, initial experiments demonstrated the need to re-engineer this interface in Cdc42 to develop a LOV-regulated variant. Therefore, while powerful, this approach requires careful case-by-case optimization for each small GTPase. Alternative approaches have centered on devising techniques to activate small GTPases regulators. For example, Maly and coworkers described an approach in which protein-protein interaction (PPI) domains are fused to a GEF leading to steric blockage of its active site (Fig. 1C) (*20, 21*). The addition of a small molecule disrupting the engineered PPI allows for restoration of GEF activity and subsequent activation of a downstream small GTPase. While powerful, this approach is hindered by the cross-reactivity of GEFs. For example, Son of sevenless (Sos-1), a well-known Ras GEF, has also been demonstrated to display Rac1-GEF activity in metazoan cells (*22, 23*). Consequently, there remains a need for development of technologies that can be applied across the small GTPase superfamily in a plug and play fashion.

To address this need, we set out to develop standardized parts for controlling the activity of virtually any small GTPase. Due to the high degree of sequence and structural homology in the small GTPase superfamily, we hypothesized that split-protein reassembly, the concentration-induced folding of a fragmented protein (*24*), could be leveraged to accomplish this goal (Fig. 1D). As a first step, we discovered the first reported fragmentation site for a small GTPase, Cdc42 (*25*). This site, termed N12/13C, facilitates small molecule-driven reassembly of a constitutively active Cdc42 mutant, enabling gated effector binding *in vitro*. In this work, we demonstrate the ability of our split-Cdc42 constructs to directly control Cdc42 signaling within living mammalian cells. Through sequence alignment, we applied our fragmentation site to a panel of GTPases, including Rac1, RhoA, and KRas. In each case, the resulting fragmented protein constructs can be used to directly control the activity of the target small GTPase within living cells without the need for optimization. Moreover, we show that these standardized parts can be interchanged with user-defined chemical-inducers of dimerization (CID) domains to tailor systems to the specific needs of an experiment. In the long-term, plug and play split-small GTPase systems will enable the direct investigation of signaling pathways within cells as well as the design of new pathways in synthetic biology applications.

### Controlling filopodia formation using split-Cdc42

We have previously identified a fragmentation site between residues 12 and 13 in the phospho-binding loop (P-loop) of Cdc42 that can be used to gate the effector binding potential of constitutively active Cdc42 *in vitro* (*25*). To investigate the ability of these fragments to gate Cdc42 signaling in living cells, we designed split-Cdc42 constructs for mammalian expression by fusing mCerulean to the N-terminus of FRB-Cdc42/N12 and mVenus followed by a CAAX box to the C-terminus of Cdc42/13C-FKBP in a vector containing an internal ribosome entry site (Fig. S1 and Tables S1 and S2). The use of a CAAX box localizes the C-terminal fragment of Cdc42 to the cytosolic face of the cellular membrane, which corresponds to the localization of the active, native protein (*26*). Constructs for two different split-Cdc42 mutants were generated corresponding to the constitutively active protein, Q61L, or dominant negative protein, T17N. Importantly, the Q61L mutant decouples split-Cdc42 signaling from endogenous regulatory pathways (e.g. GAP and GEF), since this mutant displays preferential binding to GTP and impaired GTP hydrolysis, providing a positive control (*27*). Conversely, the dominant negative mutant (T17N), displays preferential binding to GDP and serves as a negative control (*28*). Transient expression of the constructs in HeLa cells demonstrated the anticipated localization of each protein fragment (Fig. S2).

To investigate the ability of split-Cdc42 to reassemble within mammalian cells in response to a small molecule input, we treated transiently transfected CHO cells with 1 µM rapamycin for 20 min, followed by cell lysis and pull-down of reassembled Cdc42 using a known effector protein (Fig. 2A, B, and C). Given that rapamycin was not present in buffers used for cell lysis or the pull-down experiment, these results clearly indicate that reassembly of split-Cdc42-Q61L occurred in cells prior to lysis. Moreover, no pull-down of reassembled, dominant negative split- Cdc42-T17N was observed. Encouraged by this result, we next asked whether split-Cdc42 constructs could be used to control a known cellular phenotype of Cdc42 activity, namely filopodia formation at the cellular periphery (*29*). Filopodia are antenna-like protrusions that serve diverse functions in directional cell migration, wound healing, and cellular communication (*30–32*). We envisioned that reassembly of split-Cdc42-Q61L would lead to activation of the known Cdc42 effectors p21-activated kinases (PAKs) and the Wiskott-Aldrich syndrome protein (WASP), ultimately leading to filopodia assembly (Fig. 2D) (*33*). To test this, HeLa cells were transiently transfected with our split-Cdc42 constructs and either stimulated with 500 nM rapamycin or DPBS for 30 min. Filopodia formation was then imaged using confocal microscopy (Fig. 2E) and the number of filopodia detected per cell was quantified using FiloQuant (Fig. 2F) (*34*). Increased filopodia formation was only observed in split-Cdc42-Q61L expressing cells treated with rapamycin, leading to a 4.1-fold increase in filopodia formation. In contrast, the dominant negative split-Cdc42-T17N constructs did not influence filopodia formation in the absence or presence of rapamycin. Taken together, these results clearly demonstrate the ability to directly control Cdc42 signaling within living cells using split-protein reassembly.

**Fig. 2.**
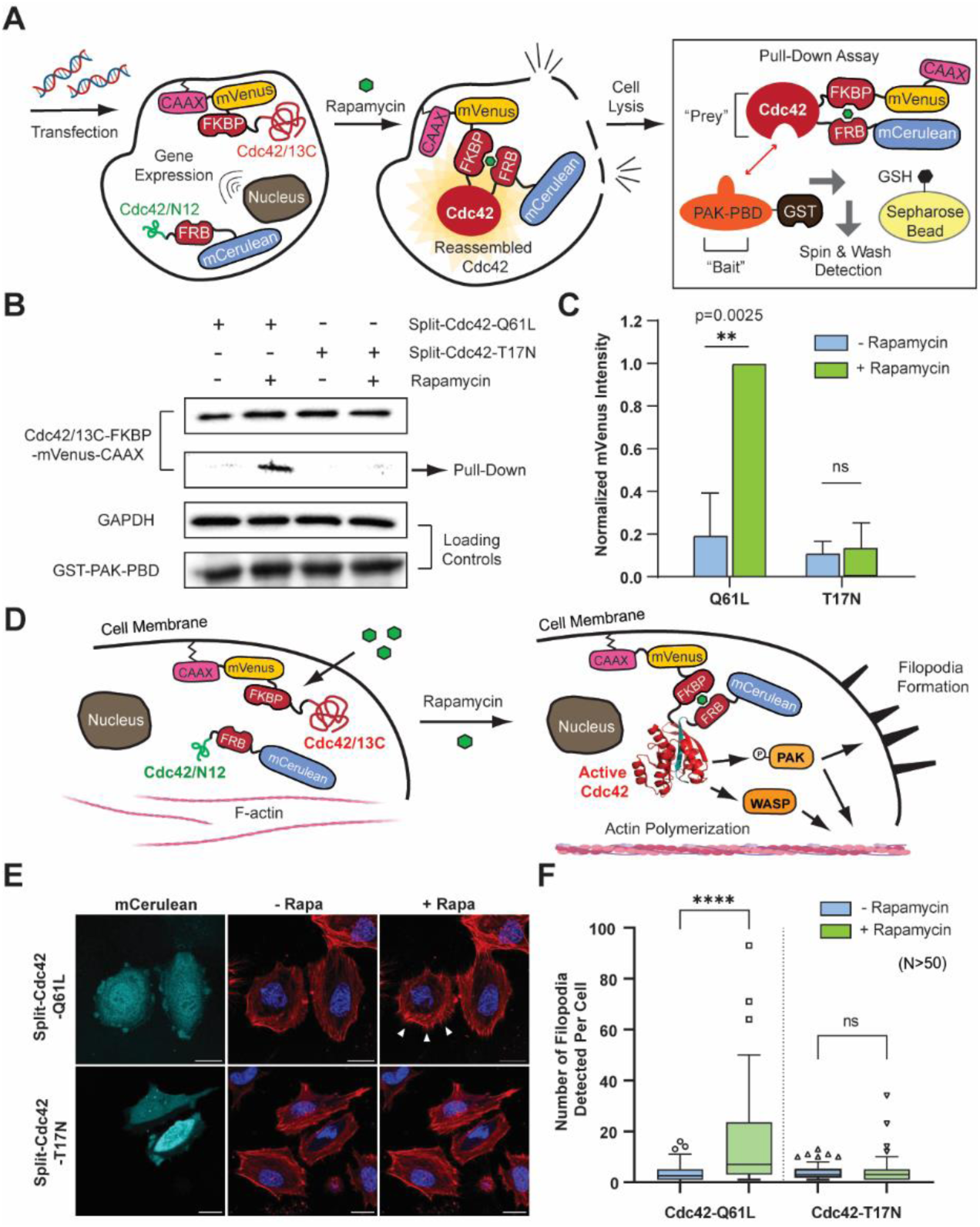
Gated control of filopodia formation in living mammalian cells using split-Cdc42. (**A**) A schematic depicting the procedure used to pull-down split-Cdc42 that reassembled in cells. Cells are treated with rapamycin and subsequently washed, lysed, and assayed using an immobilized Cdc42 effector. Rapamycin is not present during cell lysis or the subsequent pull-down assay. (**B**) A representative western blot for pull-down assays probing for reassembled split-Cdc42 in CHO cells. Equivalent amounts of PAK-PBD and cell lysates were used, see loading controls. Split-Cdc42-Q61L is significantly enriched in cells that were treated with 1 µM rapamycin before lysis. (**C**) Quantification of western blots from panel **B** using three biological replicates. (**D**) A schematic showing the expression of split-Cdc42 in mammalian cells and activation of Cdc42 signaling via rapamycin, leading to filopodia formation. (**E**) Confocal images of HeLa cells in the mCerulean channel (representing transfected cells) or merged images of CellMask Deep Red Actin tracker (red) and Hoechst stain (blue) in the absence or presence of 500 nM rapamycin for 30 min (-Rapa or +Rapa, respectively). Cells expressing split-Cdc42- Q61L display clear filopodia formation upon rapamycin stimulation (arrows), while cells expressing split-Cdc42-T17N do not. Nontransfected cells within the same field of view serve as internal controls for the effect of rapamycin. (**F**) Quantified filopodia number from panel **E** for N > 50 cells using FiloQuant shows a 4.1-fold increase in the number of filopodia formed in rapamycin treated cells expressing split-Cdc42-Q61L. Data are means ± SD. Statistical significance was determined using a two-tailed, unpaired students’ t-test. The scale bar represents 20 µm. ** indicates a p-value of <0.01, **** indicates a p-value of <0.0001, and ns indicates a p-value of >0.05.

### Using sequence alignment to generate split-protein constructs for target small GTPases

To apply our fragmentation site to other GTPases, we first sought to develop a relatively rapid *in vitro* assay for fragmented GTPase reassembly. We expressed and purified full-length Cdc42-Q61L, Cdc42-T17N, and split-Cdc42-Q61L fragments, appended to rapamycin-dependent interaction domains, from bacteria (Fig. S3, Fig. S4, and Tables S1 and S3). To assay for protein reassembly, we leveraged the fluorescence enhancement of mantGTP upon binding to a small GTPase, as a proxy for folding of split-GTPase fragments (Fig. 3A) (*35*). Gratifyingly, we observed a 5.5-fold increase in fluorescence of full-length Cdc42-Q61L relative to Cdc42-T17N, showcasing the assay’s ability to discern GTP binding differences between these two mutants (Fig. S5). In addition, a clear increase of 8.6-fold in fluorescence was observed for split-Cdc42-Q61L in the presence of rapamycin versus without rapamycin, indicating that this assay could report upon the folding and subsequent binding of mantGTP in the presence of rapamycin (Fig. 3B). With this relatively rapid assay for split-GTPase reassembly in-hand, we set out to test whether our N12/13C fragmentation site could be applied across the small GTPase superfamily.

**Fig. 3.**
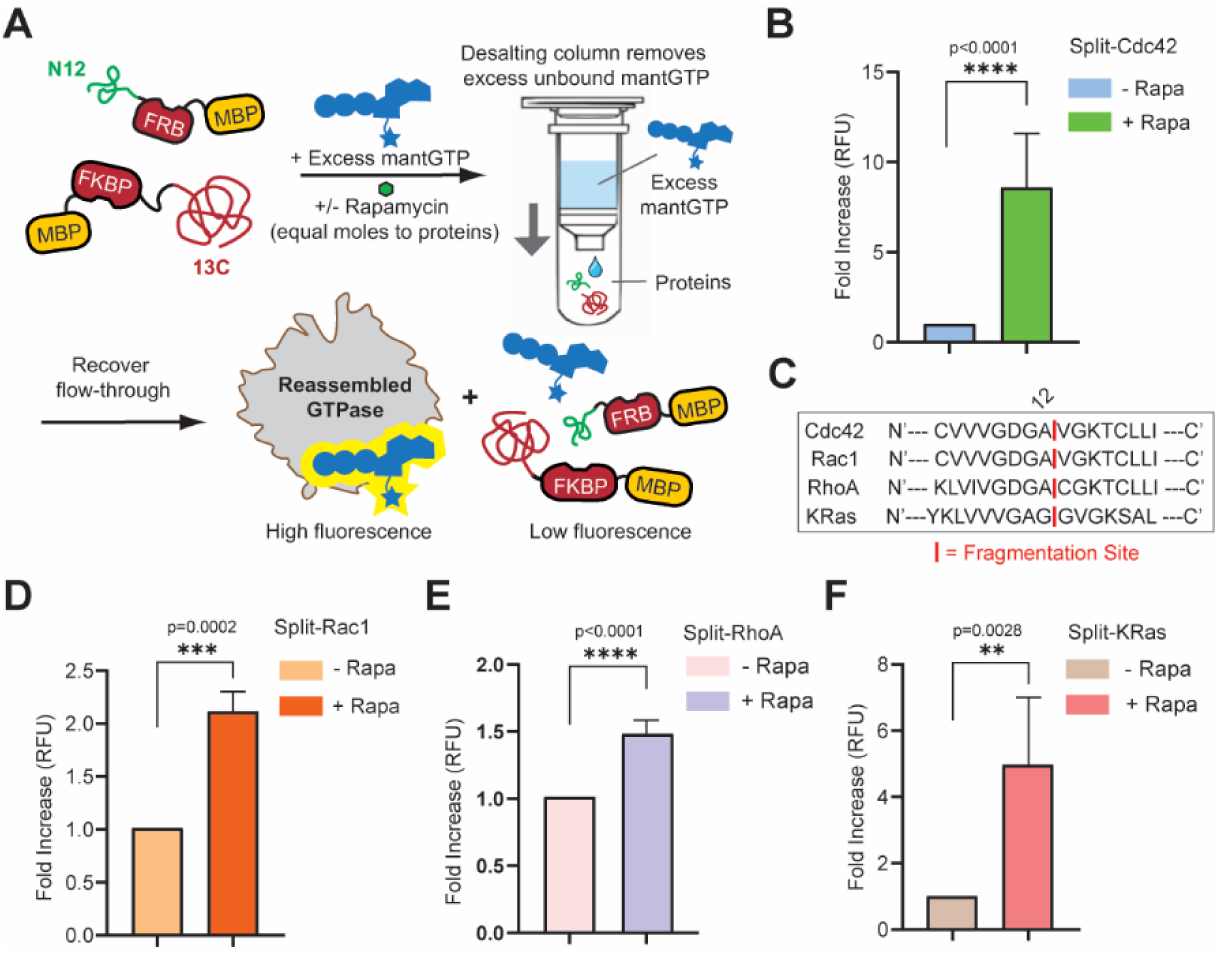
The N12/13C fragmentation site is generalizable across the small GTPase family. (**A**) A schematic showing the *in vitro* assay workflow. Equal concentrations of split-GTPase fragments (GTPase/N12 and GTPase/13C) are incubated with a stoichiometric amount of rapamycin. The reaction mixture is then incubated with a 10-fold excess of mantGTP probe, followed by locking the nucleotides in the GTPase binding pocket with excess Mg^2+^. Reactions are then desalted to remove excess, unbound mantGTP. Reassembled protein binds to mantGTP, inducing an increase in fluorescence. (**B**) Fluorescence of samples containing the indicated split-Cdc42 constructs (10 µM) in the presence or absence of rapamycin (10 µM). An 8.6-fold increase in the fluorescence of samples containing rapamycin is observed. (**C**) Sequence alignment between different small GTPases and Cdc42, Cdc42 numbering is used throughout to refer to fragmentation sites. (**D-F**) Fluorescence of samples containing split-Rac1, split-RhoA, or split- KRas constructs in the presence or absence of rapamycin. The concentrations for each fragmented protein used in the assay was: split-Rac1 (25 µM), split-RhoA (15 µM), or split-KRas (5 µM). The concentration of rapamycin was equal to the concentration of fragmented protein. A statistically significant increase in fluorescence was observed for all fragmented small GTPases in the presence of rapamycin. Data are means ± SD. ** indicates a p-value of <0.01, *** indicates a p-value of <0.001, and **** indicates a p-value of <0.0001 calculated using a two-tailed, unpaired students’ t-test.

Given the highly conserved sequence and structural homology within the small GTPase superfamily (*36*), we hypothesized that the N12/13C fragmentation site, located within the P-loop (*25*), could be extended to other small GTPases. To test this hypothesis, we selected a panel of three other small GTPases with well-stablished biological functions: Rac1 (membrane ruffling) (*37*), RhoA (cell contraction) (*38*), and KRas (cell proliferation) (*39*). Using sequence alignment, we identified the corresponding N12/13C sites in Rac1, RhoA, and KRas (Fig. 3C). Based on these alignments, we cloned constitutively active constructs for split-Rac1, RhoA, and KRas into bacterial expression plasmids (Fig. S3, Fig. S4, and Tables S1 and S3). The resulting split-small GTPases, appended to rapamycin-dependent interaction domains, were expressed and purified, with the corresponding full-length constitutively active and dominant negative mutants serving as controls (Fig. S3, Fig. S4, and Tables S1 and S3). Although varying sensitivity was observed, the mantGTP assay was capable of distinguishing between full-length constitutively active and dominant negative GTPase variants (Fig. S5). In support of our hypothesis, we observed clear evidence for reassembly, as assessed by a statistically significant increase in mantGTP fluorescence for each new fragmented GTPase (Fig. 3D, E, and F). Importantly, identification of these new split-Rac1, -RhoA, and -KRas constructs was performed solely through sequence alignment. Thus, our N12/13C site represents a plug and play strategy for developing gated systems for a target small GTPase. Next, we set out to investigate the ability to control relevant biological signaling pathways in living cells using these new split-small GTPases.

### Gated cytoskeletal remodeling using split-Rac1 and split-RhoA systems

To express split-Rac1 or split-RhoA constructs in mammalian cells, we replaced Cdc42 fragments with the corresponding fragments of Rac1 or RhoA for both constitutively active and dominant negative mutants of each protein (Fig. S1 and Table S2). Upon transient transfection into HeLa cells, we verified localization of the fragmented proteins using confocal microscopy (Fig. S6 and Fig. S7). With these mammalian expression constructs in-hand, we first asked whether split-Rac1 was capable of activating signaling within living cells. Rac1 has an established role in cell movement and actin polymerization, specifically the regulation of lamellipodia formation (*40*). These dynamic membrane protrusions, composed of branched actin filaments, are typically found at the leading edge of cells (*41*). Moreover, crosstalk between Rac1 and Cdc42 signaling is well established and attributed to cross reactivity of GEFs as well as downstream effectors of these small GTPases (*42, 43*). For example, transfection of cells with ITSN (a Cdc42 GEF) has been shown to lead to lamellipodia formation (Rac1 phenotype) rather than the expected filopodia formation (Cdc42 phenotype) (*44*). Thus, we asked whether reassembly of split-Rac1-Q61L could induce lamellipodia formation and membrane ruffling (Fig. 4A). To test this, we transiently transfected HeLa cells with either constitutively active split- Rac1-Q61L or dominant negative split-Rac1-T17N, followed by stimulation with rapamycin. Upon treatment with rapamycin, we observed a 5.4-fold increase in membrane ruffling at the cell periphery for cells transfected with split-Rac1-Q61L across biological replicates. In contrast, a 2.3-fold increase in membrane ruffling was observed in cells expressing split-Rac1-T17N in the presence of rapamycin and could be attributed to previously observed changes in membrane morphology induced by activation of Rac1-T17N (*17*) (Fig. 4, B, C, and D). Notably, compared with the thin, finger-like filopodia formed upon direct activation of Cdc42 (Fig. 2E), protrusions from Rac1 activation were broader and displayed a sheet-like spreading structure (Fig. 4C). These results clearly demonstrate the ability to directly control Rac1 signaling within mammalian cells using our plug and play system. Moreover, these results highlight the utility of direct activation of small GTPases within living cells as means to avoid signaling cross-talk.

**Fig. 4.**
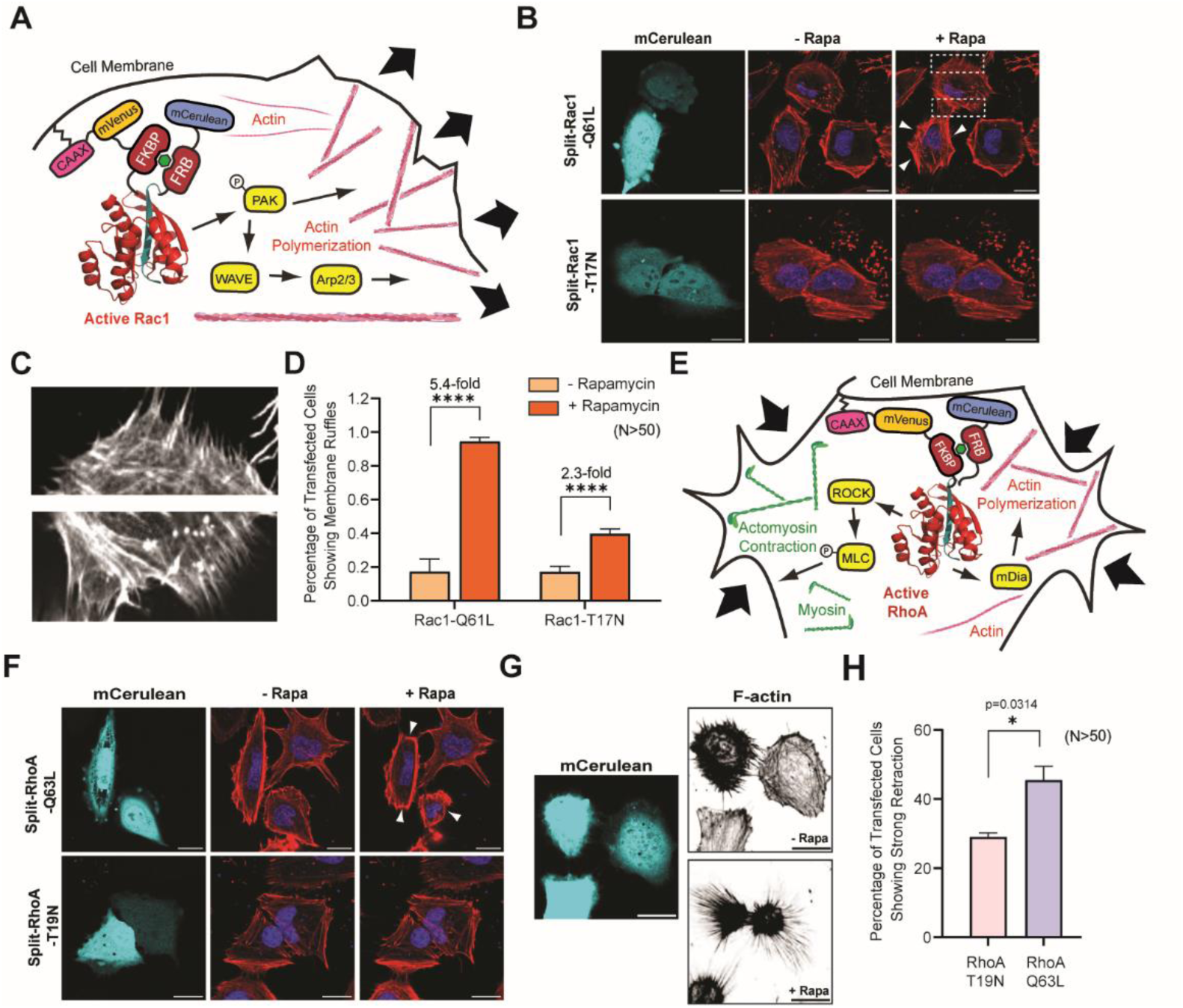
Gated control of Rac1 and RhoA signaling in living mammalian cells. (**A**) A schematic illustrating activation of split-Rac1 leading to membrane ruffling. (**B**) Confocal images of HeLa cells in the mCerulean channel (representing transfected cells) or merged images of CellMask Deep Red Actin tracker (red) and Hoechst stain (blue) in the absence or presence of 500 nM rapamycin (-Rapa or +Rapa, respectively). Cells expressing active split-Rac1-Q61L demonstrate pronounced membrane ruffling (dashed rectangle) after stimulation with rapamycin. Nontransfected cells within the same field of view serve as internal controls for the effect of rapamycin. (**C**) Magnified images of the cell periphery highlighted by dashed rectangles in **B**. (**D**) Quantified membrane ruffling from N > 50 cells in each sample. Membrane ruffling was scored by two separate, blinded raters and used to obtain inter-rater reliability for the percent of membrane ruffling. Across all samples, the median Cohen’s Kappa value was 0.82, indicative of strong agreement between independent raters. (**E**) A schematic of split-RhoA signaling leading to membrane retraction. (**F**) Confocal images of HeLa cells expressing the indicated split-RhoA construct in the absence or presence of 500 nM rapamycin for 30 min (-Rapa or +Rapa, respectively). Cells expressing active split-RhoA-Q63L demonstrate pronounced membrane retraction (arrows) after stimulation with rapamycin. Nontransfected cells within the same field of view serve as internal controls for the effect of rapamycin. (**G**) Confocal images of HeLa cells expressing split-RhoA-Q63L in the mCerulean channel (representing transfected cells) or stained for F-actin in the absence or presence of 500 nM rapamycin for 30 min (-Rapa or +Rapa, respectively). Clear formation of contractile F-actin filaments is observed in the presence of rapamycin. (**H**) Quantification of transfected cells from panel **F** demonstrating substantial cell retraction, defined as > 20% of the total cell area as measured using ImageJ. Data are means ± SD. Statistical significance was determined using a two-tailed, unpaired students’ t-test. * indicates a p-value of <0.05 and **** indicates a p-value of <0.0001. Scale bars represent 20 µm.

Next, we evaluated the ability to directly control RhoA signaling within living cells. RhoA regulates myosin-mediated cell contraction and plays an important role in signaling at the trailing edge of the cell (*38*). RhoA exerts these effects through activation of ROCK-mediated actomyosin contraction as well as actin polymerization (Fig. 4E) (*45*). HeLa cells expressing constitutively active split-RhoA-Q63L showed distinct cell membrane retraction upon rapamycin treatment (Fig. 4F). Additionally, in cells expressing split-RhoA-Q63L, rapamycin treatment resulted in elongated, contractile F-actin filaments converging towards the nucleus (Fig. 4G). Importantly, these effects were much less pronounced in the absence of rapamycin or in cells expressing split-RhoA-T19N, regardless of the presence or absence of rapamycin (Fig. 4F). Moreover, consistent rapamycin-induced membrane retraction was observed in cells expressing split-RhoA-Q63L (Fig. 4H). These data confirm the ability to directly activate RhoA signaling using split-RhoA constructs, further validating the generality of our plug and play approach.

### Direct regulation of KRas signaling using gibberellic acid

One of the most studied pathways downstream of KRas is the mitogen-activated protein kinase (MAPK) cascade, a key regulator of cell proliferation, differentiation, and survival (*39*). Having demonstrated the ability to control split-KRas reassembly *in vitro* (Fig. 3F), we asked whether rapamycin could be used to gate the activity of split-KRas in mammalian cells by monitoring downstream phosphorylation of ERK (Fig. S8A). Prior to assays, we verified equivalent expression of each construct (Fig. S8B). HeLa cells transiently transfected with split- KRas-Q61L were stimulated with 500 nM rapamycin, lysed at different time points, and p-ERK levels were assessed by western blotting. Interestingly, we observed a clear increase in p-ERK within 1 minute of stimulation with rapamycin that was sustained out to 12 minutes post stimulation while no increase in p-ERK was observed for cells expressing split-KRas-S17N (Fig. S8C, D and E). However, we also observed an increase in p-ERK levels upon stimulation of non-transfected cells that was shorter in duration compared to cells expressing split-KRas-Q61L (Fig. S8C, D and E). Indeed, rapamycin is a known inhibitor of mTOR signaling and has been reported to increase p-ERK levels due to crosstalk between these pathways (*46, 47*). We hypothesize that the effect of rapamycin on activation of ERK is muted in cells expressing split- KRas-S17N due to the dominant negative inhibitory effect of this protein (*48*). These results highlight the need for a plug and play approach for controlling small GTPase activity since inputs for gating small GTPase activity may need to be modified based on the specific requirements of an experiment. Accordingly, we asked whether an alternative CID system, devoid of any known interaction with ERK signaling, might be suitable for gating split-KRas activity.

We turned to the well-characterized gibberellic acid-gated CID system that relies on the gibberellic acid mediated interaction of GAI and GID1 (*49*). Replacing FKBP and FRB in Fig. S8A with GAI and GID1, respectively, yielded gibberellic acid gated, split-KRas constructs for expression in mammalian cells (Fig. S1 and Tables S1 and S2) that we envisioned could be used to control ERK activation (Fig. 5A). As with our rapamycin-dependent split-KRas constructs, confocal imaging demonstrated appreciate localization of GAI and GID1 fusions within cells as well as equivalent transfection efficiency for split-KRas-Q61L versus split-KRas- S17N constructs (Fig. S9). We then stimulated cells expressing split GA-KRas-Q61L with a cell permeable analog of gibberellic acid (GA_3_-AM), lysed at varying time points after stimulation, and probed for changes in p-ERK levels via western blotting. Gratifyingly, cells expressing GA- KRas-Q61L displayed a sharp 2.3-fold increase in ERK phosphorylation one minute after stimulation with GA_3_-AM that subsided to baseline by 12 minutes (Fig. 5B and C, Fig. S10). However, cells expressing split GA-KRas-S17N or non-transfected cells did not exhibit a detectable increase in ERK phosphorylation (Fig. 5B and C, Fig. S10). These experiments demonstrate the broad applicability of our N12/13C fragmentation site across the small GTPase superfamily as well as the ability to utilize user-defined inputs to gate reassembly of these fragments.

**Fig. 5.**
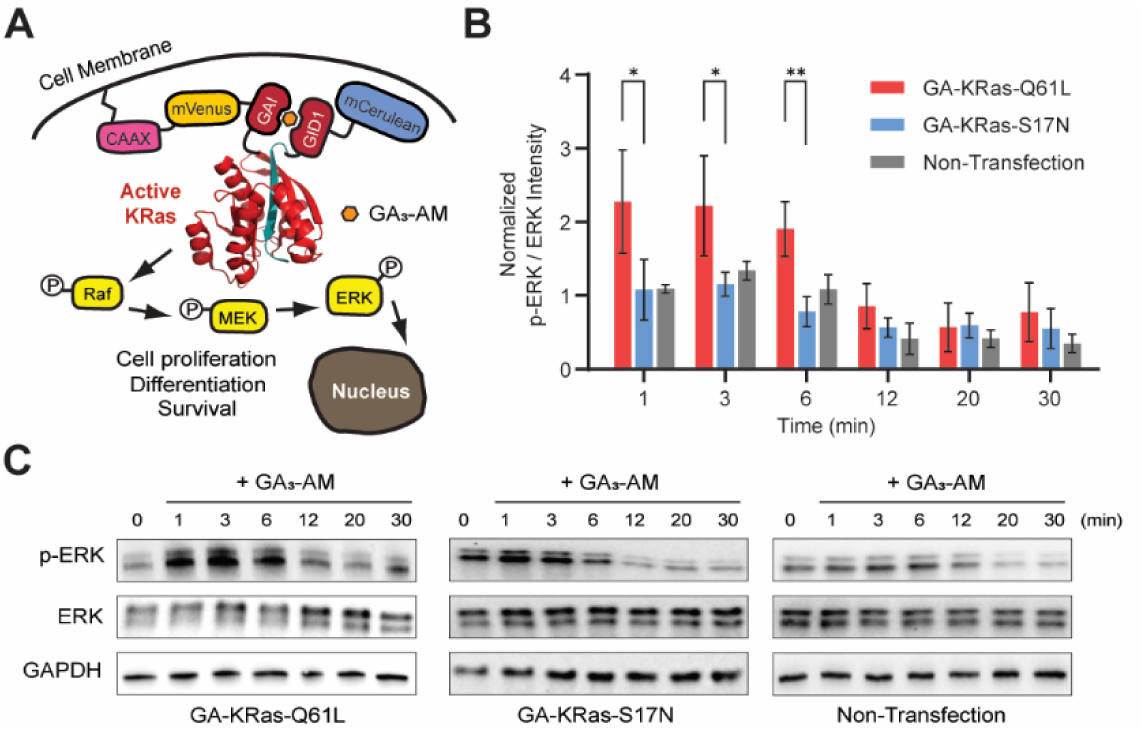
Gibberellic acid-gated split-KRas constructs for controlling ERK activation. **(A)** A schematic of gibberellic acid-gated activation of split-KRas signaling. **(B)** Quantified western blot intensities for p-ERK relative to total ERK from three independent biological replicates (mean ± SD). * Adjusted P<0.05, ** Adjusted P<0.01 using one-way ANOVA followed by Fisher’s PLSD test. **(C)** Representative western blots from HeLa cells transfected with the indicated construct and stimulated with 10 µM gibberellic acid for the indicated time. Additional western blot results are shown in Figure S10.

## Conclusions

In summary, we have developed a plug and play system for the direct activation of a target small GTPase via a user-defined input within living cells. Using sequence alignment, we demonstrate the ability to rapidly identify split-small GTPase constructs capable of controlling filopodia formation (Cdc42), membrane ruffling (Rac1), membrane retraction (RhoA), or MAPK activation (KRas). Importantly, in each case, fragmentation sites identified through sequence alignment could be used directly without the need for further optimization. Moreover, these fragmentation sites could be used interchangeably with user-defined inputs for reassembly. Thus, these plug and play split-small GTPase modules can be tailored to the specific needs of an experiment. An additional feature of this system is that it decouples small GTPase activation from endogenous regulatory pathways, thereby avoiding effects from crosstalk between upstream regulatory proteins. To demonstrate the importance of direct activation of small GTPases, we utilized direct activation of Cdc42 versus Rac1 signaling to demonstrate the difference in membrane structures formed by activation of each protein. In the long term, we believe that this technology will find broad application in the dissection of fundamental aspects of small GTPase signaling as well as synthetic biology efforts to integrate new functions into living systems.

## Acknowledgements

We thank the W.M. Keck Center for Cellular Imaging for the usage of the Leica STELLARIS 8 confocal/FLIM/tauSTED microscope system (NIH OD030409), members of the C. Stains lab for editing the manuscript and helpful conversations, and members of M. Stains lab for assistance with evaluation of inter-rater reliability. We acknowledge financial support from the NIH (R35GM148221) and the University of Virginia. The content of this work is solely the responsibility of the authors and does not necessarily represent the official views of the NIH.

## Notes

### Competing Interest Statement

The authors have declared no competing interest.

