## Supporting Information for "Standardized Parts for Activation of Small GTPase Signaling in Living Cells"

**Fig. S1**

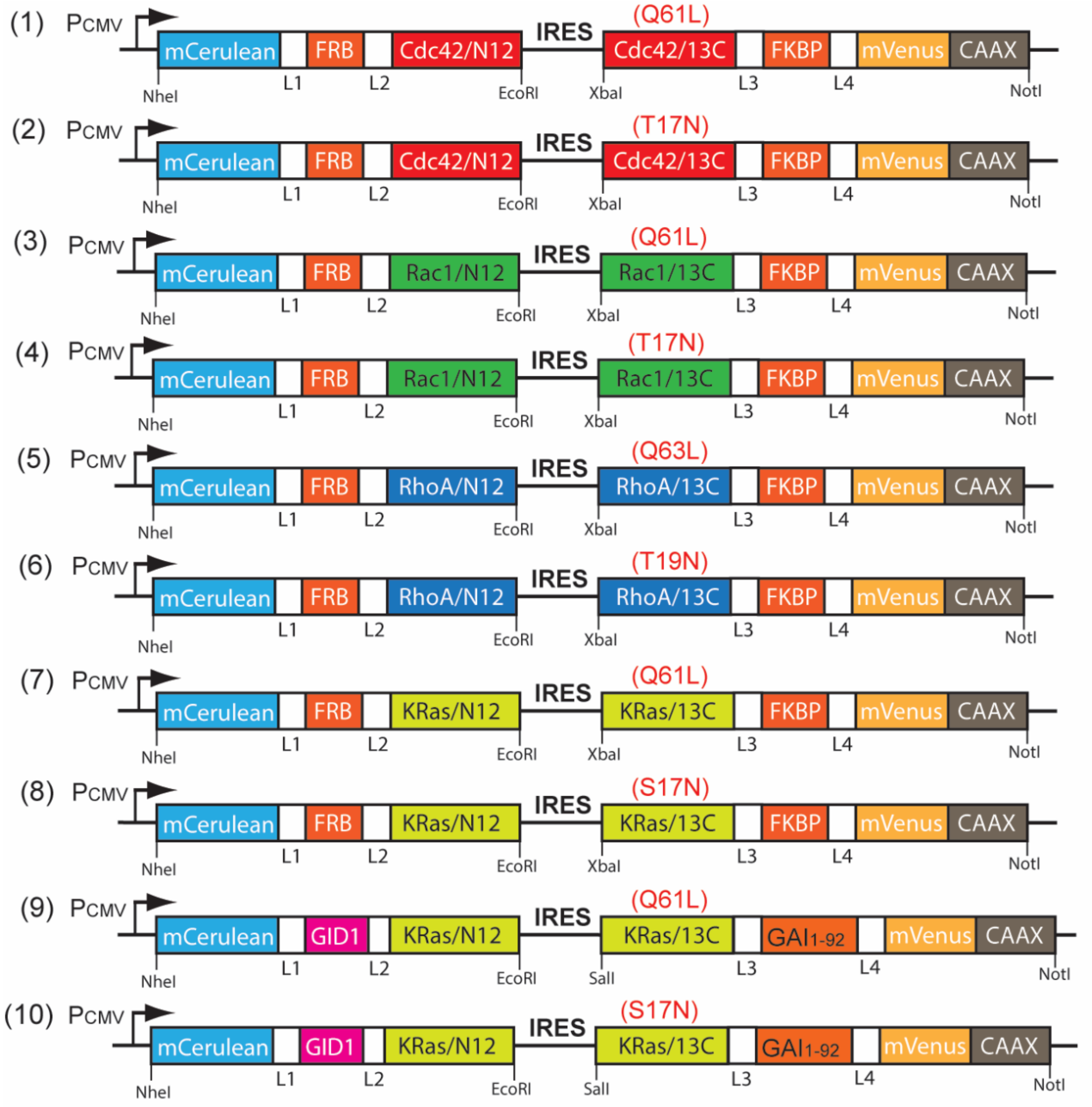

L1: GGGSSGGG
L2: QISYASRGGGSSGGGEL
L3: GDPNWELVYTARLQGGGSSGGGQISYASRG
L4: GGGSSGGG
CAAX: KKKKKKSKTKCVIM

**A schematic for protein constructs used in mammalian expression.** IRES is internal ribosomal entry site. See Table S1 for full sequence information. Q61L and Q63L represent constitutively active mutants. T17N, T19N, and S17N refer to dominant negative mutants.

**Fig. S2**

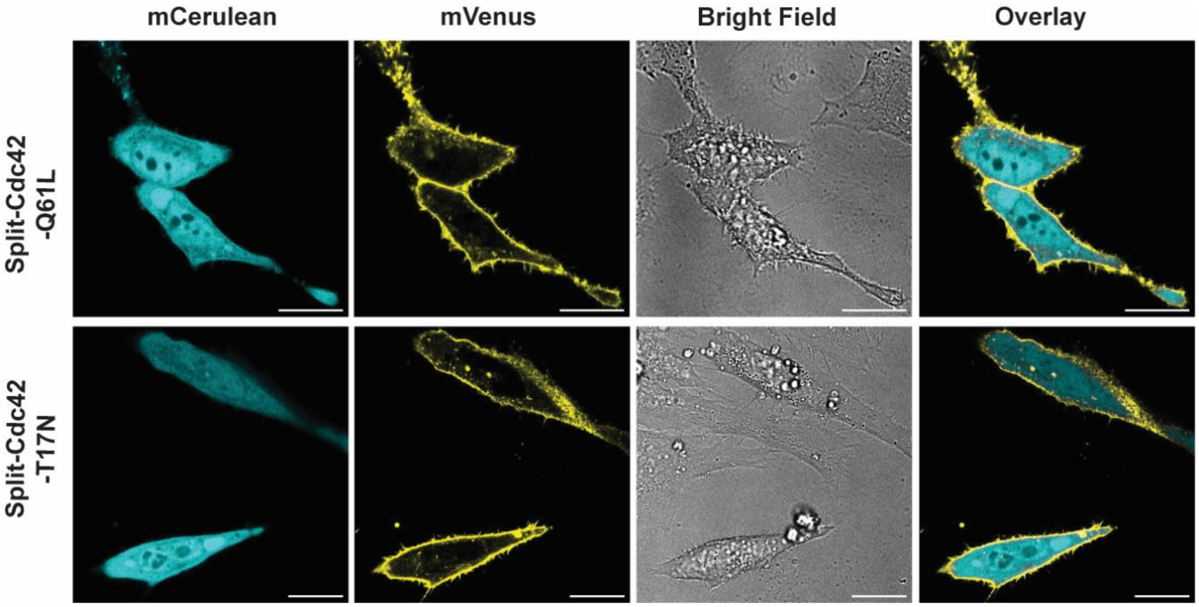

**Confocal images of HeLa cells expressing the split-Cdc42 fragments**. mCerulean fluorescence is observed in the cytosol while the mVenus fluorescence is localized to the cell membrane for split-Cdc42-Q61L (top) or split-Cdc42-T17N (bottom). Scale bar represents 20 µm.

**Fig. S3**

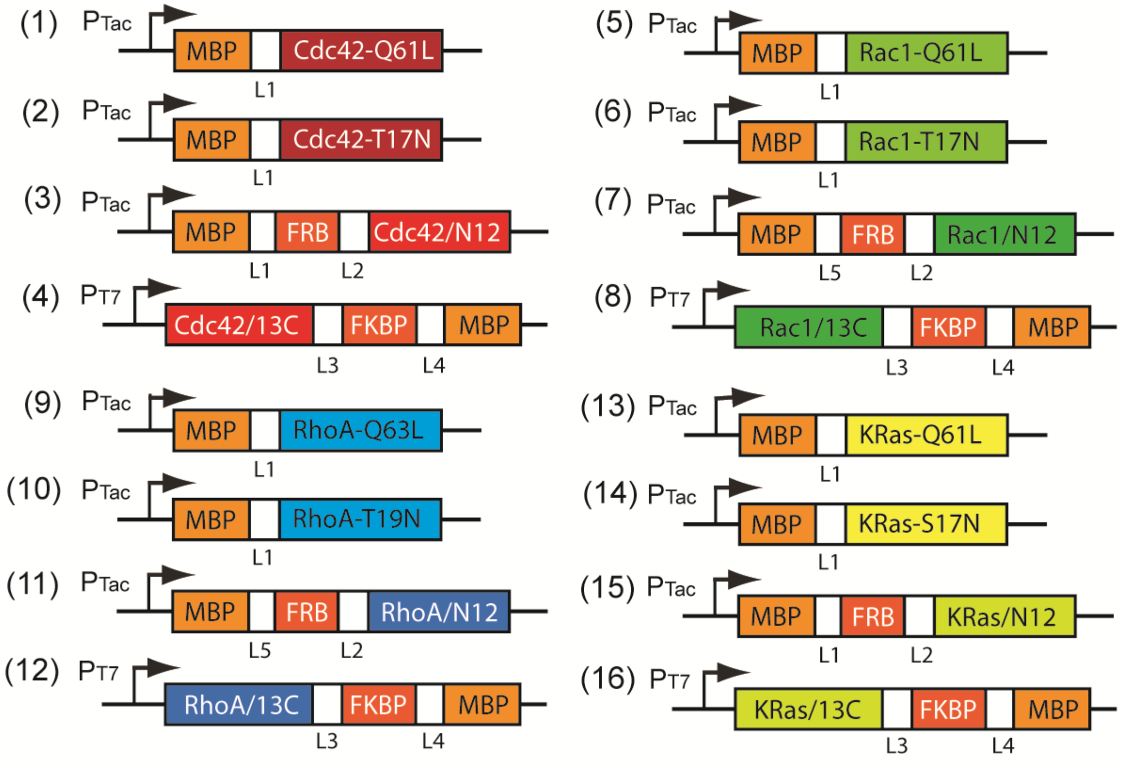

L1: NSSSNNNNNNNNNNLGIEGRISHM
L2: QISYASRGGGSSGGGEL
L3: GDPNWELVYTARLQGGGSSGGGQISYASRG
L4: KLENLYFQG
L5: NSSSNNNNNNNNNNLGIEGRISHMSMGGR

**A schematic of protein constructs used for bacterial expression**. See Table S2 for full sequence information. The GTPase/13C sequence for constructs (4), (8), (12), and (16) are based on the constitutively active mutants of corresponding small GTPases.

**Fig. S4**

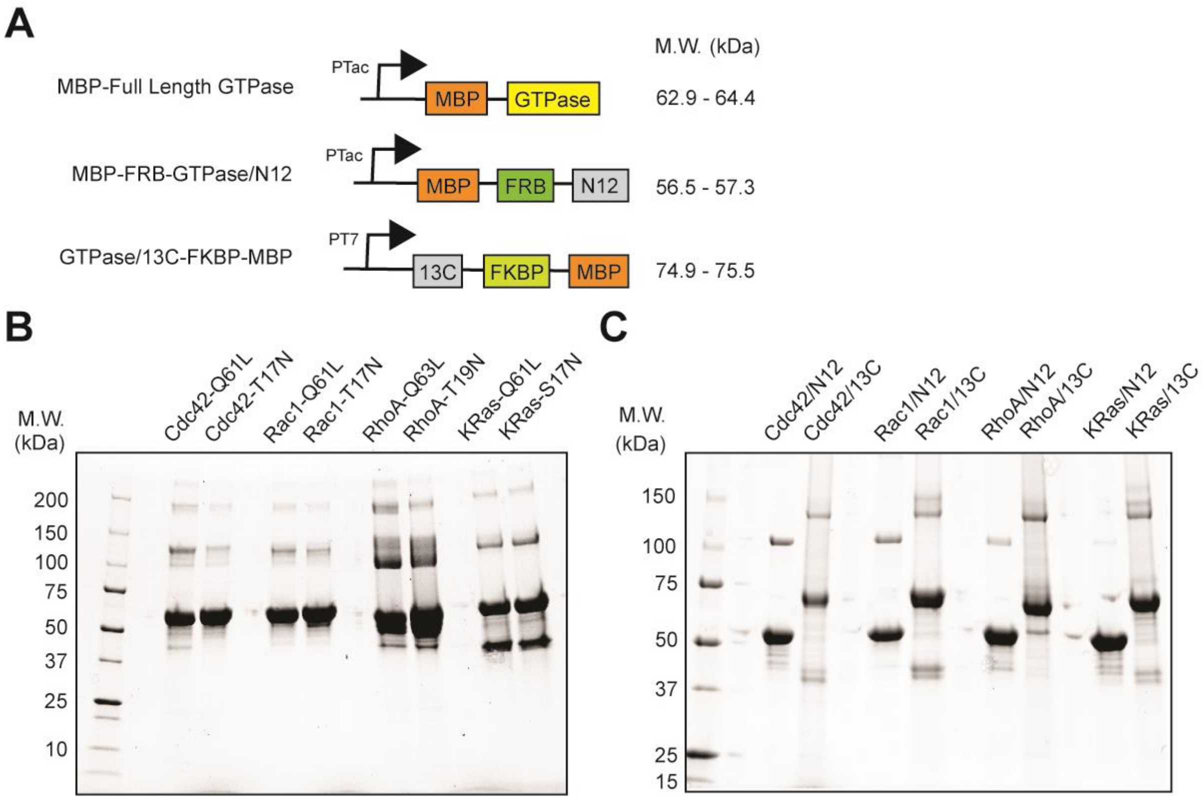

**Recombinant expression and purification of split-small GTPases.** (**A**) Constructs used for bacterial expression. Coomassie stained, SDS-PAGE gels of full-length (**B**) or fragmented (**C**) small GTPases.

**Fig. S5**

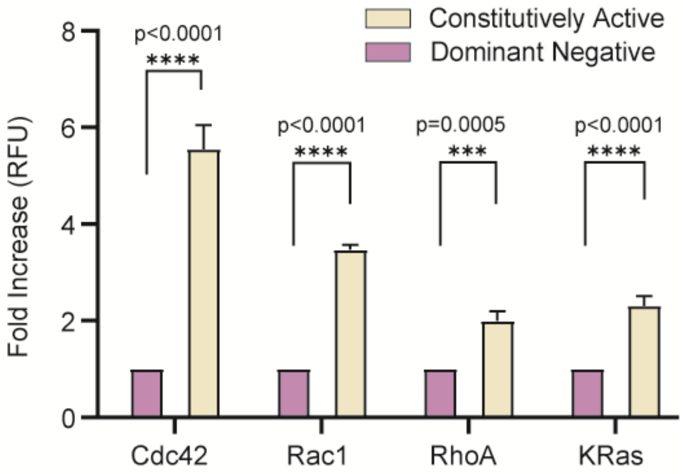

***In vitro* mantGTP assay for full-length small GTPases.** The mantGTP assay depicted in Fig. 3A can differentiate between full-length constitutively active and dominant negative versions of all small GTPases used in this work. The concentrations of full-length proteins used in the assay were 5 μM. Data represent means ± SD. *** indicates a p-value of <0.001, and **** indicates a p-value of <0.0001 calculated using a two-tailed, unpaired students' t-test.

**Fig. S6**

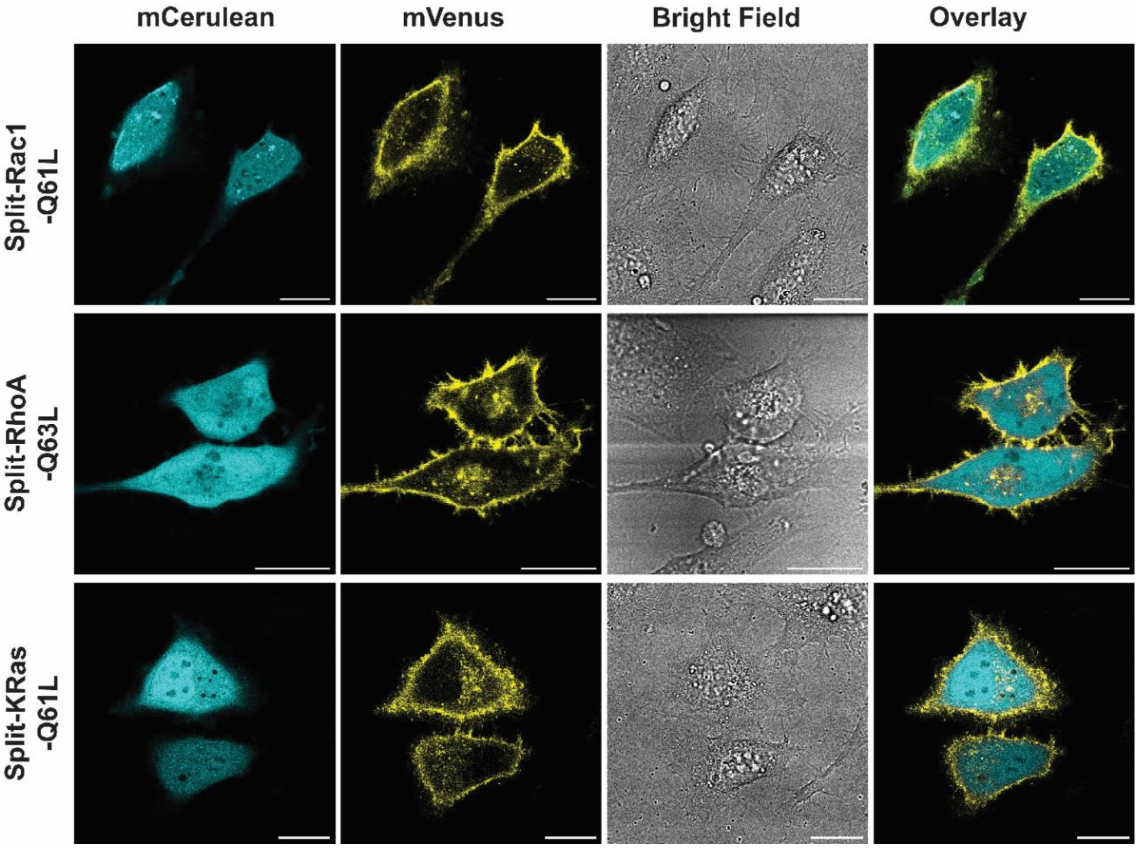

**Confocal images of HeLa cells expressing split-Rac1-Q61L, split-RhoA-Q63L, or split-KRas-Q61L fragments appended to rapamycin-dependent interaction domains.** The mCerulean fluorescence is localized to the cytosol while mVenus fluorescence is predominantly observed at the plasma membrane. Scale bar represents 20 μm.

**Fig. S7**

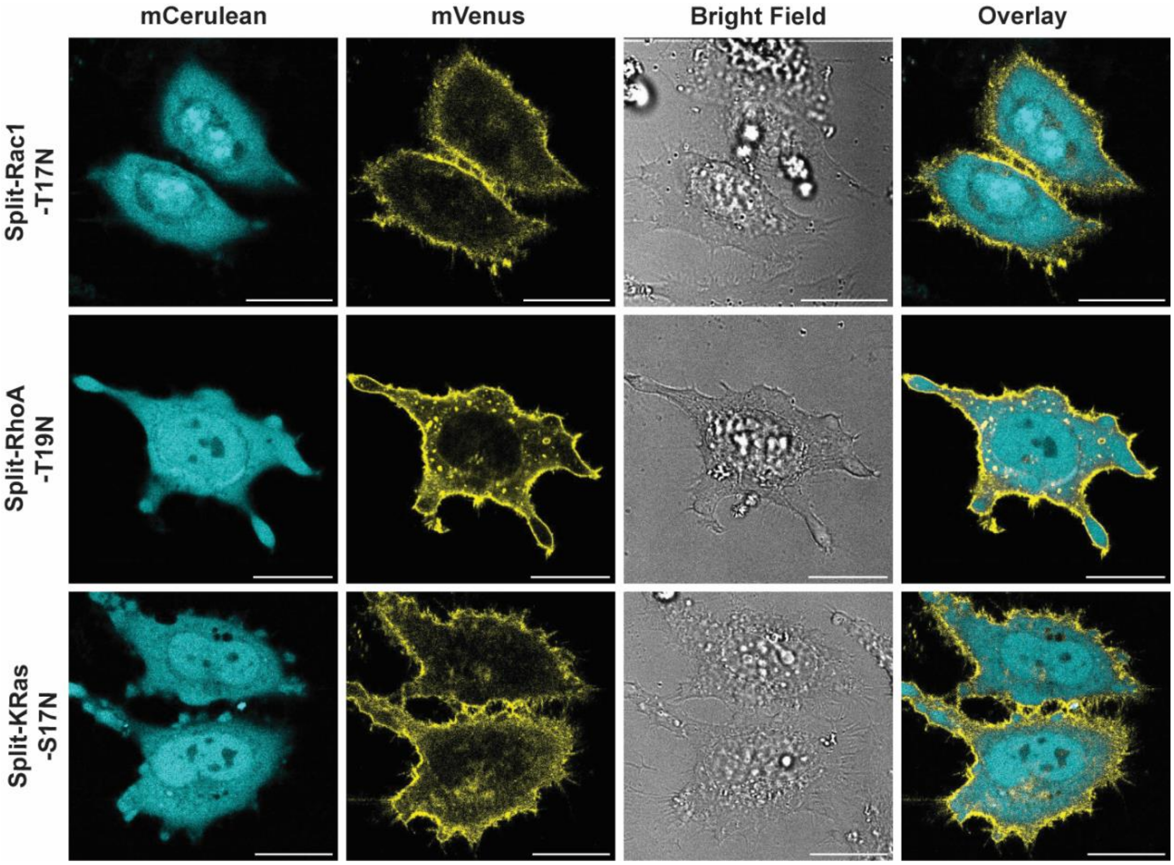

**Confocal images of HeLa cells expressing split-Rac1-T17N, split-RhoA-T19N, or split-KRas-S17N.** The mCerulean fluorescence is also localized to the cytosol while mVenus fluorescence is predominantly observed at the plasma membrane. Scale bar represents 20 μm.

**Fig. S8**

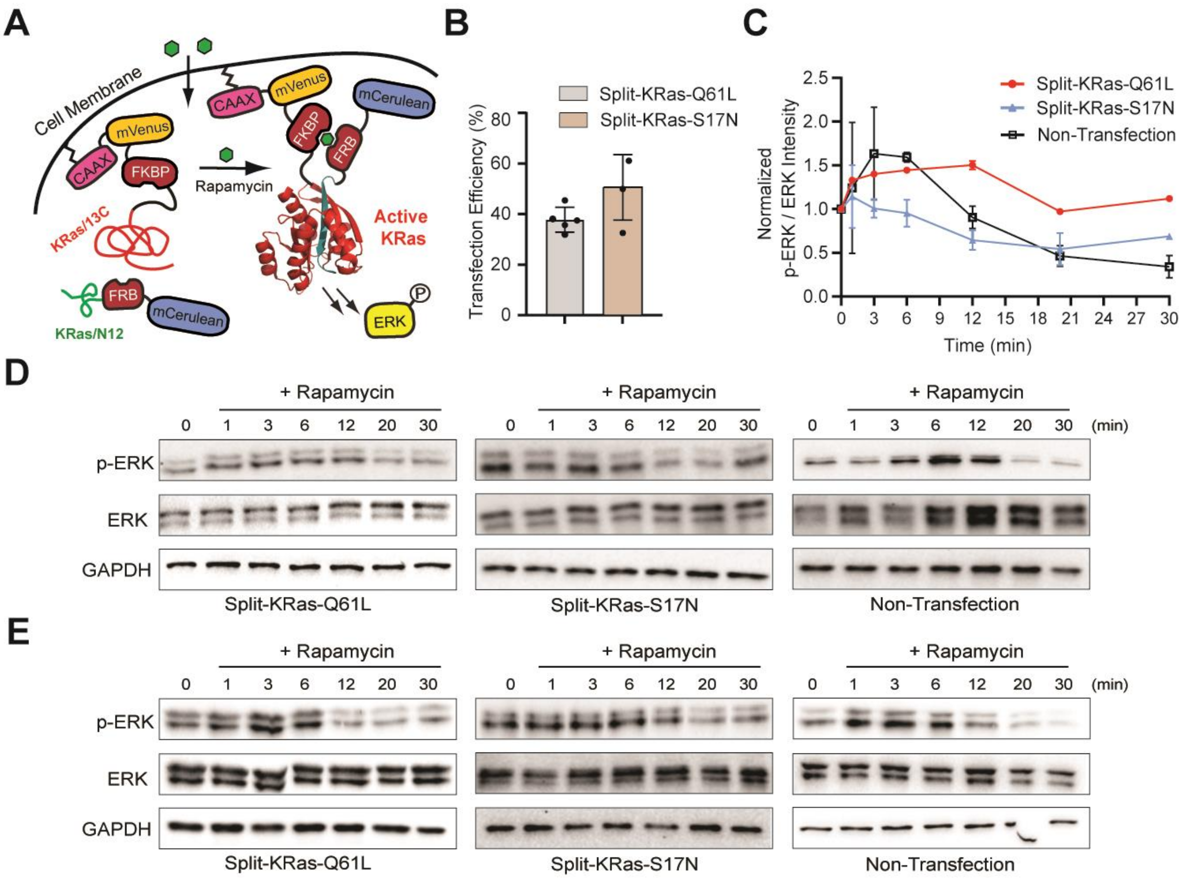

**Rapamycin gated activation of split-KRas activity in mammalian cells.** (**A**) A schematic showing split-KRas activation of ERK phosphorylation using rapamycin as a CID. (**B**) Transfection efficiency of split-KRas constructs in HeLa cells was determined by normalizing the number of fluorescent cells to the total observed cells in the bright field channel (N > 200 cells in each sample). (**C**) Quantified western blot intensities for p-ERK relative to total ERK from two biological replicates. (D) and (E) Two independent western blots from HeLa cells transfected with the indicated construct and stimulated with 500 nM rapamycin for the indicated time.

Fig. S9

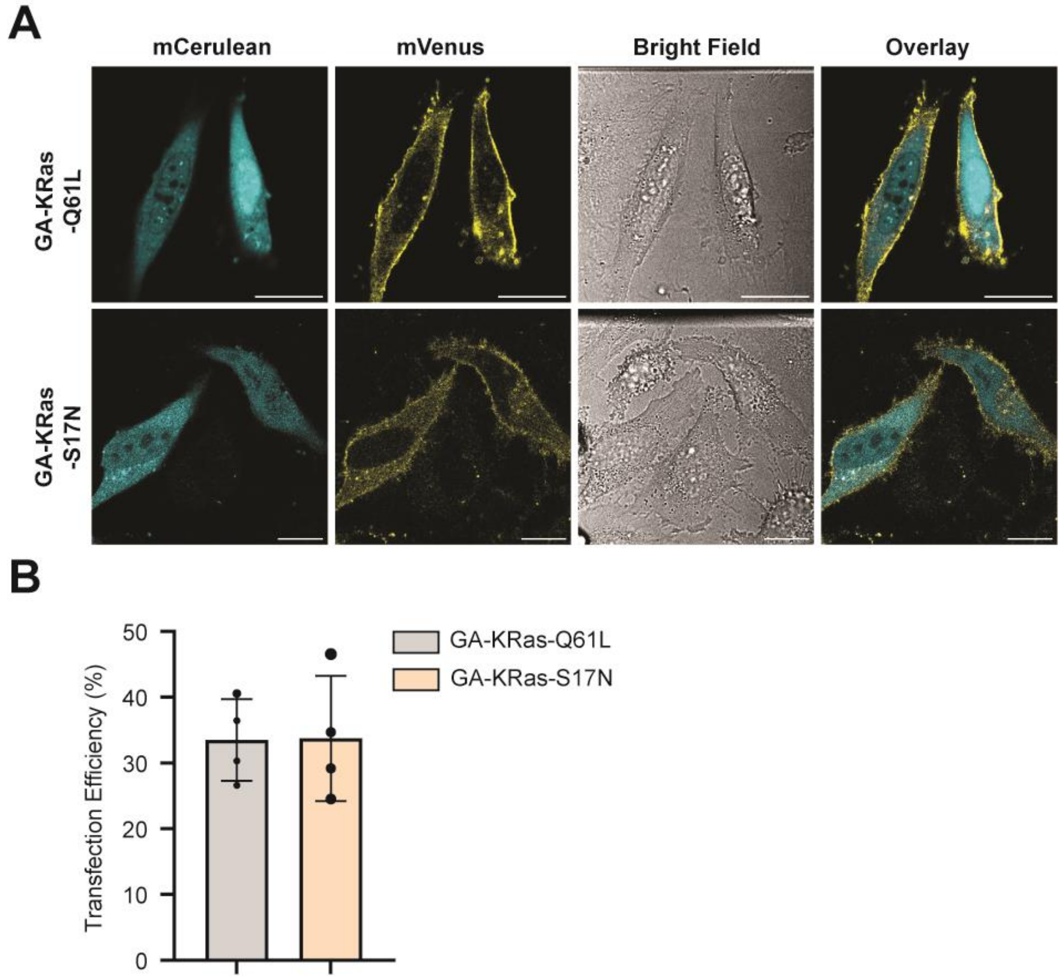

**Confocal images of HeLa cells expressing GA-gated-split-KRas constructs.** (**A**) GA-gated constructs demonstrate a mCerulean fluorescence in the cytosol and membrane-localized mVenus fluorescence. Scale bar represents 20 µm. (**B**) Transfection efficiency of GA-split-KRas constructs in HeLa cells was determined by normalizing the number of fluorescent cells to the total observed cells in the bright field channel (N > 200 cells in each sample).

**Fig. S10**

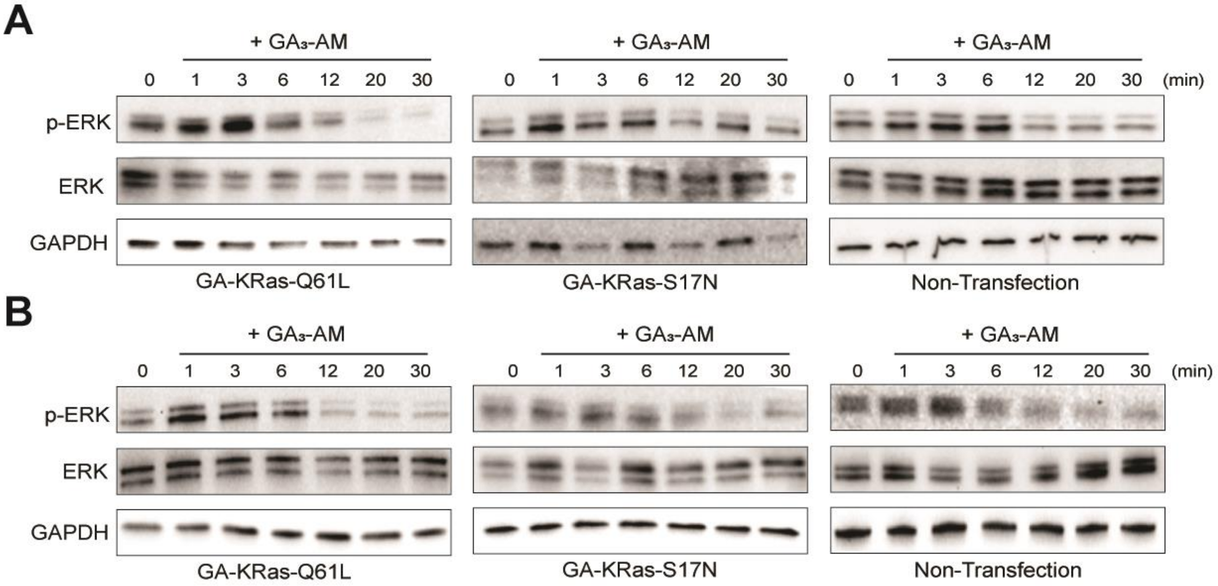

**Western blots for biological replicates of gibberellic acid-gated split-KRas induced phosphorylation of ERK. (A)** and **(B)** are from HeLa cells transfected with the indicated construct and stimulated with 10 μM gibberellic acid for the indicated time.

### General Experimental Details

*Instrumentation, and Reagents*

Polymerase chain reaction (PCR) was conducted using a thermocycler (Eppendorf, 05-414-456). DNA concentrations were quantified using a NanoDrop (Thermo Fisher, CHEM-PR1-KIT). Images from agarose gels, SDS-PAGE, and western blots were captured with a ChemiDoc XRS+ gel imager (Bio-Rad, 1708265). A Synergy H1 hybrid plate reader (Thermo Fisher, 11-120-533) was employed for mantGTP and Bradford protein assays. Counting of mammalian cells was performed using a Countess™ 3 automated cell counter (Invitrogen, AMQAX2000). Confocal images were obtained on a Leica SP5X Laser Scanning Microscope and Leica STELLARIS 8 confocal/FLIM/tauSTED microscope system equipped with tunable white light laser.

Cell culture medium, DMEM (Dulbecco's Modified Eagle Medium, Thermo Fisher, 10569010) was supplemented with 10% (v/v) fetal bovine serum,100 U/ml penicillin and 100 g/ml streptomycin. Additional reagents included Gibco™ DPBS (Thermo Fisher, 14-040-133), Giboco™ DMEM (Thermo Fisher, 21-063-029), Gibco™ Opti-MEM™ I Reduced Serum Medium (Thermo Fisher, 11-058-021), and DMSO (Sigmal-Aldrich, D8418-100ML). Enzymes and kits used for molecular biology included Platinum Taq DNA polymerase High Fidelity (Thermo Fisher, 11304011), restriction enzymes from New England Biolabs (NEB), T4 DNA ligase (NEB, M0202M), Gibson Assembly Master Mix (NEB, E2611S), and kits from Qiagen for plasmid preparation (Miniprep Kit, 27104; Maxi Kit, 12162). Site-directed mutagenesis was performed using the QuikChange II XL Site-Directed Mutagenesis Kit (Agilent Technologies, 200521). Cloning was conducted in *Escherichia coli* XL10-Gold Ultracompetent cells (Agilent Technologies, 200315), and protein expression performed in *Escherichia coli* BL21-Gold (DE3) competent cells (Agilent Technologies, 230132). Assays and protein purification used mantGTP (2'-(or-3')-O-(N-Methylanthraniloyl) Guanosine 5'-Triphosphate, Trisodium Salt) (Thermo Fisher, M12415), Zeba™ desalt spin column (Thermo Fisher, 89892), Isopropyl-β-D-thiogalactopyranoside (IPTG) (Chem Impex, 00194), B-PER™ complete bacterial protein extraction reagent (Thermo Fisher, 89821), Econo-Pac Columns (Bio-Rad, 7321010), Amylose Resin (NEB, E8021S), Amicon Ultra-15 Centrifugal Filter Unit, Ultracel, 3 KDa (EMD Millipore, UFC900324), Pierce™ Slide-A-Lyzer® Dialysis Cassettes (Thermo Fisher, 66380), Snakeskin™ Dialysis Tubing, 10K MWCO (Thermo Fisher, 68100). Cdc42 pull-down assays were performed using a commercial kit (Cytoskeleton, BK034). CIDs such as Rapamycin (Santa Cruz Biotechnology, sc-3504A) and Gibberellic Acid Acetoxymethyl Ester ($GA_3$-AM, Millipore Sigma, SML1959-50MG) were purchased from commercial sources. mantGTP assays were conducted in 96-well half-area black flat bottom polystyrene plates (Sigma-Aldrich, CLS3694-100EA). For cell lysis buffer preparation, both Protease inhibitor cocktail III (10 µl/ml, Calbiochem, 539134) and phosphatase inhibitor cocktail 1 (10 µl/ml, Sigma, P2825) were purchased commercially. The Bradford protein assay kit (Bio-Rad, 5000201) was used to determine total protein concentrations in mammalian cell lysates. Western blotting was conducted using antibodies from Cell Signaling Technologies, including Phospho-p44/42 MAPK (Erk1/2) (4377S), P44/42 MAPK (Erk1/2) (9102S), GST (2622S), GAPDH (3683S), β-actin (4967S), and HRP conjugated goat anti-rabbit IgG (7074S). Visualization of blots was achieved using Super Supersignal® West Dura Extended Duration Chemiluminescent Substrate (Thermo Fisher, 34076).

*Cloning*

Constructs used for mammalian transfection were primarily prepared using multiple overlap extension PCR steps followed by restriction enzyme-based cloning. In certain cases, were overlap extension PCR proved challenging due to repetitive linker sequences, Gibson assembly was employed (*1*). The mCerulean and mVenus sequences were derived from the mCerulean N1 (Addgene, 27795) and mVenus C1 (Addgene, 27794) plasmids, which were a gift from Steven Vogel at National Institutes of Health. The Cdc42 N12/13C fragments were obtained from previously published vectors (*2*), and were fused with FRB and FKBP. GID1 and $GAI_{1-92}$ were amplified from the pSLQ2812 pPB: CAG-GID1-VPR-IRES-Puro-WPRE PGK-GAI-tagBFP-SpdCas9 plasmid (Addgene, 84240), which was a gift from Stanley Qi at Stanford University. Several small GTPase plasmids, both constitutively active and dominant negative, were acquired from Addgene: pcDNA3-EGFP-Rac1 (Q61L) (Addgene, 13720), pcDNA3-EGFP-Rac1-T17N (Addgene, 12982), pcDNA3-EGFP-RhoA-Q63L (Addgene, 12968), pcDNA3-EGFP-RhoA-T19N (Addgene, 12967), Hs.KRAS4b Q61L (Addgene, 83134), and Hs.KRAS4b S17N (Addgene, 83156). The above-mentioned plasmids were a gift from Klaus Hahn at University of North Carolina Chapel Hill, Gary Bokoch at Scripps Research Institute, and Dominic Esposito at Frederick National Laboratory. The authors wish to thank the above-mentioned PIs for generously sharing these resources. In brief, the mCerulean sequence was amplified via PCR using a reverse primer that overlapped with the linker region of the forward primer used for amplification of the FRB-Cdc42/N12 sequence. The resulting PCR products were assembled by overlap extension PCR, digested with NheI and EcoRI, and cloned into MCS A of the pIRES vector (Takara Bio, 631605). Analogously, a Cdc42/13C-FKBP-mVenus sequence was obtained using overlap extension PCR, incorporating a CAAX motif at its 3'-end. The resulting sequence was digested with XbaI and NotI and ligated into MCS B of the pIRES vector. Cloning steps were analogous for split-Rac1, split-RhoA, split-KRas, and gibberellic acid-gated constructs. In cases where overlap extension PCR was unsuccessful, Gibson assembly was employed.

Plasmids designed for recombinant protein expression were built using pMAL-c5X-His (Genophore, GV03006) or a custom pET21b plasmid containing a C-terminal MBP purification tag. All plasmids were confirmed by DNA sequencing. Protein constructs are shown in **Figure S1 and S3,** and corresponding sequences are provided in **Table S2 and S3**.

*Mammalian cell culture and plasmid transfection*

HeLa cells (ATCC, CCL-2) and Chinese hamster ovary cells (ATCC, CCL-61) were used in this work. Cells were cultured in Dulbecco's modified Eagle's medium (DMEM) supplemented with 10% (v/v) fetal bovine serum and 100 U/ml penicillin and 100 g/ml streptomycin, maintained at 37°C in a 5% $CO_2$ humidified chamber. For plasmid transfection, cells were seeded at 1.2 x $10^5$ cells/$cm^2$ either on tissue culture treated dishes (Thomas Scientific, 1228K66) for pull-down assays and western blots, or on 35 mm poly-d-lysine coated glass-bottom dishes (MatTek Corporation, NC9005934) for confocal microscopy. After allowing 24 hr for growth, resulting in approximately 70-80% confluency, transient transfection was conducted according to the manufacturer's instructions (Invitrogen, L3000015).

*PAK-PBD pull-down assays*

After transfection, CHO cells were serum-starved in Opti-MEM and cultivated for 36 hr. Following this, cells underwent a 5 min wash with pre-warmed DPBS, then a 20 min incubation with either 2 ml DPBS or 2 ml DPBS containing 1 µM rapamycin. The media were then removed gently. Subsequently, the cells were lysed using a cell lysis buffer (Cytoskeleton, BK034) supplemented with a 1X protease inhibitor cocktail (Cytoskeleton, PIC02). After 5 min incubation on ice, lysates were centrifuged at 10,000 g at 4°C for 1 min. The Bradford protein assay kit (Bio-Rad, 5000201), using BSA as a reference, was used to determine total protein concentrations in cell lysates, which were then normalized for subsequent experiments. Equal aliquots of cell lysates (ranging from 300-800 µg) were combined with 10 µg GST-tagged PAK-PBD protein agarose beads (Cytoskeleton, PAK02) and rotated at 4°C for 1 hr. The beads were then centrifuged at 5000 g for a minute at 4°C. Supernatants were carefully pipetted out, and the beads were washed with 500 µl wash buffer (Cytoskeleton, BK034), followed by another round of centrifugation at 5000 g for 3 min at 4°C. After supernatant removal, 30 µl of wash buffer and 6 µl of SDS-loading gel buffer were introduced into the centrifuge tubes. These tubes were then heated at 80°C for 10 min. Equal volumes of cell lysates, both pre and post pull-down, were separated on a 12% SDS-PAGE and transferred to a nitrocellulose membrane overnight at 4 °C. Blots were probed with anti-Venus (YFP) antibody (Millipore Sigma, MABE1906-100UL) for Cdc42/13C-FKBP-mVenus detection. To verify consistent loading of protein samples, a GST antibody (Cell Signaling, 2622S) and GAPDH antibody (Cell Signaling, 3683S) were utilized.

*Confocal fluorescence imaging*

Confocal images were acquired on a Leica SP5X Laser Scanning Microscpe and Leica STELLARIS 8 confocal/FLIM/tauSTED microscope system equipped with tunable white light lasers and a 37-2 digital temperature controller. Image acquisition was facilitated by the LAS-AF software, and subsequent processing was done in Fiji (ImageJ). To visualize the nucleus and cytoskeletal structures, cells were pre-stained with Hoechst 33342 (Thermo Fisher, H3570) and CellMask$^{TM}$ Deep Red Actin Tracking Stain (Thermo Fisher, A57245) as per the manufacturer's guidelines. After staining, cells were rinsed three times with pre-warmed DPBS and then incubated in 1.5 ml DMEM containing 25 mM HEPES (Thermo Fisher, 21-063-029) for imaging purposes. For image acquisition following stimulation with the CID, 0.5 ml of DMEM containing either 4X rapamycin or 4X $GA_3$-AM stock solutions (relative to the final concentration) were introduced into the glass-bottom dishes via a syringe drop-wise. Fluorophores were excited using a diode laser (405 nm) or a white light laser (440-790 nm). Nucleus staining was observed with excitation at 405 nm, using a PMT detector set at 430-460 nm for emission. Actin staining was observed using excitation at 652 nm, and emission was detected at 660-710 nm. The mCerulean fluorescence was observed using 458 nm excitation and collecting emission at 468-508 nm, while the mVenus fluorescence was monitored using 514 nm excitation and emission was detected at 520-560 nm.

*Recombinant protein expression and purification*

Plasmids encoding MBP-fused GTPases were transformed into *Escherichia coli* BL21-Gold (DE3) strain. These transformed cells were then inoculated into 1L of terrific broth (TB) and shaken at 37°C until an $OD_{600}$ of 0.8 was reached. Protein expression was induced by adding 0.5 mM IPTG and incubating the culture at 37°C for 3 hr with 250 rpm shaking. This was followed by an extended incubation at 18°C for 18 hr. Cells were collected through centrifugation at 3220 g for 35 min at 4°C. The resulting cell pellets were lysed using the B-PER™ complete bacterial protein extraction reagent (Thermo Fisher, 89821) and clarified by centrifugation at 15,000 g for 20 min at 4°C. The supernatant was then diluted with six volumes of 1X column buffer (20 mM Tris-Cl, pH 7.4 at 25°C; 200 mM NaCl; 1 mM EDTA) and purified using amylose resin (NEB, E8021S) using gravity flow (0.5 mL resin per 1L bacterial culture). The column containing the amylose resin and diluted protein solution was washed with 100 mL 1X column buffer. Elution of the recombinant proteins was achieved using 10 mM maltose in 10 mL column buffer. Eluted proteins were subsequently concentrated using an ultra-15 centrifugal filter unit (EMD Millipore, UFC900324) and dialyzed into GTPase storage buffer (50 mM Tris-Cl, pH 7.5 at 25°C; 150 mM NaCl; 20% glycerol). 13C GTPase fragments were purified from inclusion bodies. Specifically, the inclusion bodies obtained after lysis by B-PER reagent for each 13C fragment were resuspended in 1X column buffer containing 8 M urea. Solubilized protein was then loaded into Snakeskin dialysis tubing and refolded using stepwise dialysis, gradually reducing the urea concentration (4 M, 2 M, 1 M, 0.5 M). These refolded proteins underwent another purification step using 0.5 mL amylose resin (from inclusion bodies obtained per 1L bacterial culture) washed with 100 mL 1X column buffer, eluted with 10 mM maltose in 10 mL 1X column buffer, concentrated via a centrifugal filter unit, and were eventually dialyzed into the GTPase storage buffer. Protein concentrations were determined using NanoDrop, based on the calculated extinction coefficient of each protein (Table S4). The molecular weight and extinction coefficient of recombinant proteins were calculated using Expasy ProtParam (https://web.expasy.org/protparam/).

*In-vitro split-GTPase reassembly assay*

Recombinant proteins were co-incubated with a 10-fold excess of mantGTP (Thermo Fisher, M12415) in a nucleotide exchange buffer (20 mM Tris-Cl, pH 7.5 at 25°C; 50 mM NaCl, 5 mM EDTA, 1% glycerol). For full-length GTPase controls, both constitutively active and dominant negative mutants were used at 5 µM, with 50 µM mantGTP for nucleotide exchange. For split-GTPase proteins, equimolar amounts of N12 and 13C fragments of each small GTPase, ranging between 5-30 µM, were combined with equimolar rapamycin (5-30 µM) and a 10-fold excess of mantGTP (50-300 µM). The nucleotide exchange process was carried out at room temperature, with shaking at 300 rpm for 30 minutes. The nucleotide exchange reaction was quenched by adding excess $MgCl_2$ (1.33 M) and placing the reaction on ice. The reaction mixture was then loaded onto a Zeba™ desalting spin column (Thermo Fisher, 89892) pre-rinsed twice with a wash buffer (20 mM Tris-Cl, pH 7.5 at 25°C; 50 mM NaCl; 10 mM $MgCl_2$; 1% glycerol) and proteins were eluted at 1000 g for 2 minutes at 4°C. The elution was transferred to a black 96-well plate, and fluorescence was recorded using 355 nm excitation and 448 nm emission over an hour using a plate reader. Fold fluorescence increase after ten minutes incubation of the plate at 30°C were used and assays were normalized to dominant negative samples within each group. Solutions containing only mantGTP were used for background subtraction. Standard deviations were obtained using error propagation.

*Lysate preparation and western blot analysis*

Transiently transfected HeLa cells were serum starved overnight in Opti-MEM before stimulation with rapamycin or $GA_3$-AM. Cells were exposed to either 500 nM rapamycin or 10 μM $GA_3$-AM in pre-warmed Opti-MEM for the indicated time, followed by rinsing with ice-cold DPBS. Subsequently, cells were lysed using the following lysis buffer: 50 mM Tris-Cl (pH = 7.5 at 25°C), 150 mM NaCl, 50 mM β-glycerophosphate, 10 mM sodium pyrophosphate, 30 mM NaF, 1% Triton X-100, 2 mM EGTA, 100 μM $Na_3VO_4$, 1 mM DTT, protease inhibitor cocktail III (10 μl/ml, Calbiochem, 539134), and phosphatase inhibitor cocktail 1 (10 μl/ml, Sigma, P2825). The lysates were placed on ice for 15 min followed by centrifugation at 17,000 g for 10 min at 4°C. A Bradford assay (Bio-Rad, 5000201) was performed to determine total protein concentrations, which were then normalized for downstream experiments. For western blotting, cell lysates (5-20 μg total protein) were separated using 12% SDS-PAGE gels and transferred onto nitrocellulose membranes. These membranes were blocked and probed with the indicated primary antibodies, and detection was performed through enhanced chemiluminescence (Thermo Scientific, 34076).

*Data Processing and Statistical analyses*

For the detection and quantification of Cdc42-induced filopodia formation, we utilized the open-source ImageJ plugin, FiloQuant, adhering to its comprehensive instructions (*3*). This software facilitates the extraction of measurable data, including the number and length of filopodia and the cell edge length (*4*). We assessed the number of detected filopodia per cell to gauge the influence of our split-Cdc42 system on filopodia formation. In each experiment, data from over 50 transfected cells were aggregated and quantified.

For evaluation of split-Rac1 signaling, two independent observers counted the number of transfected cells (>50), and evaluated the percentage of these transfected cells exhibiting evident membrane ruffling. Their collective outcomes were then normalized for inter-rater reliability. Each single transfected cell is defined as a variable, and the observers scored the same variable. These results were used to calculate a Cohen's Kappa value, which quantifies the agreement of raters (*5*). The Cohen's Kappa value for each sample ranged from 0.66 to 1.0, with a median of 0.82, exceeding the commonly accepted 0.61 - 0.80 threshold for substantial agreement, indicating almost perfect agreement.

For assessment of split-RhoA signaling, the cell area was determined using ImageJ (ROI, of the same transfected cell pre- and post-rapamycin stimulation). A decline exceeding 20% in a transfected cell's area was regarded as a significant impact triggered by RhoA activity.

Statistical analyses were conducted using unpaired, two-tailed *t*-tests (GraphPad Prism 9.5, GraphPad Software).

**Table S1. Construct description and Addgene IDs**.

| Construct | Description | Addgene ID |
|---|---|---|
| **Mammalian Cell** | | |
| Split-Cdc42-Q61L | pIRES-mCerulean-FRB-Cdc42/N12-Cdc42/13C(Q61L)-FKBP-mVenus-CAAX | 214278 |
| Split-Cdc42-T17N | pIRES-mCerulean-FRB-Cdc42/N12-Cdc42/13C(T17N)-FKBP-mVenus-CAAX | 214279 |
| Split-Rac1-Q61L | pIRES-mCerulean-FRB-Rac1/N12-Rac1/13C(Q61L)-FKBP-mVenus-CAAX | 214280 |
| Split-Rac1-T17N | pIRES-mCerulean-FRB-Rac1/N12-Rac1/13C(T17N)-FKBP-mVenus-CAAX | 214281 |
| Split-RhoA-Q63L | pIRES-mCerulean-FRB-RhoA/N12-RhoA/13C(Q63L)-FKBP-mVenus-CAAX | 214282 |
| Split-RhoA-T19N | pIRES-mCerulean-FRB-RhoA/N12-RhoA/13C(T19N)-FKBP-mVenus-CAAX | 214283 |
| Split-KRas-Q61L | pIRES-mCerulean-FRB-KRas/N12-KRas/13C(Q61L)-FKBP-mVenus-CAAX | 214284 |
| Split-KRas-S17N | pIRES-mCerulean-FRB-KRas/N12-KRas/13C(S17N)-FKBP-mVenus-CAAX | 214285 |
| GA-Split-KRas-Q61L | pIRES-mCerulean-FRB-KRas/N12-KRas/13C(Q61L)-GAI1-92-mVenus-CAAX | 214286 |
| GA-Split-KRas-S17N | pIRES-mCerulean-FRB-KRas/N12-KRas/13C(S17N)-GAI1-92-mVenus-CAAX | 214287 |
| ***E. Coli* Expression** | | |
| Full length Cdc42-Q61L | pMAL-MBP-Cdc42-Q61L | 214288 |
| Full length Cdc42-T17N | pMAL-MBP-Cdc42-T17N | 214289 |
| N12 fragment Cdc42 | pMAL-MBP-FRB-Cdc42/N12 | 214290 |
| 13C fragment Cdc42 | pET21b-Cdc42/13C-FKBP-MBP | 214291 |
| Full length Rac1-Q61L | pMAL-MBP-Rac1-Q61L | 214292 |
| Full length Rac1-T17N | pMAL-MBP-Rac1-T17N | 214293 |
| N12 fragment Rac1 | pMAL-MBP-FRB-Rac1/N12 | 214294 |
| 13C fragment Rac1 | pET21b-Rac1/13C-FKBP-MBP | 214295 |
| Full length RhoA-Q63L | pMAL-MBP-RhoA-Q63L | 214296 |
| Full length RhoA-T19N | pMAL-MBP-RhoA-T19N | 214297 |
| N12 fragment RhoA | pMAL-MBP-FRB-RhoA/N12 | 214298 |
| 13C fragment RhoA | pET21b-RhoA/13C-FKBP-MBP | 214299 |
| Full length KRas-Q61L | pMAL-MBP-KRas-Q61L | 214300 |
| Full length KRas-S17N | pMAL-MBP-KRas-S17N | 214301 |
| N12 fragment KRas | pMAL-MBP-FRB-KRas/N12 | 214302 |
| 13C fragment KRas | pET21b-KRas/13C-FKBP-MBP | 214303 |

**Table S2. Sequences for mammalian expression constructs used in this study.**

1）**pIRES-mCerulean-FRB-Cdc42/N12-Cdc42/13C(Q61L)-FKBP-mVenus-CAAX**

>Amino acid sequence (61L is underlined)

MVSKGEELFTGVVPILVELDGDVNGHKFSVSGEGEGDATYGKLTLKFICTTGKLPVPWPTLVTT
LTWGVQCFARYPDHMKQHDFFKSAMPEGYVQERTIFFKDDGNYKTRAEVKFEGDTLVNRIELK
GIDFKEDGNILGHKLEYNAISDNVYITADKQKNGIKANFKIRHNIEDGSVQLADHYQQNTPIGDGP
VLLPDNHYLSTQSKLSKDPNEKRDHMVLLEFVTAAGITLGMDELYKGGGSSGGGEMWHEGLE
EASRLYFGERNVKGMFEVLEPLHAMMERGPQTLKETSFNQAYGRDLMEAQEWCRKYMKSGN
VKDLTQAWDLYYHVFRRISKQQISYASRGGGSSGGGELQTIKCVVVGDGA*---***IRES Region***---
MGVGKTCLLISYTTNKFPSEYVPTVFDNYAVTVMIGGEPYTLGLFDTAGLEDYDRLRPLSYPQT
DVFLVCFSVVSPSSFENVKEKWVPEITHHCPKTPFLLVGTQIDLRDDPSTIEKLAKNKQKPITPET
AEKLARDLKAVKYVECSALTQRGLKNVFDEAILAALEPPETQPGDPNWELVYTARLQGGGSSG
GGQISYASRGGVQVETISPGDGRTFPKRGQTCVVHYTGMLEDGKKFDSSRDRNKPFKFMLGK
QEVIRGWEEGVAQMSVGQRAKLTISPDYAYGATGHPGIIPPHATLVFDVELLKLEGGGSSGGGV
SKGEELFTGVVPILVELDGDVNGHKFSVSGEGEGDATYGKLTLKLICTTGKLPVPWPTLVTTLGY
GLQCFARYPDHMKQHDFFKSAMPEGYVQERTIFFKDDGNYKTRAEVKFEGDTLVNRIELKGIDF
KEDGNILGHKLEYNYNSHNVYITADKQKNGIKANFKIRHNIEDGGVQLADHYQQNTPIGDGPVLL
PDNHYLSYQSKLSKDPNEKRDHMVLLEFVTAAGITLGMDELYKKKKKKKSKTKCVIM*

>DNA sequence

ATGGTGAGCAAGGGCGAGGAGCTGTTCACCGGGGTGGTGCCCATCCTGGTCGAGCTGGA
CGGCGACGTAAACGGCCACAAGTTCAGCGTGTCCGGCGAGGGCGAGGGCGATGCCACCT
ACGGCAAGCTGACCCTGAAGTTCATCTGCACCACCGGCAAGCTGCCCGTGCCCTGGCCCA
CCCTCGTGACCACCCTGACCTGGGGCGTGCAGTGCTTCGCCCGCTACCCCGACCACATGA
AGCAGCACGACTTCTTCAAGTCCGCCATGCCCGAAGGCTACGTCCAGGAGCGCACCATCT
TCTTCAAGGACGACGGCAACTACAAGACCCGCGCCGAGGTGAAGTTCGAGGGCGACACC
CTGGTGAACCGCATCGAGCTGAAGGGCATCGACTTCAAGGAGGACGGCAACATCCTGGGG
CACAAGCTGGAGTACAACGCCATCAGCGACAACGTCTATATCACCGCCGACAAGCAGAAGA
ACGGCATCAAGGCCAACTTCAAGATCCGCCACAACATCGAGGACGGCAGCGTGCAGCTCG
CCGACCACTACCAGCAGAACACCCCCATCGGCGACGGCCCCGTGCTGCTGCCCGACAAC
CACTACCTGAGCACCCAGTCCAAGCTGAGCAAAGACCCCAACGAGAAGCGCGATCACATG
GTCCTGCTGGAGTTCGTGACCGCCGCCGGGATCACTCTCGGCATGGACGAGCTGTACAAG
GGCGGTGGCTCATCTGGCGGAGGTGAGATGTGGCATGAAGGCCTGGAAGAGGCATCTCGT
TTGTACTTTGGGGAAAGGAACGTGAAAGGCATGTTTGAGGTGCTGGAGCCCTTGCATGCTA
TGATGGAACGGGGCCCCCAGACTCTGAAGGAAACATCCTTTAATCAGGCCTATGGTCGAGA
TTTAATGGAGGCCCAAGAGTGGTGCAGGAAGTACATGAAATCAGGGAATGTCAAGGACCTC
ACCCAAGCCTGGGACCTCTATTATCATGTGTTCCGACGAATCTCAAAGCAGCAGATCTCGTA
CGCGTCCCGGGGCGGTGGCTCATCTGGCGGAGGTGAGCTCCAGACAATTAAGTGTGTTGT
TGTGGGCGATGGTGCTTAA---***IRES region***---
ATGGGGGTTGGTAAAACATGTCTCCTGATATCCTACACAACAAACAAATTTCCATCGGAGTAT
GTACCGACTGTTTTTGACAACTATGCAGTCACAGTTATGATTGGTGGAGAACCATATACTCTT
GGACTTTTTGATACTGCAGGGCTAGAGGATTATGACAGATTACGACCGCTGAGTTATCCACA
AACAGATGTATTTCTAGTCTGTTTTTCAGTGGTCTCTCCATCTTCATTTGAAAACGTGAAAGA
AAAGTGGGTGCCTGAGATAACTCACCACTGTCCAAAGACTCCTTTCTTGCTTGTTGGGACT

CAAATTGATCTCAGAGATGACCCCTCTACTATTGAGAAACTTGCCAAGAACAAACAGAAGCC
TATCACTCCAGAGACTGCTGAAAAGCTGGCCCGTGACCTGAAGGCTGTCAAGTATGTGGAG
TGTTCTGCACTTACACAGAGAGGTCTGAAGAATGTGTTTGATGAGGCTATCCTAGCTGCCCT
CGAGCCTCCGGAAACTCAACCCGGGGATCCCAATTGGGAGCTCGTGTACACGGCGCGCCT
GCAGGGAGGTGGCTCATCTGGCGGAGGTCAGATCTCGTACGCGTCCCGGGGCGGAGTGC
AGGTGGAAACCATCTCCCCAGGAGACGGGCGCACCTTCCCCAAGCGCGGCCAGACCTGC
GTGGTGCACTACACCGGGATGCTTGAAGATGGAAAGAAATTTGATTCCTCCCGGGACAGAA
ACAAGCCCTTTAAGTTTATGCTAGGCAAGCAGGAGGTGATCCGAGGCTGGGAAGAAGGGG
TTGCCCAGATGAGTGTGGGTCAGAGAGCCAAACTGACTATATCTCCAGATTATGCCTATGGT
GCCACTGGGCACCCAGGCATCATCCCACCACATGCCACTCTCGTCTTCGATGTGGAGCTTC
TAAAACTGGAAGGCGGTGGCTCATCTGGCGGAGGTGTGAGCAAGGGCGAGGAGCTGTTC
ACCGGGGTGGTGCCCATCCTGGTCGAGCTGGACGGCGACGTAAACGGCCACAAGTTCAG
CGTGTCCGGCGAGGGCGAGGGCGATGCCACCTACGGCAAGCTGACCCTGAAGCTCATCT
GCACCACCGGCAAGCTGCCCGTGCCCTGGCCCACCCTCGTGACCACCCTCGGCTACGGC
CTGCAGTGCTTCGCCCGCTACCCCGACCACATGAAGCAGCACGACTTCTTCAAGTCCGCC
ATGCCCGAAGGCTACGTCCAGGAGCGCACCATCTTCTTCAAGGACGACGGCAACTACAAG
ACCCGCGCCGAGGTGAAGTTCGAGGGCGACACCCTGGTGAACCGCATCGAGCTGAAGGG
CATCGACTTCAAGGAGGACGGCAACATCCTGGGGCACAAGCTGGAGTACAACTACAACAG
CCACAACGTCTATATCACCGCCGACAAGCAGAAGAACGGCATCAAGGCCAACTTCAAGATC
CGCCACAACATCGAGGACGGCGGCGTGCAGCTCGCCGACCACTACCAGCAGAACACCCC
CATCGGCGACGGCCCCGTGCTGCTGCCCGACAACCACTACCTGAGCTACCAGTCCAAGCT
GAGCAAAGACCCCAACGAGAAGCGCGATCACATGGTCCTGCTGGAGTTCGTGACCGCCGC
CGGGATCACTCTCGGCATGGACGAGCTGTACAAGAAAAAAAAGAAGAAAAAGAGCAAGAC
CAAATGCGTGATTATGTAA

2）**pIRES-mCerulean-FRB-Cdc42/N12-Cdc42/13C(T17N)-FKBP-mVenus-CAAX**

>Amino acid sequence (17N is underlined)

MVSKGEELFTGVVPILVELDGDVNGHKFSVSGEGEGDATYGKLTLKFICTTGKLPVPWPTLVTT
LTWGVQCFARYPDHMKQHDFFKSAMPEGYVQERTIFFKDDGNYKTRAEVKFEGDTLVNRIELK
GIDFKEDGNILGHKLEYNAISDNVYITADKQKNGIKANFKIRHNIEDGSVQLADHYQQNTPIGDGP
VLLPDNHYLSTQSKLSKDPNEKRDHMVLLEFVTAAGITLGMDELYKGGGSSGGGEMWHEGLE
EASRLYFGERNVKGMFEVLEPLHAMMERGPQTLKETSFNQAYGRDLMEAQEWCRKYMKSGN
VKDLTQAWDLYYHVFRRISKQQISYASRGGGSSGGGELQTIKCVVVGDGA*---***IRES Region***---
MGVGK**N**CLLISYTTNKFPSEYVPTVFDNYAVTVMIGGEPYTLGLFDTAGQEDYDRLRPLSYPQT
DVFLVCFSVVSPSSFENVKEKWVPEITHHCPKTPFLLVGTQIDLRDDPSTIEKLAKNKQKPITPET
AEKLARDLKAVKYVECSALTQRGLKNVFDEAILAALEPPETQPGDPNWELVYTARLQGGGSSG
GGQISYASRGGVQVETISPGDGRTFPKRGQTCVVHYTGMLEDGKKFDSSRDRNKPFKFMLGK
QEVIRGWEEGVAQMSVGQRAKLTISPDYAYGATGHPGIIPPHATLVFDVELLKLEGGGSSGGGV
SKGEELFTGVVPILVELDGDVNGHKFSVSGEGEGDATYGKLTLKLICTTGKLPVPWPTLVTTLGY
GLQCFARYPDHMKQHDFFKSAMPEGYVQERTIFFKDDGNYKTRAEVKFEGDTLVNRIELKGIDF
KEDGNILGHKLEYNYNSHNVYITADKQKNGIKANFKIRHNIEDGGVQLADHYQQNTPIGDGPVLL
PDNHYLSYQSKLSKDPNEKRDHMVLLEFVTAAGITLGMDELYKKKKKKKSKTKCVIM*

>DNA sequence

ATGGTGAGCAAGGGCGAGGAGCTGTTCACCGGGGTGGTGCCCATCCTGGTCGAGCTGGA
CGGCGACGTAAACGGCCACAAGTTCAGCGTGTCCGGCGAGGGCGAGGGCGATGCCACCT
ACGGCAAGCTGACCCTGAAGTTCATCTGCACCACCGGCAAGCTGCCCGTGCCCTGGCCCA
CCCTCGTGACCACCCTGACCTGGGGCGTGCAGTGCTTCGCCCGCTACCCCGACCACATGA
AGCAGCACGACTTCTTCAAGTCCGCCATGCCCGAAGGCTACGTCCAGGAGCGCACCATCT
TCTTCAAGGACGACGGCAACTACAAGACCCGCGCCGAGGTGAAGTTCGAGGGCGACACC
CTGGTGAACCGCATCGAGCTGAAGGGCATCGACTTCAAGGAGGACGGCAACATCCTGGGG
CACAAGCTGGAGTACAACGCCATCAGCGACAACGTCTATATCACCGCCGACAAGCAGAAGA
ACGGCATCAAGGCCAACTTCAAGATCCGCCACAACATCGAGGACGGCAGCGTGCAGCTCG
CCGACCACTACCAGCAGAACACCCCCATCGGCGACGGCCCCGTGCTGCTGCCCGACAAC
CACTACCTGAGCACCCAGTCCAAGCTGAGCAAAGACCCCAACGAGAAGCGCGATCACATG
GTCCTGCTGGAGTTCGTGACCGCCGCCGGGATCACTCTCGGCATGGACGAGCTGTACAAG
GGCGGTGGCTCATCTGGCGGAGGTGAGATGTGGCATGAAGGCCTGGAAGAGGCATCTCGT
TTGTACTTTGGGGAAAGGAACGTGAAAGGCATGTTTGAGGTGCTGGAGCCCTTGCATGCTA
TGATGGAACGGGGCCCCCAGACTCTGAAGGAAACATCCTTTAATCAGGCCTATGGTCGAGA
TTTAATGGAGGCCCAAGAGTGGTGCAGGAAGTACATGAAATCAGGGAATGTCAAGGACCTC
ACCCAAGCCTGGGACCTCTATTATCATGTGTTCCGACGAATCTCAAAGCAGCAGATCTCGTA
CGCGTCCCGGGGCGGTGGCTCATCTGGCGGAGGTGAGCTCCAGACAATTAAGTGTGTTGT
TGTGGGCGATGGTGCTTAA---***IRES region***---
ATGGGGGTTGGTAAAAACTGTCTCCTGATATCCTACACAACAAACAAATTTCCATCGGAGTAT
GTACCGACTGTTTTTGACAACTATGCAGTCACAGTTATGATTGGTGGAGAACCATATACTCTT
GGACTTTTTGATACTGCAGGGCAAGAGGATTATGACAGATTACGACCGCTGAGTTATCCACA
AACAGATGTATTTCTAGTCTGTTTTTCAGTGGTCTCTCCATCTTCATTTGAAAACGTGAAAGA
AAAGTGGGTGCCTGAGATAACTCACCACTGTCCAAAGACTCCTTTCTTGCTTGTTGGGACT
CAAATTGATCTCAGAGATGACCCCTCTACTATTGAGAAACTTGCCAAGAACAAACAGAAGCC
TATCACTCCAGAGACTGCTGAAAAGCTGGCCCGTGACCTGAAGGCTGTCAAGTATGTGGAG

TGTTCTGCACTTACACAGAGAGGTCTGAAGAATGTGTTTGATGAGGCTATCCTAGCTGCCCT
CGAGCCTCCGGAAACTCAACCCGGGGATCCCAATTGGGAGCTCGTGTACACGGCGCGCCT
GCAGGGAGGTGGCTCATCTGGCGGAGGTCAGATCTCGTACGCGTCCCGGGGCGGAGTGC
AGGTGGAAACCATCTCCCCAGGAGACGGGCGCACCTTCCCCAAGCGCGGCCAGACCTGC
GTGGTGCACTACACCGGGATGCTTGAAGATGGAAAGAAATTTGATTCCTCCCGGGACAGAA
ACAAGCCCTTTAAGTTTATGCTAGGCAAGCAGGAGGTGATCCGAGGCTGGGAAGAAGGGG
TTGCCCAGATGAGTGTGGGTCAGAGAGCCAAACTGACTATATCTCCAGATTATGCCTATGGT
GCCACTGGGCACCCAGGCATCATCCCACCACATGCCACTCTCGTCTTCGATGTGGAGCTTC
TAAAACTGGAAGGCGGTGGCTCATCTGGCGGAGGTGTGAGCAAGGGCGAGGAGCTGTTC
ACCGGGGTGGTGCCCATCCTGGTCGAGCTGGACGGCGACGTAAACGGCCACAAGTTCAG
CGTGTCCGGCGAGGGCGAGGGCGATGCCACCTACGGCAAGCTGACCCTGAAGCTCATCT
GCACCACCGGCAAGCTGCCCGTGCCCTGGCCCACCCTCGTGACCACCCTCGGCTACGGC
CTGCAGTGCTTCGCCCGCTACCCCGACCACATGAAGCAGCACGACTTCTTCAAGTCCGCC
ATGCCCGAAGGCTACGTCCAGGAGCGCACCATCTTCTTCAAGGACGACGGCAACTACAAG
ACCCGCGCCGAGGTGAAGTTCGAGGGCGACACCCTGGTGAACCGCATCGAGCTGAAGGG
CATCGACTTCAAGGAGGACGGCAACATCCTGGGGCACAAGCTGGAGTACAACTACAACAG
CCACAACGTCTATATCACCGCCGACAAGCAGAAGAACGGCATCAAGGCCAACTTCAAGATC
CGCCACAACATCGAGGACGGCGGCGTGCAGCTCGCCGACCACTACCAGCAGAACACCCC
CATCGGCGACGGCCCCGTGCTGCTGCCCGACAACCACTACCTGAGCTACCAGTCCAAGCT
GAGCAAAGACCCCAACGAGAAGCGCGATCACATGGTCCTGCTGGAGTTCGTGACCGCCGC
CGGGATCACTCTCGGCATGGACGAGCTGTACAAGAAAAAAAAGAAGAAAAAGAGCAAGAC
CAAATGCGTGATTATGTAA

#### 3）**pIRES-mCerulean-FRB-Rac1/N12-Rac1/13C(Q61L)-FKBP-mVenus-CAAX**

>Amino acid sequence (61L is underlined)

MVSKGEELFTGVVPILVELDGDVNGHKFSVSGEGEGDATYGKLTLKFICTTGKLPVPWPTLVTT
LTWGVQCFARYPDHMKQHDFFKSAMPEGYVQERTIFFKDDGNYKTRAEVKFEGDTLVNRIELK
GIDFKEDGNILGHKLEYNAISDNVYITADKQKNGIKANFKIRHNIEDGSVQLADHYQQNTPIGDGP
VLLPDNHYLSTQSKLSKDPNEKRDHMVLLEFVTAAGITLGMDELYKGGGSSGGGEMWHEGLE
EASRLYFGERNVKGMFEVLEPLHAMMERGPQTLKETSFNQAYGRDLMEAQEWCRKYMKSGN
VKDLTQAWDLYYHVFRRISKQQISYASRGGGSSGGGELQAIKCVVVGDGA*---**_IRES Region_**---
MVGKTCLLISYTTNAFPGEYIPTVFDNYSANVMVDGKPVNLGLWDTAGLEDYDRLRPLSYPQT
DVFLICFSLVSPASFENVRAKWYPEVRHHCPNTPIILVGTKLDLRDDKDTIEKLKEKKLTPITYPQ
GLAMAKEIGAVKYLECSALTQRGLKTVFDEAIRAVLCPPPVGDPNWELVYTARLQGGGSSGGG
QISYASRGGVQVETISPGDGRTFPKRGQTCVVHYTGMLEDGKKFDSSRDRNKPFKFMLGKQE
VIRGWEEGVAQMSVGQRAKLTISPDYAYGATGHPGIIPPHATLVFDVELLKLEGGGSSGGGVSK
GEELFTGVVPILVELDGDVNGHKFSVSGEGEGDATYGKLTLKLICTTGKLPVPWPTLVTTLGYGL
QCFARYPDHMKQHDFFKSAMPEGYVQERTIFFKDDGNYKTRAEVKFEGDTLVNRIELKGIDFKE
DGNILGHKLEYNYNSHNVYITADKQKNGIKANFKIRHNIEDGGVQLADHYQQNTPIGDGPVLLPD
NHYLSYQSKLSKDPNEKRDHMVLLEFVTAAGITLGMDELYKKKKKKKSKTKCVIM*

>DNA sequence

ATGGTGAGCAAGGGCGAGGAGCTGTTCACCGGGGTGGTGCCCATCCTGGTCGAGCTGGA
CGGCGACGTAAACGGCCACAAGTTCAGCGTGTCCGGCGAGGGCGAGGGCGATGCCACCT
ACGGCAAGCTGACCCTGAAGTTCATCTGCACCACCGGCAAGCTGCCCGTGCCCTGGCCCA
CCCTCGTGACCACCCTGACCTGGGGCGTGCAGTGCTTCGCCCGCTACCCCGACCACATGA
AGCAGCACGACTTCTTCAAGTCCGCCATGCCCGAAGGCTACGTCCAGGAGCGCACCATCT
TCTTCAAGGACGACGGCAACTACAAGACCCGCGCCGAGGTGAAGTTCGAGGGCGACACC
CTGGTGAACCGCATCGAGCTGAAGGGCATCGACTTCAAGGAGGACGGCAACATCCTGGGG
CACAAGCTGGAGTACAACGCCATCAGCGACAACGTCTATATCACCGCCGACAAGCAGAAGA
ACGGCATCAAGGCCAACTTCAAGATCCGCCACAACATCGAGGACGGCAGCGTGCAGCTCG
CCGACCACTACCAGCAGAACACCCCCATCGGCGACGGCCCCGTGCTGCTGCCCGACAAC
CACTACCTGAGCACCCAGTCCAAGCTGAGCAAAGACCCCAACGAGAAGCGCGATCACATG
GTCCTGCTGGAGTTCGTGACCGCCGCCGGGATCACTCTCGGCATGGACGAGCTGTACAAG
GGCGGTGGCTCATCTGGCGGAGGTGAGATGTGGCATGAAGGCCTGGAAGAGGCATCTCGT
TTGTACTTTGGGGAAAGGAACGTGAAAGGCATGTTTGAGGTGCTGGAGCCCTTGCATGCTA
TGATGGAACGGGGCCCCCAGACTCTGAAGGAAACATCCTTTAATCAGGCCTATGGTCGAGA
TTTAATGGAGGCCCAAGAGTGGTGCAGGAAGTACATGAAATCAGGGAATGTCAAGGACCTC
ACCCAAGCCTGGGACCTCTATTATCATGTGTTCCGACGAATCTCAAAGCAGCAGATCTCGTA
CGCGTCCCGGGGCGGTGGCTCATCTGGCGGAGGTGAGCTCCAGGCCATCAAGTGTGTGG
TGGTGGGAGACGGAGCTTAA---**_IRES region_**---
ATGGTAGGTAAAACTTGCCTACTGATCAGTTACACAACCAATGCATTTCCTGGAGAATATATC
CCTACTGTCTTTGACAATTATTCTGCCAATGTTATGGTAGATGGAAAACCGGTGAATCTGGGC
TTATGGGATACAGCTGGACTAGAAGATTATGACAGATTACGCCCCCTATCCTATCCGCAAACA
GATGTGTTCTTAATTTGCTTTTCCCTTGTGAGTCCTGCATCATTTGAAAATGTCCGTGCAAAG
TGGTATCCTGAGGTGCGGCACCACTGTCCCAACACTCCCATCATCCTAGTGGGAACTAAAC
TTGATCTTAGGGATGATAAAGACACGATCGAGAAACTGAAGGAGAAGAAGCTGACTCCCATC
ACCTATCCGCAGGGTCTAGCCATGGCTAAGGAGATTGGTGCTGTAAAATACCTGGAGTGCT

CGGCGCTCACACAGCGAGGCCTCAAGACAGTGTTTGACGAAGCGATCCGAGCAGTCCTCT
GCCCGCCTCCCGTGGGGGATCCCAATTGGGAGCTCGTGTACACGGCGCGCCTGCAGGGA
GGTGGCTCATCTGGCGGAGGTCAGATCTCGTACGCGTCCCGGGGCGGAGTGCAGGTGGA
AACCATCTCCCCAGGAGACGGGCGCACCTTCCCCAAGCGCGGCCAGACCTGCGTGGTGC
ACTACACCGGGATGCTTGAAGATGGAAAGAAATTTGATTCCTCCCGGGACAGAAACAAGCC
CTTTAAGTTTATGCTAGGCAAGCAGGAGGTGATCCGAGGCTGGGAAGAAGGGGTTGCCCA
GATGAGTGTGGGTCAGAGAGCCAAACTGACTATATCTCCAGATTATGCCTATGGTGCCACTG
GGCACCCAGGCATCATCCCACCACATGCCACTCTCGTCTTCGATGTGGAGCTTCTAAAACT
GGAAGGCGGTGGCTCATCTGGCGGAGGTGTGAGCAAGGGCGAGGAGCTGTTCACCGGG
GTGGTGCCCATCCTGGTCGAGCTGGACGGCGACGTAAACGGCCACAAGTTCAGCGTGTCC
GGCGAGGGCGAGGGCGATGCCACCTACGGCAAGCTGACCCTGAAGCTCATCTGCACCAC
CGGCAAGCTGCCCGTGCCCTGGCCCACCCTCGTGACCACCCTCGGCTACGGCCTGCAGT
GCTTCGCCCGCTACCCCGACCACATGAAGCAGCACGACTTCTTCAAGTCCGCCATGCCCG
AAGGCTACGTCCAGGAGCGCACCATCTTCTTCAAGGACGACGGCAACTACAAGACCCGCG
CCGAGGTGAAGTTCGAGGGCGACACCCTGGTGAACCGCATCGAGCTGAAGGGCATCGAC
TTCAAGGAGGACGGCAACATCCTGGGGCACAAGCTGGAGTACAACTACAACAGCCACAAC
GTCTATATCACCGCCGACAAGCAGAAGAACGGCATCAAGGCCAACTTCAAGATCCGCCACA
ACATCGAGGACGGCGGCGTGCAGCTCGCCGACCACTACCAGCAGAACACCCCCATCGGC
GACGGCCCCGTGCTGCTGCCCGACAACCACTACCTGAGCTACCAGTCCAAGCTGAGCAAA
GACCCCAACGAGAAGCGCGATCACATGGTCCTGCTGGAGTTCGTGACCGCCGCCGGGATC
ACTCTCGGCATGGACGAGCTGTACAAGAAAAAAAAGAAGAAAAAGAGCAAGACCAAATGCG
TGATTATGTAA

4）**pIRES-mCerulean-FRB-Rac1/N12-Rac1/13C(T17N)-FKBP-mVenus-CAAX**

>Amino acid sequence (17N is underlined)

MVSKGEELFTGVVPILVELDGDVNGHKFSVSGEGEGDATYGKLTLKFICTTGKLPVPWPTLVTT
LTWGVQCFARYPDHMKQHDFFKSAMPEGYVQERTIFFKDDGNYKTRAEVKFEGDTLVNRIELK
GIDFKEDGNILGHKLEYNAISDNVYITADKQKNGIKANFKIRHNIEDGSVQLADHYQQNTPIGDGP
VLLPDNHYLSTQSKLSKDPNEKRDHMVLLEFVTAAGITLGMDELYKGGGSSGGGEMWHEGLE
EASRLYFGERNVKGMFEVLEPLHAMMERGPQTLKETSFNQAYGRDLMEAQEWCRKYMKSGN
VKDLTQAWDLYYHVFRRISKQQISYASRGGGSSGGGELQAIKCVVVGDGA*---*IRES Region*---
MVGK**N**CLLISYTTNAFPGEYIPTVFDNYSANVMVDGKPVNLGLWDTAGQEDYDRLRPLSYPQT
DVFLICFSLVSPASFENVRAKWYPEVRHHCPNTPIILVGTKLDLRDDKDTIEKLKEKKLTPITYPQ
GLAMAKEIGAVKYLECSALTQRGLKTVFDEAIRAVLCPPPVGDPNWELVYTARLQGGGSSGGG
QISYASRGGVQVETISPGDGRTFPKRGQTCVVHYTGMLEDGKKFDSSRDRNKPFKFMLGKQE
VIRGWEEGVAQMSVGQRAKLTISPDYAYGATGHPGIIPPHATLVFDVELLKLEGGGSSGGGVSK
GEELFTGVVPILVELDGDVNGHKFSVSGEGEGDATYGKLTLKLICTTGKLPVPWPTLVTTLGYGL
QCFARYPDHMKQHDFFKSAMPEGYVQERTIFFKDDGNYKTRAEVKFEGDTLVNRIELKGIDFKE
DGNILGHKLEYNYNSHNVYITADKQKNGIKANFKIRHNIEDGGVQLADHYQQNTPIGDGPVLLPD
NHYLSYQSKLSKDPNEKRDHMVLLEFVTAAGITLGMDELYKKKKKKKSKTKCVIM*

>DNA sequence

ATGGTGAGCAAGGGCGAGGAGCTGTTCACCGGGGTGGTGCCCATCCTGGTCGAGCTGGA
CGGCGACGTAAACGGCCACAAGTTCAGCGTGTCCGGCGAGGGCGAGGGCGATGCCACCT
ACGGCAAGCTGACCCTGAAGTTCATCTGCACCACCGGCAAGCTGCCCGTGCCCTGGCCCA
CCCTCGTGACCACCCTGACCTGGGGCGTGCAGTGCTTCGCCCGCTACCCCGACCACATGA
AGCAGCACGACTTCTTCAAGTCCGCCATGCCCGAAGGCTACGTCCAGGAGCGCACCATCT
TCTTCAAGGACGACGGCAACTACAAGACCCGCGCCGAGGTGAAGTTCGAGGGCGACACC
CTGGTGAACCGCATCGAGCTGAAGGGCATCGACTTCAAGGAGGACGGCAACATCCTGGGG
CACAAGCTGGAGTACAACGCCATCAGCGACAACGTCTATATCACCGCCGACAAGCAGAAGA
ACGGCATCAAGGCCAACTTCAAGATCCGCCACAACATCGAGGACGGCAGCGTGCAGCTCG
CCGACCACTACCAGCAGAACACCCCCATCGGCGACGGCCCCGTGCTGCTGCCCGACAAC
CACTACCTGAGCACCCAGTCCAAGCTGAGCAAAGACCCCAACGAGAAGCGCGATCACATG
GTCCTGCTGGAGTTCGTGACCGCCGCCGGGATCACTCTCGGCATGGACGAGCTGTACAAG
GGCGGTGGCTCATCTGGCGGAGGTGAGATGTGGCATGAAGGCCTGGAAGAGGCATCTCGT
TTGTACTTTGGGGAAAGGAACGTGAAAGGCATGTTTGAGGTGCTGGAGCCCTTGCATGCTA
TGATGGAACGGGGCCCCCAGACTCTGAAGGAAACATCCTTTAATCAGGCCTATGGTCGAGA
TTTAATGGAGGCCCAAGAGTGGTGCAGGAAGTACATGAAATCAGGGAATGTCAAGGACCTC
ACCCAAGCCTGGGACCTCTATTATCATGTGTTCCGACGAATCTCAAAGCAGCAGATCTCGTA
CGCGTCCCGGGGCGGTGGCTCATCTGGCGGAGGTGAGCTCCAGGCCATCAAGTGTGTGG
TGGTGGGAGACGGAGCTTAA---***IRES region***---
ATGGTAGGTAAAAACTGCCTACTGATCAGTTACACAACCAATGCATTTCCTGGAGAATATATC
CCTACTGTCTTTGACAATTATTCTGCCAATGTTATGGTAGATGGAAAACCGGTGAATCTGGGC
TTATGGGATACAGCTGGACAAGAAGATTATGACAGATTACGCCCCCTATCCTATCCGCAAACA
GATGTGTTCTTAATTTGCTTTTCCCTTGTGAGTCCTGCATCATTTGAAAATGTCCGTGCAAAG
TGGTATCCTGAGGTGCGGCACCACTGTCCCAACACTCCCATCATCCTAGTGGGAACTAAAC
TTGATCTTAGGGATGATAAAGACACGATCGAGAAACTGAAGGAGAAGAAGCTGACTCCCATC
ACCTATCCGCAGGGTCTAGCCATGGCTAAGGAGATTGGTGCTGTAAAATACCTGGAGTGCT

CGGCGCTCACACAGCGAGGCCTCAAGACAGTGTTTGACGAAGCGATCCGAGCAGTCCTCT
GCCCGCCTCCCGTGGGGGATCCCAATTGGGAGCTCGTGTACACGGCGCGCCTGCAGGGA
GGTGGCTCATCTGGCGGAGGTCAGATCTCGTACGCGTCCCGGGGCGGAGTGCAGGTGGA
AACCATCTCCCCAGGAGACGGGCGCACCTTCCCCAAGCGCGGCCAGACCTGCGTGGTGC
ACTACACCGGGATGCTTGAAGATGGAAAGAAATTTGATTCCTCCCGGGACAGAAACAAGCC
CTTTAAGTTTATGCTAGGCAAGCAGGAGGTGATCCGAGGCTGGGAAGAAGGGGTTGCCCA
GATGAGTGTGGGTCAGAGAGCCAAACTGACTATATCTCCAGATTATGCCTATGGTGCCACTG
GGCACCCAGGCATCATCCCACCACATGCCACTCTCGTCTTCGATGTGGAGCTTCTAAAACT
GGAAGGCGGTGGCTCATCTGGCGGAGGTGTGAGCAAGGGCGAGGAGCTGTTCACCGGG
GTGGTGCCCATCCTGGTCGAGCTGGACGGCGACGTAAACGGCCACAAGTTCAGCGTGTCC
GGCGAGGGCGAGGGCGATGCCACCTACGGCAAGCTGACCCTGAAGCTCATCTGCACCAC
CGGCAAGCTGCCCGTGCCCTGGCCCACCCTCGTGACCACCCTCGGCTACGGCCTGCAGT
GCTTCGCCCGCTACCCCGACCACATGAAGCAGCACGACTTCTTCAAGTCCGCCATGCCCG
AAGGCTACGTCCAGGAGCGCACCATCTTCTTCAAGGACGACGGCAACTACAAGACCCGCG
CCGAGGTGAAGTTCGAGGGCGACACCCTGGTGAACCGCATCGAGCTGAAGGGCATCGAC
TTCAAGGAGGACGGCAACATCCTGGGGCACAAGCTGGAGTACAACTACAACAGCCACAAC
GTCTATATCACCGCCGACAAGCAGAAGAACGGCATCAAGGCCAACTTCAAGATCCGCCACA
ACATCGAGGACGGCGGCGTGCAGCTCGCCGACCACTACCAGCAGAACACCCCCATCGGC
GACGGCCCCGTGCTGCTGCCCGACAACCACTACCTGAGCTACCAGTCCAAGCTGAGCAAA
GACCCCAACGAGAAGCGCGATCACATGGTCCTGCTGGAGTTCGTGACAAAAAAAAGAAGA
AAAAGAGCAAGACCAAATGCGTGATTATGTAA

5）**pIRES-mCerulean-FRB-RhoA/N12-RhoA/13C(Q63L)-FKBP-mVenus-CAAX**

>Amino acid sequence (63L is underlined)

MVSKGEELFTGVVPILVELDGDVNGHKFSVSGEGEGDATYGKLTLKFICTTGKLPVPWPTLVTT
LTWGVQCFARYPDHMKQHDFFKSAMPEGYVQERTIFFKDDGNYKTRAEVKFEGDTLVNRIELK
GIDFKEDGNILGHKLEYNAISDNVYITADKQKNGIKANFKIRHNIEDGSVQLADHYQQNTPIGDGP
VLLPDNHYLSTQSKLSKDPNEKRDHMVLLEFVTAAGITLGMDELYKGGGSSGGGEMWHEGLE
EASRLYFGERNVKGMFEVLEPLHAMMERGPQTLKETSFNQAYGRDLMEAQEWCRKYMKSGN
VKDLTQAWDLYYHVFRRISKQQISYASRGGGSSGGGELAAIRKKLVIVGDGA*---***IRES Region***--
MCGKTCLLIVFSKDQFPEVYVPTVFENYVADIEVDGKQVELALWDTAG<u>**L**</u>EDYDRLRPLSYPDTD
VILMCFSIDSPDSLENIPEKWTPEVKHFCPNVPIILVGNKKDLRNDEHTRRELAKMKQEPVKPEE
GRDMANRIGAFGYMECSAKTKDGVREVFEMATRAALQAGDPNWELVYTARLQGGGSSGGGQ
ISYASRGGVQVETISPGDGRTFPKRGQTCVVHYTGMLEDGKKFDSSRDRNKPFKFMLGKQEVI
RGWEEGVAQMSVGQRAKLTISPDYAYGATGHPGIIPPHATLVFDVELLKLEGGGSSGGGVSKG
EELFTGVVPILVELDGDVNGHKFSVSGEGEGDATYGKLTLKLICTTGKLPVPWPTLVTTLGYGLQ
CFARYPDHMKQHDFFKSAMPEGYVQERTIFFKDDGNYKTRAEVKFEGDTLVNRIELKGIDFKED
GNILGHKLEYNYNSHNVYITADKQKNGIKANFKIRHNIEDGGVQLADHYQQNTPIGDGPVLLPDN
HYLSYQSKLSKDPNEKRDHMVLLEFVTAAGITLGMDELYKKKKKKKSKTKCVIM*

>DNA sequence

ATGGTGAGCAAGGGCGAGGAGCTGTTCACCGGGGTGGTGCCCATCCTGGTCGAGCTGGA
CGGCGACGTAAACGGCCACAAGTTCAGCGTGTCCGGCGAGGGCGAGGGCGATGCCACCT
ACGGCAAGCTGACCCTGAAGTTCATCTGCACCACCGGCAAGCTGCCCGTGCCCTGGCCCA
CCCTCGTGACCACCCTGACCTGGGGCGTGCAGTGCTTCGCCCGCTACCCCGACCACATGA
AGCAGCACGACTTCTTCAAGTCCGCCATGCCCGAAGGCTACGTCCAGGAGCGCACCATCT
TCTTCAAGGACGACGGCAACTACAAGACCCGCGCCGAGGTGAAGTTCGAGGGCGACACC
CTGGTGAACCGCATCGAGCTGAAGGGCATCGACTTCAAGGAGGACGGCAACATCCTGGGG
CACAAGCTGGAGTACAACGCCATCAGCGACAACGTCTATATCACCGCCGACAAGCAGAAGA
ACGGCATCAAGGCCAACTTCAAGATCCGCCACAACATCGAGGACGGCAGCGTGCAGCTCG
CCGACCACTACCAGCAGAACACCCCCATCGGCGACGGCCCCGTGCTGCTGCCCGACAAC
CACTACCTGAGCACCCAGTCCAAGCTGAGCAAAGACCCCAACGAGAAGCGCGATCACATG
GTCCTGCTGGAGTTCGTGACCGCCGCCGGGATCACTCTCGGCATGGACGAGCTGTACAAG
GGCGGTGGCTCATCTGGCGGAGGTGAGATGTGGCATGAAGGCCTGGAAGAGGCATCTCGT
TTGTACTTTGGGGAAAGGAACGTGAAAGGCATGTTTGAGGTGCTGGAGCCCTTGCATGCTA
TGATGGAACGGGGCCCCCAGACTCTGAAGGAAACATCCTTTAATCAGGCCTATGGTCGAGA
TTTAATGGAGGCCCAAGAGTGGTGCAGGAAGTACATGAAATCAGGGAATGTCAAGGACCTC
ACCCAAGCCTGGGACCTCTATTATCATGTGTTCCGACGAATCTCAAAGCAGCAGATCTCGTA
CGCGTCCCGGGGCGGTGGCTCATCTGGCGGAGGTGAGCTCGCTGCCATCCGGAAGAAAC
TGGTGATTGTTGGTGATGGAGCCTAA---***IRES region***---
ATGTGTGGAAAGACATGCTTGCTCATAGTCTTCAGCAAGGACCAGTTCCCAGAGGTGTATGT
GCCCACAGTGTTTGAGAACTATGTGGCAGATATCGAGGTGGATGGAAAGCAGGTAGAGTTG
GCTTTGTGGGACACAGCTGGGCTGGAAGATTATGATCGCCTGAGGCCCCTCTCCTACCCAG
ATACCGATGTTATACTGATGTGTTTTTCCATCGACAGCCCTGATAGTTTAGAAAACATCCCAG
AAAAGTGGACCCCAGAAGTCAAGCATTTCTGTCCCAACGTGCCCATCATCCTGGTTGGGAA
TAAGAAGGATCTTCGGAATGATGAGCACACAAGGCGGGAGCTAGCCAAGATGAAGCAGGA

GCCGGTGAAACCTGAAGAAGGCAGAGATATGGCAAACAGGATTGGCGCTTTTGGGTACATG
GAGTGTTCAGCAAAGACCAAAGATGGAGTGAGAGAGGTTTTTGAAATGGCTACGAGAGCTG
CTCTGCAAGCTGGGGATCCCAATTGGGAGCTCGTGTACACGGCGCGCCTGCAGGGAGGT
GGCTCATCTGGCGGAGGTCAGATCTCGTACGCGTCCCGGGGCGGAGTGCAGGTGGAAAC
CATCTCCCCAGGAGACGGGCGCACCTTCCCCAAGCGCGGCCAGACCTGCGTGGTGCACT
ACACCGGGATGCTTGAAGATGGAAAGAAATTTGATTCCTCCCGGGACAGAAACAAGCCCTT
TAAGTTTATGCTAGGCAAGCAGGAGGTGATCCGAGGCTGGGAAGAAGGGGTTGCCCAGAT
GAGTGTGGGTCAGAGAGCCAAACTGACTATATCTCCAGATTATGCCTATGGTGCCACTGGG
CACCCAGGCATCATCCCACCACATGCCACTCTCGTCTTCGATGTGGAGCTTCTAAAACTGG
AAGGCGGTGGCTCATCTGGCGGAGGTGTGAGCAAGGGCGAGGAGCTGTTCACCGGGGTG
GTGCCCATCCTGGTCGAGCTGGACGGCGACGTAAACGGCCACAAGTTCAGCGTGTCCGG
CGAGGGCGAGGGCGATGCCACCTACGGCAAGCTGACCCTGAAGCTCATCTGCACCACCG
GCAAGCTGCCCGTGCCCTGGCCCACCCTCGTGACCACCCTCGGCTACGGCCTGCAGTGC
TTCGCCCGCTACCCCGACCACATGAAGCAGCACGACTTCTTCAAGTCCGCCATGCCCGAA
GGCTACGTCCAGGAGCGCACCATCTTCTTCAAGGACGACGGCAACTACAAGACCCGCGCC
GAGGTGAAGTTCGAGGGCGACACCCTGGTGAACCGCATCGAGCTGAAGGGCATCGACTTC
AAGGAGGACGGCAACATCCTGGGGCACAAGCTGGAGTACAACTACAACAGCCACAACGTC
TATATCACCGCCGACAAGCAGAAGAACGGCATCAAGGCCAACTTCAAGATCCGCCACAACA
TCGAGGACGGCGGCGTGCAGCTCGCCGACCACTACCAGCAGAACACCCCCATCGGCGAC
GGCCCCGTGCTGCTGCCCGACAACCACTACCTGAGCTACCAGTCCAAGCTGAGCAAAGAC
CCCAACGAGAAGCGCGATCACATGGTCCTGCTGGAGTTCGTGACCGCCGCCGGGATCACT
CTCGGCATGGACGAGCTGTACAAGAAAAAAAAGAAGAAAAAGAGCAAGACCAAATGCGTGA
TTATGTAA

6）**pIRES-mCerulean-FRB-RhoA/N12-RhoA/13C(T19N)-FKBP-mVenus-CAAX**

>Amino acid sequence (19N is underlined)

MVSKGEELFTGVVPILVELDGDVNGHKFSVSGEGEGDATYGKLTLKFICTTGKLPVPWPTLVTT
LTWGVQCFARYPDHMKQHDFFKSAMPEGYVQERTIFFKDDGNYKTRAEVKFEGDTLVNRIELK
GIDFKEDGNILGHKLEYNAISDNVYITADKQKNGIKANFKIRHNIEDGSVQLADHYQQNTPIGDGP
VLLPDNHYLSTQSKLSKDPNEKRDHMVLLEFVTAAGITLGMDELYKGGGSSGGGEMWHEGLE
EASRLYFGERNVKGMFEVLEPLHAMMERGPQTLKETSFNQAYGRDLMEAQEWCRKYMKSGN
VKDLTQAWDLYYHVFRRISKQQISYASRGGGSSGGGELAAIRKKLVIVGDGA*---***IRES Region***--
MCGK**N**CLLIVFSKDQFPEVYVPTVFENYVADIEVDGKQVELALWDTAGQEDYDRLRPLSYPDT
DVILMCFSIDSPDSLENIPEKWTPEVKHFCPNVPIILVGNKKDLRNDEHTRRELAKMKQEPVKPE
EGRDMANRIGAFGYMECSAKTKDGVREVFEMATRAALQAGDPNWELVYTARLQGGGSSGGG
QISYASRGGVQVETISPGDGRTFPKRGQTCVVHYTGMLEDGKKFDSSRDRNKPFKFMLGKQE
VIRGWEEGVAQMSVGQRAKLTISPDYAYGATGHPGIIPPHATLVFDVELLKLEGGGSSGGGVSK
GEELFTGVVPILVELDGDVNGHKFSVSGEGEGDATYGKLTLKLICTTGKLPVPWPTLVTTLGYGL
QCFARYPDHMKQHDFFKSAMPEGYVQERTIFFKDDGNYKTRAEVKFEGDTLVNRIELKGIDFKE
DGNILGHKLEYNYNSHNVYITADKQKNGIKANFKIRHNIEDGGVQLADHYQQNTPIGDGPVLLPD
NHYLSYQSKLSKDPNEKRDHMVLLEFVTAAGITLGMDELYKKKKKKKSKTKCVIM*

>DNA sequence

ATGGTGAGCAAGGGCGAGGAGCTGTTCACCGGGGTGGTGCCCATCCTGGTCGAGCTGGA
CGGCGACGTAAACGGCCACAAGTTCAGCGTGTCCGGCGAGGGCGAGGGCGATGCCACCT
ACGGCAAGCTGACCCTGAAGTTCATCTGCACCACCGGCAAGCTGCCCGTGCCCTGGCCCA
CCCTCGTGACCACCCTGACCTGGGGCGTGCAGTGCTTCGCCCGCTACCCCGACCACATGA
AGCAGCACGACTTCTTCAAGTCCGCCATGCCCGAAGGCTACGTCCAGGAGCGCACCATCT
TCTTCAAGGACGACGGCAACTACAAGACCCGCGCCGAGGTGAAGTTCGAGGGCGACACC
CTGGTGAACCGCATCGAGCTGAAGGGCATCGACTTCAAGGAGGACGGCAACATCCTGGGG
CACAAGCTGGAGTACAACGCCATCAGCGACAACGTCTATATCACCGCCGACAAGCAGAAGA
ACGGCATCAAGGCCAACTTCAAGATCCGCCACAACATCGAGGACGGCAGCGTGCAGCTCG
CCGACCACTACCAGCAGAACACCCCCATCGGCGACGGCCCCGTGCTGCTGCCCGACAAC
CACTACCTGAGCACCCAGTCCAAGCTGAGCAAAGACCCCAACGAGAAGCGCGATCACATG
GTCCTGCTGGAGTTCGTGACCGCCGCCGGGATCACTCTCGGCATGGACGAGCTGTACAAG
GGCGGTGGCTCATCTGGCGGAGGTGAGATGTGGCATGAAGGCCTGGAAGAGGCATCTCGT
TTGTACTTTGGGGAAAGGAACGTGAAAGGCATGTTTGAGGTGCTGGAGCCCTTGCATGCTA
TGATGGAACGGGGCCCCCAGACTCTGAAGGAAACATCCTTTAATCAGGCCTATGGTCGAGA
TTTAATGGAGGCCCAAGAGTGGTGCAGGAAGTACATGAAATCAGGGAATGTCAAGGACCTC
ACCCAAGCCTGGGACCTCTATTATCATGTGTTCCGACGAATCTCAAAGCAGCAGATCTCGTA
CGCGTCCCGGGGCGGTGGCTCATCTGGCGGAGGTGAGCTCGCTGCCATCCGGAAGAAAC
TGGTGATTGTTGGTGATGGAGCCTAA---***IRES region***---
ATGTGTGGAAAGAACTGCTTGCTCATAGTCTTCAGCAAGGACCAGTTCCCAGAGGTGTATG
TGCCCACAGTGTTTGAGAACTATGTGGCAGATATCGAGGTGGATGGAAAGCAGGTAGAGTT
GGCTTTGTGGGACACAGCTGGGCAGGAAGATTATGATCGCCTGAGGCCCCTCTCCTACCC
AGATACCGATGTTATACTGATGTGTTTTTCCATCGACAGCCCTGATAGTTTAGAAAACATCCC
AGAAAAGTGGACCCCAGAAGTCAAGCATTTCTGTCCCAACGTGCCCATCATCCTGGTTGGG
AATAAGAAGGATCTTCGGAATGATGAGCACACAAGGCGGGAGCTAGCCAAGATGAAGCAGG
AGCCGGTGAAACCTGAAGAAGGCAGAGATATGGCAAACAGGATTGGCGCTTTTGGGTACAT
GGAGTGTTCAGCAAAGACCAAAGATGGAGTGAGAGAGGTTTTTGAAATGGCTACGAGAGCT

GCTCTGCAAGCTGGGGATCCCAATTGGGAGCTCGTGTACACGGCGCGCCTGCAGGGAGG
TGGCTCATCTGGCGGAGGTCAGATCTCGTACGCGTCCCGGGGCGGAGTGCAGGTGGAAA
CCATCTCCCCAGGAGACGGGCGCACCTTCCCCAAGCGCGGCCAGACCTGCGTGGTGCAC
TACACCGGGATGCTTGAAGATGGAAAGAAATTTGATTCCTCCCGGGACAGAAACAAGCCCT
TTAAGTTTATGCTAGGCAAGCAGGAGGTGATCCGAGGCTGGGAAGAAGGGGTTGCCCAGA
TGAGTGTGGGTCAGAGAGCCAAACTGACTATATCTCCAGATTATGCCTATGGTGCCACTGGG
CACCCAGGCATCATCCCACCACATGCCACTCTCGTCTTCGATGTGGAGCTTCTAAAACTGG
AAGGCGGTGGCTCATCTGGCGGAGGTGTGAGCAAGGGCGAGGAGCTGTTCACCGGGGTG
GTGCCCATCCTGGTCGAGCTGGACGGCGACGTAAACGGCCACAAGTTCAGCGTGTCCGG
CGAGGGCGAGGGCGATGCCACCTACGGCAAGCTGACCCTGAAGCTCATCTGCACCACCG
GCAAGCTGCCCGTGCCCTGGCCCACCCTCGTGACCACCCTCGGCTACGGCCTGCAGTGC
TTCGCCCGCTACCCCGACCACATGAAGCAGCACGACTTCTTCAAGTCCGCCATGCCCGAA
GGCTACGTCCAGGAGCGCACCATCTTCTTCAAGGACGACGGCAACTACAAGACCCGCGCC
GAGGTGAAGTTCGAGGGCGACACCCTGGTGAACCGCATCGAGCTGAAGGGCATCGACTTC
AAGGAGGACGGCAACATCCTGGGGCACAAGCTGGAGTACAACTACAACAGCCACAACGTC
TATATCACCGCCGACAAGCAGAAGAACGGCATCAAGGCCAACTTCAAGATCCGCCACAACA
TCGAGGACGGCGGCGTGCAGCTCGCCGACCACTACCAGCAGAACACCCCCATCGGCGAC
GGCCCCGTGCTGCTGCCCGACAACCACTACCTGAGCTACCAGTCCAAGCTGAGCAAAGAC
CCCAACGAGAAGCGCGATCACATGGTCCTGCTGGAGTTCGTGACCGCCGCCGGGATCACT
CTCGGCATGGACGAGCTGTACAAGAAAAAAAAGAAGAAAAAGAGCAAGACCAAATGCGTGA
TTATGTAA

7）**pIRES-mCerulean-FRB-KRas/N12-KRas/13C(Q61L)-FKBP-mVenus-CAAX**

>Amino acid sequence (<u>61L is underlined</u>)

MVSKGEELFTGVVPILVELDGDVNGHKFSVSGEGEGDATYGKLTLKFICTTGKLPVPWPTLVTT
LTWGVQCFARYPDHMKQHDFFKSAMPEGYVQERTIFFKDDGNYKTRAEVKFEGDTLVNRIELK
GIDFKEDGNILGHKLEYNAISDNVYITADKQKNGIKANFKIRHNIEDGSVQLADHYQQNTPIGDGP
VLLPDNHYLSTQSKLSKDPNEKRDHMVLLEFVTAAGITLGMDELYKGGGSSGGGEMWHEGLE
EASRLYFGERNVKGMFEVLEPLHAMMERGPQTLKETSFNQAYGRDLMEAQEWCRKYMKSGN
VKDLTQAWDLYYHVFRRISKQQISYASRGGGSSGGGELMTEYKLVVVGAG*---***IRES Region***---
MGVGKSALTIQLIQNHFVDEYDPTIEDSYRKQVVIDGETCLLDILDTAG<u>**L**</u>EEYSAMRDQYMRTGE
GFLCVFAINNTKSFEDIHHYREQIKRVKDSEDVPMVLVGNKCDLPSRTVDTKQAQDLARSYGIP
FIETSAKTRQGVDDAFYTLVREIRKHKEKMSKDGGDPNWELVYTARLQGGGSSGGGQISYASR
GGVQVETISPGDGRTFPKRGQTCVVHYTGMLEDGKKFDSSRDRNKPFKFMLGKQEVIRGWEE
GVAQMSVGQRAKLTISPDYAYGATGHPGIIPPHATLVFDVELLKLEGGGSSGGGVSKGEELFTG
VVPILVELDGDVNGHKFSVSGEGEGDATYGKLTLKLICTTGKLPVPWPTLVTTLGYGLQCFARYP
DHMKQHDFFKSAMPEGYVQERTIFFKDDGNYKTRAEVKFEGDTLVNRIELKGIDFKEDGNILGH
KLEYNYNSHNVYITADKQKNGIKANFKIRHNIEDGGVQLADHYQQNTPIGDGPVLLPDNHYLSY
QSKLSKDPNEKRDHMVLLEFVTAAGITLGMDELYKKKKKKKSKTKCVIM*

>DNA sequence

ATGGTGAGCAAGGGCGAGGAGCTGTTCACCGGGGTGGTGCCCATCCTGGTCGAGCTGGA
CGGCGACGTAAACGGCCACAAGTTCAGCGTGTCCGGCGAGGGCGAGGGCGATGCCACCT
ACGGCAAGCTGACCCTGAAGTTCATCTGCACCACCGGCAAGCTGCCCGTGCCCTGGCCCA
CCCTCGTGACCACCCTGACCTGGGGCGTGCAGTGCTTCGCCCGCTACCCCGACCACATGA
AGCAGCACGACTTCTTCAAGTCCGCCATGCCCGAAGGCTACGTCCAGGAGCGCACCATCT
TCTTCAAGGACGACGGCAACTACAAGACCCGCGCCGAGGTGAAGTTCGAGGGCGACACC
CTGGTGAACCGCATCGAGCTGAAGGGCATCGACTTCAAGGAGGACGGCAACATCCTGGGG
CACAAGCTGGAGTACAACGCCATCAGCGACAACGTCTATATCACCGCCGACAAGCAGAAGA
ACGGCATCAAGGCCAACTTCAAGATCCGCCACAACATCGAGGACGGCAGCGTGCAGCTCG
CCGACCACTACCAGCAGAACACCCCCATCGGCGACGGCCCCGTGCTGCTGCCCGACAAC
CACTACCTGAGCACCCAGTCCAAGCTGAGCAAAGACCCCAACGAGAAGCGCGATCACATG
GTCCTGCTGGAGTTCGTGACCGCCGCCGGGATCACTCTCGGCATGGACGAGCTGTACAAG
GGCGGTGGCTCATCTGGCGGAGGTGAGATGTGGCATGAAGGCCTGGAAGAGGCATCTCGT
TTGTACTTTGGGGAAAGGAACGTGAAAGGCATGTTTGAGGTGCTGGAGCCCTTGCATGCTA
TGATGGAACGGGGCCCCCAGACTCTGAAGGAAACATCCTTTAATCAGGCCTATGGTCGAGA
TTTAATGGAGGCCCAAGAGTGGTGCAGGAAGTACATGAAATCAGGGAATGTCAAGGACCTC
ACCCAAGCCTGGGACCTCTATTATCATGTGTTCCGACGAATCTCAAAGCAGCAGATCTCGTA
CGCGTCCCGGGGCGGTGGCTCATCTGGCGGAGGTGAGCTCATGACTGAATATAAACTTGT
GGTAGTTGGAGCTGGTTA---***IRES region***---
ATGGGCGTAGGCAAGAGTGCCTTGACGATACAGCTAATTCAGAATCATTTTGTGGACGAATA
TGATCCAACAATAGAGGATTCCTACAGGAAGCAAGTAGTAATTGATGGAGAAACCTGTCTCT
TGGATATTCTCGACACAGCAGGTCTGGAGGAGTACAGTGCAATGAGGGACCAGTACATGAG
GACTGGGGAGGGCTTTCTTTGTGTATTTGCCATAAATAATACTAAATCATTTGAAGATATTCAC
CATTATAGAGAACAAATTAAAAGAGTTAAGGACTCTGAAGATGTACCTATGGTCCTAGTAGGA
AATAAATGTGATTTGCCTTCCAGAACAGTAGACACAAAACAGGCTCAGGACTTAGCAAGAAG
TTATGGAATTCCTTTTATTGAAACATCAGCAAAGACAAGACAGGGTGTTGATGATGCCTTCTA

TACATTAGTTCGAGAAATTCGAAAACATAAAGAAAAGATGAGCAAAGATGGTGGGGATCCCA
ATTGGGAGCTCGTGTACACGGCGCGCCTGCAGGGAGGTGGCTCATCTGGCGGAGGTCAG
ATCTCGTACGCGTCCCGGGGCGGAGTGCAGGTGGAAACCATCTCCCCAGGAGACGGGCG
CACCTTCCCCAAGCGCGGCCAGACCTGCGTGGTGCACTACACCGGGATGCTTGAAGATGG
AAAGAAATTTGATTCCTCCCGGGACAGAAACAAGCCCTTTAAGTTTATGCTAGGCAAGCAGG
AGGTGATCCGAGGCTGGGAAGAAGGGGTTGCCCAGATGAGTGTGGGTCAGAGAGCCAAA
CTGACTATATCTCCAGATTATGCCTATGGTGCCACTGGGCACCCAGGCATCATCCCACCACA
TGCCACTCTCGTCTTCGATGTGGAGCTTCTAAAACTGGAAGGCGGTGGCTCATCTGGCGGA
GGTGTGAGCAAGGGCGAGGAGCTGTTCACCGGGGTGGTGCCCATCCTGGTCGAGCTGGA
CGGCGACGTAAACGGCCACAAGTTCAGCGTGTCCGGCGAGGGCGAGGGCGATGCCACCT
ACGGCAAGCTGACCCTGAAGCTCATCTGCACCACCGGCAAGCTGCCCGTGCCCTGGCCCA
CCCTCGTGACCACCCTCGGCTACGGCCTGCAGTGCTTCGCCCGCTACCCCGACCACATGA
AGCAGCACGACTTCTTCAAGTCCGCCATGCCCGAAGGCTACGTCCAGGAGCGCACCATCT
TCTTCAAGGACGACGGCAACTACAAGACCCGCGCCGAGGTGAAGTTCGAGGGCGACACC
CTGGTGAACCGCATCGAGCTGAAGGGCATCGACTTCAAGGAGGACGGCAACATCCTGGGG
CACAAGCTGGAGTACAACTACAACAGCCACAACGTCTATATCACCGCCGACAAGCAGAAGA
ACGGCATCAAGGCCAACTTCAAGATCCGCCACAACATCGAGGACGGCGGCGTGCAGCTCG
CCGACCACTACCAGCAGAACACCCCCATCGGCGACGGCCCCGTGCTGCTGCCCGACAAC
CACTACCTGAGCTACCAGTCCAAGCTGAGCAAAGACCCCAACGAGAAGCGCGATCACATG
GTCCTGCTGGAGTTCGTGACCGCCGCCGGGATCACTCTCGGCATGGACGAGCTGTACAAG
AAAAAAAAGAAGAAAAAGAGCAAGACCAAATGCGTGATTATGTAA

8）**pIRES-mCerulean-FRB-KRas/N12-KRas/13C(S17N)-FKBP-mVenus-CAAX**

>Amino acid sequence (17N is underlined)

MVSKGEELFTGVVPILVELDGDVNGHKFSVSGEGEGDATYGKLTLKFICTTGKLPVPWPTLVTT
LTWGVQCFARYPDHMKQHDFFKSAMPEGYVQERTIFFKDDGNYKTRAEVKFEGDTLVNRIELK
GIDFKEDGNILGHKLEYNAISDNVYITADKQKNGIKANFKIRHNIEDGSVQLADHYQQNTPIGDGP
VLLPDNHYLSTQSKLSKDPNEKRDHMVLLEFVTAAGITLGMDELYKGGGSSGGGEMWHEGLE
EASRLYFGERNVKGMFEVLEPLHAMMERGPQTLKETSFNQAYGRDLMEAQEWCRKYMKSGN
VKDLTQAWDLYYHVFRRISKQQISYASRGGGSSGGGELMTEYKLVVVGAG∗---***IRES Region***---
MGVGK**N**ALTIQLIQNHFVDEYDPTIEDSYRKQVVIDGETCLLDILDTAGQEEYSAMRDQYMRTG
EGFLCVFAINNTKSFEDIHHYREQIKRVKDSEDVPMVLVGNKCDLPSRTVDTKQAQDLARSYGI
PFIETSAKTRQGVDDAFYTLVREIRKHKEKMSKDGGDPNWELVYTARLQGGGSSGGGQISYAS
RGGVQVETISPGDGRTFPKRGQTCVVHYTGMLEDGKKFDSSRDRNKPFKFMLGKQEVIRGWE
EGVAQMSVGQRAKLTISPDYAYGATGHPGIIPPHATLVFDVELLKLEGGGSSGGGVSKGEELFT
GVVPILVELDGDVNGHKFSVSGEGEGDATYGKLTLKLICTTGKLPVPWPTLVTTLGYGLQCFAR
YPDHMKQHDFFKSAMPEGYVQERTIFFKDDGNYKTRAEVKFEGDTLVNRIELKGIDFKEDGNIL
GHKLEYNYNSHNVYITADKQKNGIKANFKIRHNIEDGGVQLADHYQQNTPIGDGPVLLPDNHYL
SYQSKLSKDPNEKRDHMVLLEFVTAAGITLGMDELYKKKKKKKSKTKCVIM∗

>DNA sequence

ATGGTGAGCAAGGGCGAGGAGCTGTTCACCGGGGTGGTGCCCATCCTGGTCGAGCTGGA
CGGCGACGTAAACGGCCACAAGTTCAGCGTGTCCGGCGAGGGCGAGGGCGATGCCACCT
ACGGCAAGCTGACCCTGAAGTTCATCTGCACCACCGGCAAGCTGCCCGTGCCCTGGCCCA
CCCTCGTGACCACCCTGACCTGGGGCGTGCAGTGCTTCGCCCGCTACCCCGACCACATGA
AGCAGCACGACTTCTTCAAGTCCGCCATGCCCGAAGGCTACGTCCAGGAGCGCACCATCT
TCTTCAAGGACGACGGCAACTACAAGACCCGCGCCGAGGTGAAGTTCGAGGGCGACACC
CTGGTGAACCGCATCGAGCTGAAGGGCATCGACTTCAAGGAGGACGGCAACATCCTGGGG
CACAAGCTGGAGTACAACGCCATCAGCGACAACGTCTATATCACCGCCGACAAGCAGAAGA
ACGGCATCAAGGCCAACTTCAAGATCCGCCACAACATCGAGGACGGCAGCGTGCAGCTCG
CCGACCACTACCAGCAGAACACCCCCATCGGCGACGGCCCCGTGCTGCTGCCCGACAAC
CACTACCTGAGCACCCAGTCCAAGCTGAGCAAAGACCCCAACGAGAAGCGCGATCACATG
GTCCTGCTGGAGTTCGTGACCGCCGCCGGGATCACTCTCGGCATGGACGAGCTGTACAAG
GGCGGTGGCTCATCTGGCGGAGGTGAGATGTGGCATGAAGGCCTGGAAGAGGCATCTCGT
TTGTACTTTGGGGAAAGGAACGTGAAAGGCATGTTTGAGGTGCTGGAGCCCTTGCATGCTA
TGATGGAACGGGGCCCCCAGACTCTGAAGGAAACATCCTTTAATCAGGCCTATGGTCGAGA
TTTAATGGAGGCCCAAGAGTGGTGCAGGAAGTACATGAAATCAGGGAATGTCAAGGACCTC
ACCCAAGCCTGGGACCTCTATTATCATGTGTTCCGACGAATCTCAAAGCAGCAGATCTCGTA
CGCGTCCCGGGGCGGTGGCTCATCTGGCGGAGGTGAGCTCATGACTGAATATAAACTTGT
GGTAGTTGGAGCTGGTTA---***IRES region***---
ATGGGCGTAGGCAAGAATGCCTTGACGATACAGCTAATTCAGAATCATTTTGTGGACGAATAT
GATCCAACAATAGAGGATTCCTACAGGAAGCAAGTAGTAATTGATGGAGAAACCTGTCTCTT
GGATATTCTCGACACAGCAGGTCAAGAGGAGTACAGTGCAATGAGGGACCAGTACATGAGG
ACTGGGGAGGGCTTTCTTTGTGTATTTGCCATAAATAATACTAAATCATTTGAAGATATTCACC
ATTATAGAGAACAAATTAAAAGAGTTAAGGACTCTGAAGATGTACCTATGGTCCTAGTAGGAA
ATAAATGTGATTTGCCTTCCAGAACAGTAGACACAAAACAGGCTCAGGACTTAGCAAGAAGT
TATGGAATTCCTTTTATTGAAACATCAGCAAAGACAAGACAGGGTGTTGATGATGCCTTCTAT

ACATTAGTTCGAGAAATTCGAAAACATAAAGAAAAGATGAGCAAAGATGGTGGGGATCCCAA
TTGGGAGCTCGTGTACACGGCGCGCCTGCAGGGAGGTGGCTCATCTGGCGGAGGTCAGA
TCTCGTACGCGTCCCGGGGCGGAGTGCAGGTGGAAACCATCTCCCCAGGAGACGGGCGC
ACCTTCCCCAAGCGCGGCCAGACCTGCGTGGTGCACTACACCGGGATGCTTGAAGATGGA
AAGAAATTTGATTCCTCCCGGGACAGAAACAAGCCCTTTAAGTTTATGCTAGGCAAGCAGGA
GGTGATCCGAGGCTGGGAAGAAGGGGTTGCCCAGATGAGTGTGGGTCAGAGAGCCAAAC
TGACTATATCTCCAGATTATGCCTATGGTGCCACTGGGCACCCAGGCATCATCCCACCACAT
GCCACTCTCGTCTTCGATGTGGAGCTTCTAAAACTGGAAGGCGGTGGCTCATCTGGCGGA
GGTGTGAGCAAGGGCGAGGAGCTGTTCACCGGGGTGGTGCCCATCCTGGTCGAGCTGGA
CGGCGACGTAAACGGCCACAAGTTCAGCGTGTCCGGCGAGGGCGAGGGCGATGCCACCT
ACGGCAAGCTGACCCTGAAGCTCATCTGCACCACCGGCAAGCTGCCCGTGCCCTGGCCCA
CCCTCGTGACCACCCTCGGCTACGGCCTGCAGTGCTTCGCCCGCTACCCCGACCACATGA
AGCAGCACGACTTCTTCAAGTCCGCCATGCCCGAAGGCTACGTCCAGGAGCGCACCATCT
TCTTCAAGGACGACGGCAACTACAAGACCCGCGCCGAGGTGAAGTTCGAGGGCGACACC
CTGGTGAACCGCATCGAGCTGAAGGGCATCGACTTCAAGGAGGACGGCAACATCCTGGGG
CACAAGCTGGAGTACAACTACAACAGCCACAACGTCTATATCACCGCCGACAAGCAGAAGA
ACGGCATCAAGGCCAACTTCAAGATCCGCCACAACATCGAGGACGGCGGCGTGCAGCTCG
CCGACCACTACCAGCAGAACACCCCCATCGGCGACGGCCCCGTGCTGCTGCCCGACAAC
CACTACCTGAGCTACCAGTCCAAGCTGAGCAAAGACCCCAACGAGAAGCGCGATCACATG
GTCCTGCTGGAGTTCGTGACCGCCGCCGGGATCACTCTCGGCATGGACGAGCTGTACAAG
AAAAAAAAGAAGAAAAAGAGCAAGACCAAATGCGTGATTATGTAA

9）**pIRES-mCerulean-GID1-KRas/N12-KRas/13C(Q61L)-GAI$_{1\text{-}92}$-mVenus-CAAX**

>Amino acid sequence (61L is underlined)

MVSKGEELFTGVVPILVELDGDVNGHKFSVSGEGEGDATYGKLTLKFICTTGKLPVPWPTLVTT
LTWGVQCFARYPDHMKQHDFFKSAMPEGYVQERTIFFKDDGNYKTRAEVKFEGDTLVNRIELK
GIDFKEDGNILGHKLEYNAISDNVYITADKQKNGIKANFKIRHNIEDGSVQLADHYQQNTPIGDGP
VLLPDNHYLSTQSKLSKDPNEKRDHMVLLEFVTAAGITLGMDELYKGGGSSGGGMAASDEVNL
IESRTVVPLNTWVLISNFKVAYNILRRPDGTFNRHLAEYLDRKVTANANPVDGVFSFDVLIDRRIN
LLSRVYRPAYADQEQPPSILDLEKPVDGDIVPVILFFHGGSFAHSSANSAIYDTLCRRLVGLCKCV
VVSVNYRRAPENPYPCAYDDGWIALNWVNSRSWLKSKKDSKVHIFLAGDSSGGNIAHNVALRA
GESGIDVLGNILLNPMFGGNERTESEKSLDGKYFVTVRDRDWYWKAFLPEGEDREHPACNPF
SPRGKSLEGVSFPKSLVVVAGLDLIRDWQLAYAEGLKKAGQEVKLMHLEKATVGFYLLPNNNHF
HNVMDEISAFVNAECQISYASRGGGSSGGGELMTEYKLVVVGAG∗---***IRES Region***---
MGVGKSALTIQLIQNHFVDEYDPTIEDSYRKQVVIDGETCLLDILDTAG**L**EEYSAMRDQYMRTGE
GFLCVFAINNTKSFEDIHHYREQIKRVKDSEDVPMVLVGNKCDLPSRTVDTKQAQDLARSYGIP
FIETSAKTRQGVDDAFYTLVREIRKHKEKMSKDGGDPNWELVYTARLQGGGSSGGGQISYASR
GMKRDHHHHHHQDKKTMMMNEEDDGNGMDELLAVLGYKVRSSEMADVAQKLEQLEVMMSN
VQEDDLSQLATETVHYNPAELYTWLDSMLTDLNGGGSSGGGVSKGEELFTGVVPILVELDGDV
NGHKFSVSGEGEGDATYGKLTLKLICTTGKLPVPWPTLVTTLGYGLQCFARYPDHMKQHDFFK
SAMPEGYVQERTIFFKDDGNYKTRAEVKFEGDTLVNRIELKGIDFKEDGNILGHKLEYNYNSHN
VYITADKQKNGIKANFKIRHNIEDGGVQLADHYQQNTPIGDGPVLLPDNHYLSYQSKLSKDPNE
KRDHMVLLEFVTAAGITLGMDELYKKKKKKKSKTKCVIM∗

>DNA sequence

ATGGTGAGCAAGGGCGAGGAGCTGTTCACCGGGGTGGTGCCCATCCTGGTCGAGCTGGA
CGGCGACGTAAACGGCCACAAGTTCAGCGTGTCCGGCGAGGGCGAGGGCGATGCCACCT
ACGGCAAGCTGACCCTGAAGTTCATCTGCACCACCGGCAAGCTGCCCGTGCCCTGGCCCA
CCCTCGTGACCACCCTGACCTGGGGCGTGCAGTGCTTCGCCCGCTACCCCGACCACATGA
AGCAGCACGACTTCTTCAAGTCCGCCATGCCCGAAGGCTACGTCCAGGAGCGCACCATCT
TCTTCAAGGACGACGGCAACTACAAGACCCGCGCCGAGGTGAAGTTCGAGGGCGACACC
CTGGTGAACCGCATCGAGCTGAAGGGCATCGACTTCAAGGAGGACGGCAACATCCTGGGG
CACAAGCTGGAGTACAACGCCATCAGCGACAACGTCTATATCACCGCCGACAAGCAGAAGA
ACGGCATCAAGGCCAACTTCAAGATCCGCCACAACATCGAGGACGGCAGCGTGCAGCTCG
CCGACCACTACCAGCAGAACACCCCCATCGGCGACGGCCCCGTGCTGCTGCCCGACAAC
CACTACCTGAGCACCCAGTCCAAGCTGAGCAAAGACCCCAACGAGAAGCGCGATCACATG
GTCCTGCTGGAGTTCGTGACCGCCGCCGGGATCACTCTCGGCATGGACGAGCTGTACAAG
GGCGGTGGCTCATCTGGCGGAGGTATGGCTGCGAGCGATGAAGTTAATCTTATTGAGAGCA
GAACAGTGGTTCCTCTCAATACATGGGTTTTAATATCCAACTTCAAAGTAGCCTACAATATCCT
TCGTCGCCCTGATGGAACCTTTAACCGACACTTAGCTGAGTATCTAGACCGTAAAGTCACTG
CAAACGCCAATCCGGTTGATGGGGTTTTCTCGTTCGATGTCTTGATTGATCGCAGGATCAAT
CTTCTAAGCAGAGTCTATAGACCAGCTTATGCAGATCAAGAGCAACCTCCTAGTATTTTAGAT
CTCGAGAAGCCTGTTGATGGCGACATTGTCCCTGTTATATTGTTCTTCCATGGAGGTAGCTT
TGCTCATTCTTCTGCAAACAGTGCCATCTACGATACTCTTTGTCGCAGGCTTGTTGGTTTGT
GCAAGTGTGTTGTTGTCTCTGTGAATTATCGGCGTGCACCAGAGAATCCATACCCTTGTGCT
TATGATGATGGTTGGATTGCTCTTAATTGGGTTAACTCGAGATCTTGGCTTAAATCCAAGAAA
GACTCAAAGGTCCATATTTTCTTGGCTGGTGATAGCTCTGGAGGTAACATCGCGCATAATGT

GGCTTTAAGAGCGGGTGAATCGGGAATCGATGTTTTGGGGAACATTCTGCTGAATCCTATGT
TTGGTGGGAATGAGAGAACGGAGTCTGAGAAAAGTTTGGATGGGAAATACTTTGTGACGGT
TAGAGACCGCGATTGGTACTGGAAAGCGTTTTTACCCGAGGGAGAAGATAGAGAGCATCCA
GCGTGTAATCCGTTTAGCCCGAGAGGGAAAAGCTTAGAAGGAGTGAGTTTCCCCAAGAGTC
TTGTGGTTGTCGCGGGTTTGGATTTGATTAGAGATTGGCAGTTGGCATACGCGGAAGGGCT
CAAGAAAGCGGGTCAAGAGGTTAAGCTTATGCATTTAGAGAAAGCAACTGTTGGGTTTTACC
TCTTGCCTAATAACAATCATTTCCATAATGTTATGGATGAGATTTCGGCGTTTGTAAACGCGGA
ATGTCAGATCTCGTACGCGTCCCGGGGCGGTGGCTCATCTGGCGGAGGTGAGCTCATGAC
TGAATATAAACTTGTGGTAGTTGGAGCTGGTTA---***IRES region***---
ATGGGCGTAGGCAAGAGTGCCTTGACGATACAGCTAATTCAGAATCATTTTGTGGACGAATA
TGATCCAACAATAGAGGATTCCTACAGGAAGCAAGTAGTAATTGATGGAGAAACCTGTCTCT
TGGATATTCTCGACACAGCAGGTCTGGAGGAGTACAGTGCAATGAGGGACCAGTACATGAG
GACTGGGGAGGGCTTTCTTTGTGTATTTGCCATAAATAATACTAAATCATTTGAAGATATTCAC
CATTATAGAGAACAAATTAAAAGAGTTAAGGACTCTGAAGATGTACCTATGGTCCTAGTAGGA
AATAAATGTGATTTGCCTTCCAGAACAGTAGACACAAAACAGGCTCAGGACTTAGCAAGAAG
TTATGGAATTCCTTTTATTGAAACATCAGCAAAGACAAGACAGGGTGTTGATGATGCCTTCTA
TACATTAGTTCGAGAAATTCGAAAACATAAAGAAAAGATGAGCAAAGATGGTGGGGATCCCA
ATTGGGAGCTCGTGTACACGGCGCGCCTGCAGGGAGGTGGCTCATCTGGCGGAGGTCAG
ATCTCGTACGCGTCCCGGGGCATGAAGAGAGATCATCATCATCATCATCATCAAGATAAGAA
GACTATGATGATGAATGAAGAAGACGACGGTAACGGCATGGATGAGCTTCTAGCTGTTCTTG
GTTACAAGGTTAGGTCATCCGAAATGGCTGATGTTGCTCAGAAACTCGAGCAGCTTGAAGTT
ATGATGTCTAATGTTCAAGAAGACGATCTTTCTCAACTCGCTACTGAGACTGTTCACTATAAT
CCGGCGGAGCTTTACACGTGGCTTGATTCTATGCTCACCGACCTTAATGGCGGTGGCTCAT
CTGGCGGAGGTGTGAGCAAGGGCGAGGAGCTGTTCACCGGGGTGGTGCCCATCCTGGTC
GAGCTGGACGGCGACGTAAACGGCCACAAGTTCAGCGTGTCCGGCGAGGGCGAGGGCGA
TGCCACCTACGGCAAGCTGACCCTGAAGCTCATCTGCACCACCGGCAAGCTGCCCGTGCC
CTGGCCCACCCTCGTGACCACCCTCGGCTACGGCCTGCAGTGCTTCGCCCGCTACCCCGA
CCACATGAAGCAGCACGACTTCTTCAAGTCCGCCATGCCCGAAGGCTACGTCCAGGAGCG
CACCATCTTCTTCAAGGACGACGGCAACTACAAGACCCGCGCCGAGGTGAAGTTCGAGGG
CGACACCCTGGTGAACCGCATCGAGCTGAAGGGCATCGACTTCAAGGAGGACGGCAACAT
CCTGGGGCACAAGCTGGAGTACAACTACAACAGCCACAACGTCTATATCACCGCCGACAAG
CAGAAGAACGGCATCAAGGCCAACTTCAAGATCCGCCACAACATCGAGGACGGCGGCGTG
CAGCTCGCCGACCACTACCAGCAGAACACCCCCATCGGCGACGGCCCCGTGCTGCTGCC
CGACAACCACTACCTGAGCTACCAGTCCAAGCTGAGCAAAGACCCCAACGAGAAGCGCGA
TCACATGGTCCTGCTGGAGTTCGTGACCGCCGCCGGGATCACTCTCGGCATGGACGAGCT
GTACAAGAAAAAAAAGAAGAAAAAGAGCAAGACCAAATGCGTGATTATGTAA

10） **pIRES-mCerulean-GID1-KRas/N12-KRas/13C(S17N)-GAI$_{1-92}$-mVenus-CAAX**

>Amino acid sequence (17N is underlined)

MVSKGEELFTGVVPILVELDGDVNGHKFSVSGEGEGDATYGKLTLKFICTTGKLPVPWPTLVTT
LTWGVQCFARYPDHMKQHDFFKSAMPEGYVQERTIFFKDDGNYKTRAEVKFEGDTLVNRIELK
GIDFKEDGNILGHKLEYNAISDNVYITADKQKNGIKANFKIRHNIEDGSVQLADHYQQNTPIGDGP
VLLPDNHYLSTQSKLSKDPNEKRDHMVLLEFVTAAGITLGMDELYKGGGSSGGGMAASDEVNL
IESRTVVPLNTWVLISNFKVAYNILRRPDGTFNRHLAEYLDRKVTANANPVDGVFSFDVLIDRRIN
LLSRVYRPAYADQEQPPSILDLEKPVDGDIVPVILFFHGGSFAHSSANSAIYDTLCRRLVGLCKCV
VVSVNYRRAPENPYPCAYDDGWIALNWVNSRSWLKSKKDSKVHIFLAGDSSGGNIAHNVALRA
GESGIDVLGNILLNPMFGGNERTESEKSLDGKYFVTVRDRDWYWKAFLPEGEDREHPACNPF
SPRGKSLEGVSFPKSLVVVAGLDLIRDWQLAYAEGLKKAGQEVKLMHLEKATVGFYLLPNNNHF
HNVMDEISAFVNAECQISYASRGGGSSGGGELMTEYKLVVVGAG*---***IRES Region***---
MGVGK**N**ALTIQLIQNHFVDEYDPTIEDSYRKQVVIDGETCLLDILDTAGQEEYSAMRDQYMRTG
EGFLCVFAINNTKSFEDIHHYREQIKRVKDSEDVPMVLVGNKCDLPSRTVDTKQAQDLARSYGI
PFIETSAKTRQGVDDAFYTLVREIRKHKEKMSKDGGDPNWELVYTARLQGGGSSGGGQISYAS
RGMKRDHHHHHHQDKKTMMMNEEDDGNGMDELLAVLGYKVRSSEMADVAQKLEQLEVMMS
NVQEDDLSQLATETVHYNPAELYTWLDSMLTDLNGGGSSGGGVSKGEELFTGVVPILVELDGD
VNGHKFSVSGEGEGDATYGKLTLKLICTTGKLPVPWPTLVTTLGYGLQCFARYPDHMKQHDFF
KSAMPEGYVQERTIFFKDDGNYKTRAEVKFEGDTLVNRIELKGIDFKEDGNILGHKLEYNYNSH
NVYITADKQKNGIKANFKIRHNIEDGGVQLADHYQQNTPIGDGPVLLPDNHYLSYQSKLSKDPN
EKRDHMVLLEFVTAAGITLGMDELYKKKKKKKSKTKCVIM*

>DNA sequence

ATGGTGAGCAAGGGCGAGGAGCTGTTCACCGGGGTGGTGCCCATCCTGGTCGAGCTGGA
CGGCGACGTAAACGGCCACAAGTTCAGCGTGTCCGGCGAGGGCGAGGGCGATGCCACCT
ACGGCAAGCTGACCCTGAAGTTCATCTGCACCACCGGCAAGCTGCCCGTGCCCTGGCCCA
CCCTCGTGACCACCCTGACCTGGGGCGTGCAGTGCTTCGCCCGCTACCCCGACCACATGA
AGCAGCACGACTTCTTCAAGTCCGCCATGCCCGAAGGCTACGTCCAGGAGCGCACCATCT
TCTTCAAGGACGACGGCAACTACAAGACCCGCGCCGAGGTGAAGTTCGAGGGCGACACC
CTGGTGAACCGCATCGAGCTGAAGGGCATCGACTTCAAGGAGGACGGCAACATCCTGGGG
CACAAGCTGGAGTACAACGCCATCAGCGACAACGTCTATATCACCGCCGACAAGCAGAAGA
ACGGCATCAAGGCCAACTTCAAGATCCGCCACAACATCGAGGACGGCAGCGTGCAGCTCG
CCGACCACTACCAGCAGAACACCCCCATCGGCGACGGCCCCGTGCTGCTGCCCGACAAC
CACTACCTGAGCACCCAGTCCAAGCTGAGCAAAGACCCCAACGAGAAGCGCGATCACATG
GTCCTGCTGGAGTTCGTGACCGCCGCCGGGATCACTCTCGGCATGGACGAGCTGTACAAG
GGCGGTGGCTCATCTGGCGGAGGTATGGCTGCGAGCGATGAAGTTAATCTTATTGAGAGCA
GAACAGTGGTTCCTCTCAATACATGGGTTTTAATATCCAACTTCAAAGTAGCCTACAATATCCT
TCGTCGCCCTGATGGAACCTTTAACCGACACTTAGCTGAGTATCTAGACCGTAAAGTCACTG
CAAACGCCAATCCGGTTGATGGGGTTTTCTCGTTCGATGTCTTGATTGATCGCAGGATCAAT
CTTCTAAGCAGAGTCTATAGACCAGCTTATGCAGATCAAGAGCAACCTCCTAGTATTTTAGAT
CTCGAGAAGCCTGTTGATGGCGACATTGTCCCTGTTATATTGTTCTTCCATGGAGGTAGCTT
TGCTCATTCTTCTGCAAACAGTGCCATCTACGATACTCTTTGTCGCAGGCTTGTTGGTTTGT
GCAAGTGTGTTGTTGTCTCTGTGAATTATCGGCGTGCACCAGAGAATCCATACCCTTGTGCT
TATGATGATGGTTGGATTGCTCTTAATTGGGTTAACTCGAGATCTTGGCTTAAATCCAAGAAA
GACTCAAAGGTCCATATTTTCTTGGCTGGTGATAGCTCTGGAGGTAACATCGCGCATAATGT

GGCTTTAAGAGCGGGTGAATCGGGAATCGATGTTTTGGGGAACATTCTGCTGAATCCTATGT
TTGGTGGGAATGAGAGAACGGAGTCTGAGAAAAGTTTGGATGGGAAATACTTTGTGACGGT
TAGAGACCGCGATTGGTACTGGAAAGCGTTTTTACCCGAGGGAGAAGATAGAGAGCATCCA
GCGTGTAATCCGTTTAGCCCGAGAGGGAAAAGCTTAGAAGGAGTGAGTTTCCCCAAGAGTC
TTGTGGTTGTCGCGGGTTTGGATTTGATTAGAGATTGGCAGTTGGCATACGCGGAAGGGCT
CAAGAAAGCGGGTCAAGAGGTTAAGCTTATGCATTTAGAGAAAGCAACTGTTGGGTTTTACC
TCTTGCCTAATAACAATCATTTCCATAATGTTATGGATGAGATTTCGGCGTTTGTAAACGCGGA
ATGTCAGATCTCGTACGCGTCCCGGGGCGGTGGCTCATCTGGCGGAGGTGAGCTCATGAC
TGAATATAAACTTGTGGTAGTTGGAGCTGGTTA---***IRES region***---
ATGGGCGTAGGCAAGAATGCCTTGACGATACAGCTAATTCAGAATCATTTTGTGGACGAATAT
GATCCAACAATAGAGGATTCCTACAGGAAGCAAGTAGTAATTGATGGAGAAACCTGTCTCTT
GGATATTCTCGACACAGCAGGTCAAGAGGAGTACAGTGCAATGAGGGACCAGTACATGAGG
ACTGGGGAGGGCTTTCTTTGTGTATTTGCCATAAATAATACTAAATCATTTGAAGATATTCACC
ATTATAGAGAACAAATTAAAAGAGTTAAGGACTCTGAAGATGTACCTATGGTCCTAGTAGGAA
ATAAATGTGATTTGCCTTCCAGAACAGTAGACACAAAACAGGCTCAGGACTTAGCAAGAAGT
TATGGAATTCCTTTTATTGAAACATCAGCAAAGACAAGACAGGGTGTTGATGATGCCTTCTAT
ACATTAGTTCGAGAAATTCGAAAACATAAAGAAAAGATGAGCAAAGATGGTGGGGATCCCAA
TTGGGAGCTCGTGTACACGGCGCGCCTGCAGGGAGGTGGCTCATCTGGCGGAGGTCAGA
TCTCGTACGCGTCCCGGGGCATGAAGAGAGATCATCATCATCATCATCATCAAGATAAGAAG
ACTATGATGATGAATGAAGAAGACGACGGTAACGGCATGGATGAGCTTCTAGCTGTTCTTGG
TTACAAGGTTAGGTCATCCGAAATGGCTGATGTTGCTCAGAAACTCGAGCAGCTTGAAGTTA
TGATGTCTAATGTTCAAGAAGACGATCTTTCTCAACTCGCTACTGAGACTGTTCACTATAATC
CGGCGGAGCTTTACACGTGGCTTGATTCTATGCTCACCGACCTTAATGGCGGTGGCTCATC
TGGCGGAGGTGTGAGCAAGGGCGAGGAGCTGTTCACCGGGGTGGTGCCCATCCTGGTCG
AGCTGGACGGCGACGTAAACGGCCACAAGTTCAGCGTGTCCGGCGAGGGCGAGGGCGAT
GCCACCTACGGCAAGCTGACCCTGAAGCTCATCTGCACCACCGGCAAGCTGCCCGTGCCC
TGGCCCACCCTCGTGACCACCCTCGGCTACGGCCTGCAGTGCTTCGCCCGCTACCCCGAC
CACATGAAGCAGCACGACTTCTTCAAGTCCGCCATGCCCGAAGGCTACGTCCAGGAGCGC
ACCATCTTCTTCAAGGACGACGGCAACTACAAGACCCGCGCCGAGGTGAAGTTCGAGGGC
GACACCCTGGTGAACCGCATCGAGCTGAAGGGCATCGACTTCAAGGAGGACGGCAACATC
CTGGGGCACAAGCTGGAGTACAACTACAACAGCCACAACGTCTATATCACCGCCGACAAGC
AGAAGAACGGCATCAAGGCCAACTTCAAGATCCGCCACAACATCGAGGACGGCGGCGTGC
AGCTCGCCGACCACTACCAGCAGAACACCCCCATCGGCGACGGCCCCGTGCTGCTGCCC
GACAACCACTACCTGAGCTACCAGTCCAAGCTGAGCAAAGACCCCAACGAGAAGCGCGAT
CACATGGTCCTGCTGGAGTTCGTGACCGCCGCCGGGATCACTCTCGGCATGGACGAGCTG
TACAAGAAAAAAAAGAAGAAAAAGAGCAAGACCAAATGCGTGATTATGTAA

**Table S3. Sequences for bacterial expression constructs used in this study.**

1）**pMAL-MBP-Cdc42-Q61L**

>Amino acid sequence

MKIEEGKLVIWINGDKGYNGLAEVGKKFEKDTGIKVTVEHPDKLEEKFPQVAATGDGPDIIFWAH
DRFGGYAQSGLLAEITPDKAFQDKLYPFTWDAVRYNGKLIAYPIAVEALSLIYNKDLLPNPPKTW
EEIPALDKELKAKGKSALMFNLQEPYFTWPLIAADGGYAFKYENGKYDIKDVGVDNAGAKAGLT
FLVDLIKNKHMNADTDYSIAEAAFNKGETAMTINGPWAWSNIDTSKVNYGVTVLPTFKGQPSKP
FVGVLSAGINAASPNKELAKEFLENYLLTDEGLEAVNKDKPLGAVALKSYEEELVKDPRIAATME
NAQKGEIMPNIPQMSAFWYAVRTAVINAASGRQTVDEALKDAQTNSSSNNNNNNNNNNLGIEG
RISHMQTIKCVVVGDGAVGKTCLLISYTTNKFPSEYVPTVFDNYAVTVMIGGEPYTLGLFDTAGL
EDYDRLRPLSYPQTDVFLVCFSVVSPSSFENVKEKWVPEITHHCPKTPFLLVGTQIDLRDDPSTI
EKLAKNKQKPITPETAEKLARDLKAVKYVECSALTQRGLKNVFDEAILAALEPPETQP*

>DNA sequence

ATGAAAATCGAAGAAGGTAAACTGGTAATCTGGATTAACGGCGATAAAGGCTATAACGGTCT
CGCTGAAGTCGGTAAGAAATTCGAGAAAGATACCGGAATTAAAGTCACCGTTGAGCATCCG
GATAAACTGGAAGAGAAATTCCCACAGGTTGCGGCAACTGGCGATGGCCCTGACATTATCT
TCTGGGCACACGACCGCTTTGGTGGCTACGCTCAATCTGGCCTGTTGGCTGAAATCACCCC
GGACAAAGCGTTCCAGGACAAGCTGTATCCGTTTACCTGGGATGCCGTACGTTACAACGGC
AAGCTGATTGCTTACCCGATCGCTGTTGAAGCGTTATCGCTGATTTATAACAAAGATCTGCTG
CCGAACCCGCCAAAAACCTGGGAAGAGATCCCGGCGCTGGATAAAGAACTGAAAGCGAAA
GGTAAGAGCGCGCTGATGTTCAACCTGCAAGAACCGTACTTCACCTGGCCGCTGATTGCTG
CTGACGGGGGTTATGCGTTCAAGTATGAAAACGGCAAGTACGACATTAAAGACGTGGGCGT
GGATAACGCTGGCGCGAAAGCGGGTCTGACCTTCCTGGTTGACCTGATTAAAAACAAACAC
ATGAATGCAGACACCGATTACTCCATCGCAGAAGCTGCCTTTAATAAAGGCGAAACAGCGAT
GACCATCAACGGCCCGTGGGCATGGTCCAACATCGACACCAGCAAAGTGAATTATGGTGTA
ACGGTACTGCCGACCTTCAAGGGTCAACCATCCAAACCGTTCGTTGGCGTGCTGAGCGCA
GGTATTAACGCCGCCAGTCCGAACAAAGAGCTGGCAAAAGAGTTCCTCGAAAACTATCTGC
TGACTGATGAAGGTCTGGAAGCGGTTAATAAAGACAAACCGCTGGGTGCCGTAGCGCTGAA
GTCTTACGAGGAAGAGTTGGTGAAAGATCCGCGTATTGCCGCCACTATGGAAAACGCCCAG
AAAGGTGAAATCATGCCGAACATCCCGCAGATGTCCGCTTTCTGGTATGCCGTGCGTACTG
CGGTGATCAACGCCGCCAGCGGTCGTCAGACTGTCGATGAAGCCCTGAAAGACGCGCAGA
CTAATTCGAGCTCGAACAACAACAACAATAACAATAACAACAACCTCGGGATCGAGGGAAGG
ATTTCACATATGCAGACAATTAAGTGTGTTGTTGTGGGCGATGGTGCTGTTGGTAAAACATGT
CTCCTGATATCCTACACAACAAACAAATTTCCATCGGAGTATGTACCGACTGTTTTTGACAAC
TATGCAGTCACAGTTATGATTGGTGGAGAACCATATACTCTTGGACTTTTTGATACTGCAGGG
CTAGAGGATTATGACAGATTACGACCGCTGAGTTATCCACAAACAGATGTATTTCTAGTCTGT
TTTTCAGTGGTCTCTCCATCTTCATTTGAAAACGTGAAAGAAAAGTGGGTGCCTGAGATAAC
TCACCACTGTCCAAAGACTCCTTTCTTGCTTGTTGGGACTCAAATTGATCTCAGAGATGACC
CCTCTACTATTGAGAAACTTGCCAAGAACAAACAGAAGCCTATCACTCCAGAGACTGCTGAA
AAGCTGGCCCGTGACCTGAAGGCTGTCAAGTATGTGGAGTGTTCTGCACTTACACAGAGAG
GTCTGAAGAATGTGTTTGATGAGGCTATCCTAGCTGCCCTCGAGCCTCCGGAAACTCAACC
CTAA

2）**pMAL-MBP-Cdc42-T17N**

>Amino acid sequence

MKIEEGKLVIWINGDKGYNGLAEVGKKFEKDTGIKVTVEHPDKLEEKFPQVAATGDGPDIIFWAH
DRFGGYAQSGLLAEITPDKAFQDKLYPFTWDAVRYNGKLIAYPIAVEALSLIYNKDLLPNPPKTW
EEIPALDKELKAKGKSALMFNLQEPYFTWPLIAADGGYAFKYENGKYDIKDVGVDNAGAKAGLT
FLVDLIKNKHMNADTDYSIAEAAFNKGETAMTINGPWAWSNIDTSKVNYGVTVLPTFKGQPSKP
FVGVLSAGINAASPNKELAKEFLENYLLTDEGLEAVNKDKPLGAVALKSYEEELVKDPRIAATME
NAQKGEIMPNIPQMSAFWYAVRTAVINAASGRQTVDEALKDAQTNSSSNNNNNNNNNNLGIEG
RISHMQTIKCVVVGDGAVGKNCLLISYTTNKFPSEYVPTVFDNYAVTVMIGGEPYTLGLFDTAGQ
EDYDRLRPLSYPQTDVFLVCFSVVSPSSFENVKEKWVPEITHHCPKTPFLLVGTQIDLRDDPSTI
EKLAKNKQKPITPETAEKLARDLKAVKYVECSALTQRGLKNVFDEAILAALEPPETQP*

>DNA sequence

ATGAAAATCGAAGAAGGTAAACTGGTAATCTGGATTAACGGCGATAAAGGCTATAACGGTCT
CGCTGAAGTCGGTAAGAAATTCGAGAAAGATACCGGAATTAAAGTCACCGTTGAGCATCCG
GATAAACTGGAAGAGAAATTCCCACAGGTTGCGGCAACTGGCGATGGCCCTGACATTATCT
TCTGGGCACACGACCGCTTTGGTGGCTACGCTCAATCTGGCCTGTTGGCTGAAATCACCCC
GGACAAAGCGTTCCAGGACAAGCTGTATCCGTTTACCTGGGATGCCGTACGTTACAACGGC
AAGCTGATTGCTTACCCGATCGCTGTTGAAGCGTTATCGCTGATTTATAACAAAGATCTGCTG
CCGAACCCGCCAAAAACCTGGGAAGAGATCCCGGCGCTGGATAAAGAACTGAAAGCGAAA
GGTAAGAGCGCGCTGATGTTCAACCTGCAAGAACCGTACTTCACCTGGCCGCTGATTGCTG
CTGACGGGGGTTATGCGTTCAAGTATGAAAACGGCAAGTACGACATTAAAGACGTGGGCGT
GGATAACGCTGGCGCGAAAGCGGGTCTGACCTTCCTGGTTGACCTGATTAAAAACAAACAC
ATGAATGCAGACACCGATTACTCCATCGCAGAAGCTGCCTTTAATAAAGGCGAAACAGCGAT
GACCATCAACGGCCCGTGGGCATGGTCCAACATCGACACCAGCAAAGTGAATTATGGTGTA
ACGGTACTGCCGACCTTCAAGGGTCAACCATCCAAACCGTTCGTTGGCGTGCTGAGCGCA
GGTATTAACGCCGCCAGTCCGAACAAAGAGCTGGCAAAAGAGTTCCTCGAAAACTATCTGC
TGACTGATGAAGGTCTGGAAGCGGTTAATAAAGACAAACCGCTGGGTGCCGTAGCGCTGAA
GTCTTACGAGGAAGAGTTGGTGAAAGATCCGCGTATTGCCGCCACTATGGAAAACGCCCAG
AAAGGTGAAATCATGCCGAACATCCCGCAGATGTCCGCTTTCTGGTATGCCGTGCGTACTG
CGGTGATCAACGCCGCCAGCGGTCGTCAGACTGTCGATGAAGCCCTGAAAGACGCGCAGA
CTAATTCGAGCTCGAACAACAACAACAATAACAATAACAACAACCTCGGGATCGAGGGAAGG
ATTTCACATATGCAGACAATTAAGTGTGTTGTTGTGGGCGATGGCGCCGTTGGTAAAAACTG
TCTCCTGATATCCTACACAACAAACAAATTTCCATCGGAGTATGTACCGACTGTTTTTGACAA
CTATGCAGTCACAGTTATGATTGGTGGAGAACCATATACTCTTGGACTTTTTGATACTGCAGG
GCAAGAGGATTATGACAGATTACGACCGCTGAGTTATCCACAAACAGATGTATTTCTAGTCTG
TTTTTCAGTGGTCTCTCCATCTTCATTTGAAAACGTGAAAGAAAAGTGGGTGCCTGAGATAA
CTCACCACTGTCCAAAGACTCCTTTCTTGCTTGTTGGGACTCAAATTGATCTCAGAGATGAC
CCCTCTACTATTGAGAAACTTGCCAAGAACAAACAGAAGCCTATCACTCCAGAGACTGCTGA
AAAGCTGGCCCGTGACCTGAAGGCTGTCAAGTATGTGGAGTGTTCTGCACTTACACAGAGA
GGTCTGAAGAATGTGTTTGATGAGGCTATCCTAGCTGCCCTCGAGCCTCCGGAAACTCAAC
CCTAA

3）**pMAL-MBP-FRB-Cdc42/N12**

>Amino acid sequence

MKIEEGKLVIWINGDKGYNGLAEVGKKFEKDTGIKVTVEHPDKLEEKFPQVAATGDGPDIIFWAH
DRFGGYAQSGLLAEITPDKAFQDKLYPFTWDAVRYNGKLIAYPIAVEALSLIYNKDLLPNPPKTW
EEIPALDKELKAKGKSALMFNLQEPYFTWPLIAADGGYAFKYENGKYDIKDVGVDNAGAKAGLT
FLVDLIKNKHMNADTDYSIAEAAFNKGETAMTINGPWAWSNIDTSKVNYGVTVLPTFKGQPSKP
FVGVLSAGINAASPNKELAKEFLENYLLTDEGLEAVNKDKPLGAVALKSYEEELVKDPRIAATME
NAQKGEIMPNIPQMSAFWYAVRTAVINAASGRQTVDEALKDAQTNSSSNNNNNNNNNNLGIEG
RISHMEMWHEGLEEASRLYFGERNVKGMFEVLEPLHAMMERGPQTLKETSFNQAYGRDLMEA
QEWCRKYMKSGNVKDLTQAWDLYYHVFRRISKQQISYASRGGGSSGGGELQTIKCVVVGDGA
*

>DNA sequence

ATGAAAATCGAAGAAGGTAAACTGGTAATCTGGATTAACGGCGATAAAGGCTATAACGGTCT
CGCTGAAGTCGGTAAGAAATTCGAGAAAGATACCGGAATTAAAGTCACCGTTGAGCATCCG
GATAAACTGGAAGAGAAATTCCCACAGGTTGCGGCAACTGGCGATGGCCCTGACATTATCT
TCTGGGCACACGACCGCTTTGGTGGCTACGCTCAATCTGGCCTGTTGGCTGAAATCACCCC
GGACAAAGCGTTCCAGGACAAGCTGTATCCGTTTACCTGGGATGCCGTACGTTACAACGGC
AAGCTGATTGCTTACCCGATCGCTGTTGAAGCGTTATCGCTGATTTATAACAAAGATCTGCTG
CCGAACCCGCCAAAAACCTGGGAAGAGATCCCGGCGCTGGATAAAGAACTGAAAGCGAAA
GGTAAGAGCGCGCTGATGTTCAACCTGCAAGAACCGTACTTCACCTGGCCGCTGATTGCTG
CTGACGGGGGTTATGCGTTCAAGTATGAAAACGGCAAGTACGACATTAAAGACGTGGGCGT
GGATAACGCTGGCGCGAAAGCGGGTCTGACCTTCCTGGTTGACCTGATTAAAAACAAACAC
ATGAATGCAGACACCGATTACTCCATCGCAGAAGCTGCCTTTAATAAAGGCGAAACAGCGAT
GACCATCAACGGCCCGTGGGCATGGTCCAACATCGACACCAGCAAAGTGAATTATGGTGTA
ACGGTACTGCCGACCTTCAAGGGTCAACCATCCAAACCGTTCGTTGGCGTGCTGAGCGCA
GGTATTAACGCCGCCAGTCCGAACAAAGAGCTGGCAAAAGAGTTCCTCGAAAACTATCTGC
TGACTGATGAAGGTCTGGAAGCGGTTAATAAAGACAAACCGCTGGGTGCCGTAGCGCTGAA
GTCTTACGAGGAAGAGTTGGTGAAAGATCCGCGTATTGCCGCCACTATGGAAAACGCCCAG
AAAGGTGAAATCATGCCGAACATCCCGCAGATGTCCGCTTTCTGGTATGCCGTGCGTACTG
CGGTGATCAACGCCGCCAGCGGTCGTCAGACTGTCGATGAAGCCCTGAAAGACGCGCAGA
CTAATTCGAGCTCGAACAACAACAACAATAACAATAACAACAACCTCGGGATCGAGGGAAGG
ATTTCACATATGGAGATGTGGCATGAAGGCCTGGAAGAGGCATCTCGTTTGTACTTTGGGGA
AAGGAACGTGAAAGGCATGTTTGAGGTGCTGGAGCCCTTGCATGCTATGATGGAACGGGG
CCCCCAGACTCTGAAGGAAACATCCTTTAATCAGGCCTATGGTCGAGATTTAATGGAGGCCC
AAGAGTGGTGCAGGAAGTACATGAAATCAGGGAATGTCAAGGACCTCACCCAAGCCTGGG
ACCTCTATTATCATGTGTTCCGACGAATCTCAAAGCAGCAGATCTCGTACGCGTCCCGGGGC
GGTGGCTCATCTGGCGGAGGTGAGCTCCAGACAATTAAGTGTGTTGTTGTGGGCGATGGT
GCTTAA

4） **pET21b-Cdc42/13C-FKBP-MBP**

>Amino acid sequence

MGVGKTCLLISYTTNKFPSEYVPTVFDNYAVTVMIGGEPYTLGLFDTAGLEDYDRLRPLSYPQT
DVFLVCFSVVSPSSFENVKEKWVPEITHHCPKTPFLLVGTQIDLRDDPSTIEKLAKNKQKPITPET
AEKLARDLKAVKYVECSALTQRGLKNVFDEAILAALEPPETQPGDPNWELVYTARLQGGGSSG
GGQISYASRGGVQVETISPGDGRTFPKRGQTCVVHYTGMLEDGKKFDSSRDRNKPFKFMLGK
QEVIRGWEEGVAQMSVGQRAKLTISPDYAYGATGHPGIIPPHATLVFDVELLKLEKLENLYFQGE
EGKLVIWINGDKGYNGLAEVGKKFEKDTGIKVTVEHPDKLEEKFPQVAATGDGPDIIFWAHDRF
GGYAQSGLLAEITPDKAFQDKLYPFTWDAVRYNGKLIAYPIAVEALSLIYNKDLLPNPPKTWEEIP
ALDKELKAKGKSALMFNLQEPYFTWPLIAADGGYAFKYENGKYDIKDVGVDNAGAKAGLTFLVD
LIKNKHMNADTDYSIAEAAFNKGETAMTINGPWAWSNIDTSKVNYGVTVLPTFKGQPSKPFVGV
LSAGINAASPNKELAKEFLENYLLTDEGLEAVNKDKPLGAVALKSYEEELAKDPRIAATMENAQK
GEIMPNIPQMSAFWYAVRTAVINAASGRQTVDEALKDAQTNSSS*

>DNA sequence

ATGGGGGTTGGTAAAACATGTCTCCTGATATCCTACACAACAAACAAATTTCCATCGGAGTAT
GTACCGACTGTTTTTGACAACTATGCAGTCACAGTTATGATTGGTGGAGAACCATATACTCTT
GGACTTTTTGATACTGCAGGGCTAGAGGATTATGACAGATTACGACCGCTGAGTTATCCACA
AACAGATGTATTTCTAGTCTGTTTTTCAGTGGTCTCTCCATCTTCATTTGAAAACGTGAAAGA
AAAGTGGGTGCCTGAGATAACTCACCACTGTCCAAAGACTCCTTTCTTGCTTGTTGGGACT
CAAATTGATCTCAGAGATGACCCCTCTACTATTGAGAAACTTGCCAAGAACAAACAGAAGCC
TATCACTCCAGAGACTGCTGAAAAGCTGGCCCGTGACCTGAAGGCTGTCAAGTATGTGGAG
TGTTCTGCACTTACACAGAGAGGTCTGAAGAATGTGTTTGATGAGGCTATCCTAGCTGCCCT
CGAGCCTCCGGAAACTCAACCCGGGGATCCCAATTGGGAGCTCGTGTACACGGCGCGCCT
GCAGGGAGGTGGCTCATCTGGCGGAGGTCAGATCTCGTACGCGTCCCGGGGCGGAGTGC
AGGTGGAAACCATCTCCCCAGGAGACGGGCGCACCTTCCCCAAGCGCGGCCAGACCTGC
GTGGTGCACTACACCGGGATGCTTGAAGATGGAAAGAAATTTGATTCCTCCCGGGACAGAA
ACAAGCCCTTTAAGTTTATGCTAGGCAAGCAGGAGGTGATCCGAGGCTGGGAAGAAGGGG
TTGCCCAGATGAGTGTGGGTCAGAGAGCCAAACTGACTATATCTCCAGATTATGCCTATGGT
GCCACTGGGCACCCAGGCATCATCCCACCACATGCCACTCTCGTCTTCGATGTGGAGCTTC
TAAAACTGGAAAAGCTTGAGAACCTGTACTTCCAGGGCGAAGAAGGTAAACTGGTAATCTG
GATTAACGGCGATAAAGGCTATAACGGTCTCGCTGAAGTCGGTAAGAAATTCGAGAAAGATA
CCGGAATTAAAGTCACCGTTGAGCATCCGGATAAACTGGAAGAGAAATTCCCACAGGTTGC
GGCAACTGGCGATGGCCCTGACATTATCTTCTGGGCACACGACCGCTTTGGTGGCTACGCT
CAATCTGGCCTGTTGGCTGAAATCACCCCGGACAAAGCGTTCCAGGACAAGCTGTATCCGT
TTACCTGGGATGCCGTACGTTACAACGGCAAGCTGATTGCTTACCCGATCGCTGTTGAAGC
GTTATCGCTGATTTATAACAAAGACCTGCTGCCGAACCCGCCAAAAACCTGGGAAGAGATCC
CGGCGCTGGATAAAGAACTGAAAGCGAAAGGTAAGAGCGCGCTGATGTTCAACCTGCAAG
AACCGTACTTCACCTGGCCGCTGATTGCTGCTGACGGGGGTTATGCGTTCAAGTATGAAAA
CGGCAAGTACGACATTAAAGACGTGGGCGTGGATAACGCTGGCGCGAAAGCGGGTCTGAC
CTTCCTGGTTGACCTGATTAAAAACAAACACATGAATGCAGACACCGATTACTCCATCGCAG
AAGCTGCCTTTAATAAAGGCGAAACAGCGATGACCATCAACGGCCCGTGGGCATGGTCCAA
CATCGACACCAGCAAAGTGAATTATGGTGTAACGGTACTGCCGACCTTCAAGGGTCAACCAT
CCAAACCGTTCGTTGGCGTGCTGAGCGCAGGTATTAACGCCGCCAGTCCGAACAAAGAGC
TGGCAAAAGAGTTCCTCGAAAACTATCTGCTGACTGATGAAGGTCTGGAAGCGGTTAATAAA

GACAAACCGCTGGGTGCCGTAGCGCTGAAGTCTTACGAGGAAGAGTTGGCGAAAGATCCA
CGTATTGCCGCCACTATGGAAAACGCCCAGAAAGGTGAAATCATGCCGAACATCCCGCAGA
TGTCCGCTTTCTGGTATGCCGTGCGTACTGCGGTGATCAACGCCGCCAGCGGTCGTCAGA
CTGTCGATGAAGCCCTGAAAGACGCGCAGACTAATTCCAGCTCGTAG

5） **pMAL-MBP-Rac1-Q61L**

>Amino acid sequence

MKIEEGKLVIWINGDKGYNGLAEVGKKFEKDTGIKVTVEHPDKLEEKFPQVAATGDGPDIIFWAH
DRFGGYAQSGLLAEITPDKAFQDKLYPFTWDAVRYNGKLIAYPIAVEALSLIYNKDLLPNPPKTW
EEIPALDKELKAKGKSALMFNLQEPYFTWPLIAADGGYAFKYENGKYDIKDVGVDNAGAKAGLT
FLVDLIKNKHMNADTDYSIAEAAFNKGETAMTINGPWAWSNIDTSKVNYGVTVLPTFKGQPSKP
FVGVLSAGINAASPNKELAKEFLENYLLTDEGLEAVNKDKPLGAVALKSYEEELVKDPRIAATME
NAQKGEIMPNIPQMSAFWYAVRTAVINAASGRQTVDEALKDAQTNSSSNNNNNNNNNNLGIEG
RISHMQAIKCVVVGDGAVGKTCLLISYTTNAFPGEYIPTVFDNYSANVMVDGKPVNLGLWDTAG
LDYDRLRPLSYPQTDVFLICFSLVSPASFENVRAKWYPEVRHHCPNTPIILVGTKLDLRDDKDTIE
KLKEKKLTPITYPQGLAMAKEIGAVKYLECSALTQRGLKTVFDEAIRAVLCPPPV*

>DNA sequence

ATGAAAATCGAAGAAGGTAAACTGGTAATCTGGATTAACGGCGATAAAGGCTATAACGGTCT
CGCTGAAGTCGGTAAGAAATTCGAGAAAGATACCGGAATTAAAGTCACCGTTGAGCATCCG
GATAAACTGGAAGAGAAATTCCCACAGGTTGCGGCAACTGGCGATGGCCCTGACATTATCT
TCTGGGCACACGACCGCTTTGGTGGCTACGCTCAATCTGGCCTGTTGGCTGAAATCACCCC
GGACAAAGCGTTCCAGGACAAGCTGTATCCGTTTACCTGGGATGCCGTACGTTACAACGGC
AAGCTGATTGCTTACCCGATCGCTGTTGAAGCGTTATCGCTGATTTATAACAAAGATCTGCTG
CCGAACCCGCCAAAAACCTGGGAAGAGATCCCGGCGCTGGATAAAGAACTGAAAGCGAAA
GGTAAGAGCGCGCTGATGTTCAACCTGCAAGAACCGTACTTCACCTGGCCGCTGATTGCTG
CTGACGGGGGTTATGCGTTCAAGTATGAAAACGGCAAGTACGACATTAAAGACGTGGGCGT
GGATAACGCTGGCGCGAAAGCGGGTCTGACCTTCCTGGTTGACCTGATTAAAAACAAACAC
ATGAATGCAGACACCGATTACTCCATCGCAGAAGCTGCCTTTAATAAAGGCGAAACAGCGAT
GACCATCAACGGCCCGTGGGCATGGTCCAACATCGACACCAGCAAAGTGAATTATGGTGTA
ACGGTACTGCCGACCTTCAAGGGTCAACCATCCAAACCGTTCGTTGGCGTGCTGAGCGCA
GGTATTAACGCCGCCAGTCCGAACAAAGAGCTGGCAAAAGAGTTCCTCGAAAACTATCTGC
TGACTGATGAAGGTCTGGAAGCGGTTAATAAAGACAAACCGCTGGGTGCCGTAGCGCTGAA
GTCTTACGAGGAAGAGTTGGTGAAAGATCCGCGTATTGCCGCCACTATGGAAAACGCCCAG
AAAGGTGAAATCATGCCGAACATCCCGCAGATGTCCGCTTTCTGGTATGCCGTGCGTACTG
CGGTGATCAACGCCGCCAGCGGTCGTCAGACTGTCGATGAAGCCCTGAAAGACGCGCAGA
CTAATTCGAGCTCGAACAACAACAACAATAACAATAACAACAACCTCGGGATCGAGGGAAGG
ATTTCACATATGCAGGCCATCAAGTGTGTGGTGGTGGGAGACGGAGCTGTAGGTAAAACTT
GCCTACTGATCAGTTACACAACCAATGCATTTCCTGGAGAATATATCCCTACTGTCTTTGACA
ATTATTCTGCCAATGTTATGGTAGATGGAAAACCGGTGAATCTGGGCTTATGGGATACAGCTG
GACTAGATTATGACAGATTACGCCCCCTATCCTATCCGCAAACAGATGTGTTCTTAATTTGCTT
TTCCCTTGTGAGTCCTGCATCATTTGAAAATGTCCGTGCAAAGTGGTATCCTGAGGTGCGG
CACCACTGTCCCAACACTCCCATCATCCTAGTGGGAACTAAACTTGATCTTAGGGATGATAA
AGACACGATCGAGAAACTGAAGGAGAAGAAGCTGACTCCCATCACCTATCCGCAGGGTCTA
GCCATGGCTAAGGAGATTGGTGCTGTAAAATACCTGGAGTGCTCGGCGCTCACACAGCGA
GGCCTCAAGACAGTGTTTGACGAAGCGATCCGAGCAGTCCTCTGCCCGCCTCCCGTGTAA

6）**pMAL-MBP-Rac1-T17N**

>Amino acid sequence

MKIEEGKLVIWINGDKGYNGLAEVGKKFEKDTGIKVTVEHPDKLEEKFPQVAATGDGPDIIFWAH
DRFGGYAQSGLLAEITPDKAFQDKLYPFTWDAVRYNGKLIAYPIAVEALSLIYNKDLLPNPPKTW
EEIPALDKELKAKGKSALMFNLQEPYFTWPLIAADGGYAFKYENGKYDIKDVGVDNAGAKAGLT
FLVDLIKNKHMNADTDYSIAEAAFNKGETAMTINGPWAWSNIDTSKVNYGVTVLPTFKGQPSKP
FVGVLSAGINAASPNKELAKEFLENYLLTDEGLEAVNKDKPLGAVALKSYEEELVKDPRIAATME
NAQKGEIMPNIPQMSAFWYAVRTAVINAASGRQTVDEALKDAQTNSSSNNNNNNNNNNLGIEG
RISHMQAIKCVVVGDGAVGKNCLLISYTTNAFPGEYIPTVFDNYSANVMVDGKPVNLGLWDTAG
QEDYDRLRPLSYPQTDVFLICFSLVSPASFENVRAKWYPEVRHHCPNTPIILVGTKLDLRDDKDT
IEKLKEKKLTPITYPQGLAMAKEIGAVKYLECSALTQRGLKTVFDEAIRAVLCPPPV*

>DNA sequence

ATGAAAATCGAAGAAGGTAAACTGGTAATCTGGATTAACGGCGATAAAGGCTATAACGGTCT
CGCTGAAGTCGGTAAGAAATTCGAGAAAGATACCGGAATTAAAGTCACCGTTGAGCATCCG
GATAAACTGGAAGAGAAATTCCCACAGGTTGCGGCAACTGGCGATGGCCCTGACATTATCT
TCTGGGCACACGACCGCTTTGGTGGCTACGCTCAATCTGGCCTGTTGGCTGAAATCACCCC
GGACAAAGCGTTCCAGGACAAGCTGTATCCGTTTACCTGGGATGCCGTACGTTACAACGGC
AAGCTGATTGCTTACCCGATCGCTGTTGAAGCGTTATCGCTGATTTATAACAAAGATCTGCTG
CCGAACCCGCCAAAAACCTGGGAAGAGATCCCGGCGCTGGATAAAGAACTGAAAGCGAAA
GGTAAGAGCGCGCTGATGTTCAACCTGCAAGAACCGTACTTCACCTGGCCGCTGATTGCTG
CTGACGGGGGTTATGCGTTCAAGTATGAAAACGGCAAGTACGACATTAAAGACGTGGGCGT
GGATAACGCTGGCGCGAAAGCGGGTCTGACCTTCCTGGTTGACCTGATTAAAAACAAACAC
ATGAATGCAGACACCGATTACTCCATCGCAGAAGCTGCCTTTAATAAAGGCGAAACAGCGAT
GACCATCAACGGCCCGTGGGCATGGTCCAACATCGACACCAGCAAAGTGAATTATGGTGTA
ACGGTACTGCCGACCTTCAAGGGTCAACCATCCAAACCGTTCGTTGGCGTGCTGAGCGCA
GGTATTAACGCCGCCAGTCCGAACAAAGAGCTGGCAAAAGAGTTCCTCGAAAACTATCTGC
TGACTGATGAAGGTCTGGAAGCGGTTAATAAAGACAAACCGCTGGGTGCCGTAGCGCTGAA
GTCTTACGAGGAAGAGTTGGTGAAAGATCCGCGTATTGCCGCCACTATGGAAAACGCCCAG
AAAGGTGAAATCATGCCGAACATCCCGCAGATGTCCGCTTTCTGGTATGCCGTGCGTACTG
CGGTGATCAACGCCGCCAGCGGTCGTCAGACTGTCGATGAAGCCCTGAAAGACGCGCAGA
CTAATTCGAGCTCGAACAACAACAACAATAACAATAACAACAACCTCGGGATCGAGGGAAGG
ATTTCACATATGCAGGCCATCAAGTGTGTGGTGGTGGGAGACGGAGCTGTAGGTAAAAACT
GCCTACTGATCAGTTACACAACCAATGCATTTCCTGGAGAATATATCCCTACTGTCTTTGACA
ATTATTCTGCCAATGTTATGGTAGATGGAAAACCGGTGAATCTGGGCTTATGGGATACAGCTG
GACAAGAAGATTATGACAGATTACGCCCCCTATCCTATCCGCAAACAGATGTGTTCTTAATTT
GCTTTTCCCTTGTGAGTCCTGCATCATTTGAAAATGTCCGTGCAAAGTGGTATCCTGAGGTG
CGGCACCACTGTCCCAACACTCCCATCATCCTAGTGGGAACTAAACTTGATCTTAGGGATGA
TAAAGACACGATTGAGAAACTGAAGGAGAAGAAGCTGACTCCCATCACCTATCCGCAGGGT
CTAGCCATGGCTAAGGAGATTGGTGCTGTAAAATACCTGGAGTGCTCGGCGCTCACACAGC
GAGGCCTCAAGACAGTGTTTGACGAAGCGATCCGAGCAGTCCTCTGCCCGCCTCCCGTGT
AA

7）**pMAL-MBP-FRB-Rac1/N12**

>Amino acid sequence

MKIEEGKLVIWINGDKGYNGLAEVGKKFEKDTGIKVTVEHPDKLEEKFPQVAATGDGPDIIFWAH
DRFGGYAQSGLLAEITPDKAFQDKLYPFTWDAVRYNGKLIAYPIAVEALSLIYNKDLLPNPPKTW
EEIPALDKELKAKGKSALMFNLQEPYFTWPLIAADGGYAFKYENGKYDIKDVGVDNAGAKAGLT
FLVDLIKNKHMNADTDYSIAEAAFNKGETAMTINGPWAWSNIDTSKVNYGVTVLPTFKGQPSKP
FVGVLSAGINAASPNKELAKEFLENYLLTDEGLEAVNKDKPLGAVALKSYEEELVKDPRIAATME
NAQKGEIMPNIPQMSAFWYAVRTAVINAASGRQTVDEALKDAQTNSSSNNNNNNNNNLGIEG
RISHMSMGGREMWHEGLEEASRLYFGERNVKGMFEVLEPLHAMMERGPQTLKETSFNQAYG
RDLMEAQEWCRKYMKSGNVKDLTQAWDLYYHVFRRISKQQISYASRGGGSSGGGELQAIKCV
VVGDGA*

>DNA sequence

ATGAAAATCGAAGAAGGTAAACTGGTAATCTGGATTAACGGCGATAAAGGCTATAACGGTCT
CGCTGAAGTCGGTAAGAAATTCGAGAAAGATACCGGAATTAAAGTCACCGTTGAGCATCCG
GATAAACTGGAAGAGAAATTCCCACAGGTTGCGGCAACTGGCGATGGCCCTGACATTATCT
TCTGGGCACACGACCGCTTTGGTGGCTACGCTCAATCTGGCCTGTTGGCTGAAATCACCCC
GGACAAAGCGTTCCAGGACAAGCTGTATCCGTTTACCTGGGATGCCGTACGTTACAACGGC
AAGCTGATTGCTTACCCGATCGCTGTTGAAGCGTTATCGCTGATTTATAACAAAGATCTGCTG
CCGAACCCGCCAAAAACCTGGGAAGAGATCCCGGCGCTGGATAAAGAACTGAAAGCGAAA
GGTAAGAGCGCGCTGATGTTCAACCTGCAAGAACCGTACTTCACCTGGCCGCTGATTGCTG
CTGACGGGGGTTATGCGTTCAAGTATGAAAACGGCAAGTACGACATTAAAGACGTGGGCGT
GGATAACGCTGGCGCGAAAGCGGGTCTGACCTTCCTGGTTGACCTGATTAAAAACAAACAC
ATGAATGCAGACACCGATTACTCCATCGCAGAAGCTGCCTTTAATAAAGGCGAAACAGCGAT
GACCATCAACGGCCCGTGGGCATGGTCCAACATCGACACCAGCAAAGTGAATTATGGTGTA
ACGGTACTGCCGACCTTCAAGGGTCAACCATCCAAACCGTTCGTTGGCGTGCTGAGCGCA
GGTATTAACGCCGCCAGTCCGAACAAAGAGCTGGCAAAAGAGTTCCTCGAAAACTATCTGC
TGACTGATGAAGGTCTGGAAGCGGTTAATAAAGACAAACCGCTGGGTGCCGTAGCGCTGAA
GTCTTACGAGGAAGAGTTGGTGAAAGATCCGCGTATTGCCGCCACTATGGAAAACGCCCAG
AAAGGTGAAATCATGCCGAACATCCCGCAGATGTCCGCTTTCTGGTATGCCGTGCGTACTG
CGGTGATCAACGCCGCCAGCGGTCGTCAGACTGTCGATGAAGCCCTGAAAGACGCGCAGA
CTAATTCGAGCTCGAACAACAACAACAATAACAATAACAACAACCTCGGGATCGAGGGAAGG
ATTTCACATATGTCCATGGGCGGCCGCGAGATGTGGCATGAAGGCCTGGAAGAGGCATCTC
GTTTGTACTTTGGGGAAAGGAACGTGAAAGGCATGTTTGAGGTGCTGGAGCCCTTGCATGC
TATGATGGAACGGGGCCCCCAGACTCTGAAGGAAACATCCTTTAATCAGGCCTATGGTCGA
GATTTAATGGAGGCCCAAGAGTGGTGCAGGAAGTACATGAAATCAGGGAATGTCAAGGACC
TCACCCAAGCCTGGGACCTCTATTATCATGTGTTCCGACGAATCTCAAAGCAGCAGATCTCG
TACGCGTCCCGGGGCGGTGGCTCATCTGGCGGAGGTGAGCTCCAGGCCATCAAGTGTGT
GGTGGTGGGAGACGGAGCTTAA

8）**pET21b-Rac1/13C-FKBP-MBP**

>Amino acid sequence

MVGKTCLLISYTTNAFPGEYIPTVFDNYSANVMVDGKPVNLGLWDTAGLEDYDRLRPLSYPQTD
VFLICFSLVSPASFENVRAKWYPEVRHHCPNTPIILVGTKLDLRDDKDTIEKLKEKKLTPITYPQGL
AMAKEIGAVKYLECSALTQRGLKTVFDEAIRAVLCPPPVGDPNWELVYTARLQGGGSSGGGQIS
YASRGGVQVETISPGDGRTFPKRGQTCVVHYTGMLEDGKKFDSSRDRNKPFKFMLGKQEVIR
GWEEGVAQMSVGQRAKLTISPDYAYGATGHPGIIPPHATLVFDVELLKLEKLENLYFQGEEGKLV
IWINGDKGYNGLAEVGKKFEKDTGIKVTVEHPDKLEEKFPQVAATGDGPDIIFWAHDRFGGYAQ
SGLLAEITPDKAFQDKLYPFTWDAVRYNGKLIAYPIAVEALSLIYNKDLLPNPPKTWEEIPALDKEL
KAKGKSALMFNLQEPYFTWPLIAADGGYAFKYENGKYDIKDVGVDNAGAKAGLTFLVDLIKNKH
MNADTDYSIAEAAFNKGETAMTINGPWAWSNIDTSKVNYGVTVLPTFKGQPSKPFVGVLSAGIN
AASPNKELAKEFLENYLLTDEGLEAVNKDKPLGAVALKSYEEELAKDPRIAATMENAQKGEIMPN
IPQMSAFWYAVRTAVINAASGRQTVDEALKDAQTNSSS*

>DNA sequence

ATGGTAGGTAAAACTTGCCTACTGATCAGTTACACAACCAATGCATTTCCTGGAGAATATATC
CCTACTGTCTTTGACAATTATTCTGCCAATGTTATGGTAGATGGAAAACCGGTGAATCTGGGC
TTATGGGATACAGCTGGACTAGAAGATTATGACAGATTACGCCCCCTATCCTATCCGCAAACA
GATGTGTTCTTAATTTGCTTTTCCCTTGTGAGTCCTGCATCATTTGAAAATGTCCGTGCAAAG
TGGTATCCTGAGGTGCGGCACCACTGTCCCAACACTCCCATCATCCTAGTGGGAACTAAAC
TTGATCTTAGGGATGATAAAGACACGATCGAGAAACTGAAGGAGAAGAAGCTGACTCCCATC
ACCTATCCGCAGGGTCTAGCCATGGCTAAGGAGATTGGTGCTGTAAAATACCTGGAGTGCT
CGGCGCTCACACAGCGAGGCCTCAAGACAGTGTTTGACGAAGCGATCCGAGCAGTCCTCT
GCCCGCCTCCCGTGGGGGATCCCAATTGGGAGCTCGTGTACACGGCGCGCCTGCAGGGA
GGTGGCTCATCTGGCGGAGGTCAGATCTCGTACGCGTCCCGGGGCGGAGTGCAGGTGGA
AACCATCTCCCCAGGAGACGGGCGCACCTTCCCCAAGCGCGGCCAGACCTGCGTGGTGC
ACTACACCGGGATGCTTGAAGATGGAAAGAAATTTGATTCCTCCCGGGACAGAAACAAGCC
CTTTAAGTTTATGCTAGGCAAGCAGGAGGTGATCCGAGGCTGGGAAGAAGGGGTTGCCCA
GATGAGTGTGGGTCAGAGAGCCAAACTGACTATATCTCCAGATTATGCCTATGGTGCCACTG
GGCACCCAGGCATCATCCCACCACATGCCACTCTCGTCTTCGATGTGGAGCTTCTAAAACT
GGAAAAGCTTGAGAACCTGTACTTCCAGGGCGAAGAAGGTAAACTGGTAATCTGGATTAAC
GGCGATAAAGGCTATAACGGTCTCGCTGAAGTCGGTAAGAAATTCGAGAAAGATACCGGAAT
TAAAGTCACCGTTGAGCATCCGGATAAACTGGAAGAGAAATTCCCACAGGTTGCGGCAACT
GGCGATGGCCCTGACATTATCTTCTGGGCACACGACCGCTTTGGTGGCTACGCTCAATCTG
GCCTGTTGGCTGAAATCACCCCGGACAAAGCGTTCCAGGACAAGCTGTATCCGTTTACCTG
GGATGCCGTACGTTACAACGGCAAGCTGATTGCTTACCCGATCGCTGTTGAAGCGTTATCG
CTGATTTATAACAAAGACCTGCTGCCGAACCCGCCAAAAACCTGGGAAGAGATCCCGGCGC
TGGATAAAGAACTGAAAGCGAAAGGTAAGAGCGCGCTGATGTTCAACCTGCAAGAACCGTA
CTTCACCTGGCCGCTGATTGCTGCTGACGGGGGTTATGCGTTCAAGTATGAAAACGGCAAG
TACGACATTAAAGACGTGGGCGTGGATAACGCTGGCGCGAAAGCGGGTCTGACCTTCCTG
GTTGACCTGATTAAAAACAAACACATGAATGCAGACACCGATTACTCCATCGCAGAAGCTGC
CTTTAATAAAGGCGAAACAGCGATGACCATCAACGGCCCGTGGGCATGGTCCAACATCGAC
ACCAGCAAAGTGAATTATGGTGTAACGGTACTGCCGACCTTCAAGGGTCAACCATCCAAAC
CGTTCGTTGGCGTGCTGAGCGCAGGTATTAACGCCGCCAGTCCGAACAAAGAGCTGGCAA
AAGAGTTCCTCGAAAACTATCTGCTGACTGATGAAGGTCTGGAAGCGGTTAATAAAGACAAA

CCGCTGGGTGCCGTAGCGCTGAAGTCTTACGAGGAAGAGTTGGCGAAAGATCCACGTATT
GCCGCCACTATGGAAAACGCCCAGAAAGGTGAAATCATGCCGAACATCCCGCAGATGTCCG
CTTTCTGGTATGCCGTGCGTACTGCGGTGATCAACGCCGCCAGCGGTCGTCAGACTGTCG
ATGAAGCCCTGAAAGACGCGCAGACTAATTCCAGCTCGTAG

#### 9）pMAL-MBP-RhoA-Q63L

>Amino acid sequence

MKIEEGKLVIWINGDKGYNGLAEVGKKFEKDTGIKVTVEHPDKLEEKFPQVAATGDGPDIIFWAH
DRFGGYAQSGLLAEITPDKAFQDKLYPFTWDAVRYNGKLIAYPIAVEALSLIYNKDLLPNPPKTW
EEIPALDKELKAKGKSALMFNLQEPYFTWPLIAADGGYAFKYENGKYDIKDVGVDNAGAKAGLT
FLVDLIKNKHMNADTDYSIAEAAFNKGETAMTINGPWAWSNIDTSKVNYGVTVLPTFKGQPSKP
FVGVLSAGINAASPNKELAKEFLENYLLTDEGLEAVNKDKPLGAVALKSYEEELVKDPRIAATME
NAQKGEIMPNIPQMSAFWYAVRTAVINAASGRQTVDEALKDAQTNSSSNNNNNNNNNNLGIEG
RISHMAAIRKKLVIVGDGACGKTCLLIVFSKDQFPEVYVPTVFENYVADIEVDGKQVELALWDTA
GLEDYDRLRPLSYPDTDVILMCFSIDSPDSLENIPEKWTPEVKHFCPNVPIILVGNKKDLRNDEH
TRRELAKMKQEPVKPEEGRDMANRIGAFGYMECSAKTKDGVREVFEMATRAALQA*

>DNA sequence

ATGAAAATCGAAGAAGGTAAACTGGTAATCTGGATTAACGGCGATAAAGGCTATAACGGTCT
CGCTGAAGTCGGTAAGAAATTCGAGAAAGATACCGGAATTAAAGTCACCGTTGAGCATCCG
GATAAACTGGAAGAGAAATTCCCACAGGTTGCGGCAACTGGCGATGGCCCTGACATTATCT
TCTGGGCACACGACCGCTTTGGTGGCTACGCTCAATCTGGCCTGTTGGCTGAAATCACCCC
GGACAAAGCGTTCCAGGACAAGCTGTATCCGTTTACCTGGGATGCCGTACGTTACAACGGC
AAGCTGATTGCTTACCCGATCGCTGTTGAAGCGTTATCGCTGATTTATAACAAAGATCTGCTG
CCGAACCCGCCAAAAACCTGGGAAGAGATCCCGGCGCTGGATAAAGAACTGAAAGCGAAA
GGTAAGAGCGCGCTGATGTTCAACCTGCAAGAACCGTACTTCACCTGGCCGCTGATTGCTG
CTGACGGGGGTTATGCGTTCAAGTATGAAAACGGCAAGTACGACATTAAAGACGTGGGCGT
GGATAACGCTGGCGCGAAAGCGGGTCTGACCTTCCTGGTTGACCTGATTAAAAACAAACAC
ATGAATGCAGACACCGATTACTCCATCGCAGAAGCTGCCTTTAATAAAGGCGAAACAGCGAT
GACCATCAACGGCCCGTGGGCATGGTCCAACATCGACACCAGCAAAGTGAATTATGGTGTA
ACGGTACTGCCGACCTTCAAGGGTCAACCATCCAAACCGTTCGTTGGCGTGCTGAGCGCA
GGTATTAACGCCGCCAGTCCGAACAAAGAGCTGGCAAAAGAGTTCCTCGAAAACTATCTGC
TGACTGATGAAGGTCTGGAAGCGGTTAATAAAGACAAACCGCTGGGTGCCGTAGCGCTGAA
GTCTTACGAGGAAGAGTTGGTGAAAGATCCGCGTATTGCCGCCACTATGGAAAACGCCCAG
AAAGGTGAAATCATGCCGAACATCCCGCAGATGTCCGCTTTCTGGTATGCCGTGCGTACTG
CGGTGATCAACGCCGCCAGCGGTCGTCAGACTGTCGATGAAGCCCTGAAAGACGCGCAGA
CTAATTCGAGCTCGAACAACAACAACAATAACAATAACAACAACCTCGGGATCGAGGGAAGG
ATTTCACATATGGCTGCCATCCGGAAGAAACTGGTGATTGTTGGTGATGGAGCCTGTGGAAA
GACATGCTTGCTCATAGTCTTCAGCAAGGACCAGTTCCCAGAGGTGTATGTGCCCACAGTG
TTTGAGAACTATGTGGCAGATATCGAGGTGGATGGAAAGCAGGTAGAGTTGGCTTTGTGGG
ACACAGCTGGGCTGGAAGATTATGATCGCCTGAGGCCCCTCTCCTACCCAGATACCGATGT
TATACTGATGTGTTTTTCCATCGACAGCCCTGATAGTTTAGAAAACATCCCAGAAAAGTGGAC
CCCAGAAGTCAAGCATTTCTGTCCCAACGTGCCCATCATCCTGGTTGGGAATAAGAAGGAT
CTTCGGAATGATGAGCACACAAGGCGGGAGCTAGCCAAGATGAAGCAGGAGCCGGTGAAA
CCTGAAGAAGGCAGAGATATGGCAAACAGGATTGGCGCTTTTGGGTACATGGAGTGTTCAG
CAAAGACCAAAGATGGAGTGAGAGAGGTTTTTGAAATGGCTACGAGAGCTGCTCTGCAAGC
TTAA

10） **pMAL-MBP-RhoA-T19N**

>Amino acid sequence

MKIEEGKLVIWINGDKGYNGLAEVGKKFEKDTGIKVTVEHPDKLEEKFPQVAATGDGPDIIFWAH
DRFGGYAQSGLLAEITPDKAFQDKLYPFTWDAVRYNGKLIAYPIAVEALSLIYNKDLLPNPPKTW
EEIPALDKELKAKGKSALMFNLQEPYFTWPLIAADGGYAFKYENGKYDIKDVGVDNAGAKAGLT
FLVDLIKNKHMNADTDYSIAEAAFNKGETAMTINGPWAWSNIDTSKVNYGVTVLPTFKGQPSKP
FVGVLSAGINAASPNKELAKEFLENYLLTDEGLEAVNKDKPLGAVALKSYEEELVKDPRIAATME
NAQKGEIMPNIPQMSAFWYAVRTAVINAASGRQTVDEALKDAQTNSSSNNNNNNNNNNLGIEG
RISHMAAIRKKLVIVGDGACGKNCLLIVFSKDQFPEVYVPTVFENYVADIEVDGKQVELALWDTA
GQEDYDRLRPLSYPDTDVILMCFSIDSPDSLENIPEKWTPEVKHFCPNVPIILVGNKKDLRNDEH
TRRELAKMKQEPVKPEEGRDMANRIGAFGYMECSAKTKDGVREVFEMATRAALQA*

>DNA sequence

ATGAAAATCGAAGAAGGTAAACTGGTAATCTGGATTAACGGCGATAAAGGCTATAACGGTCT
CGCTGAAGTCGGTAAGAAATTCGAGAAAGATACCGGAATTAAAGTCACCGTTGAGCATCCG
GATAAACTGGAAGAGAAATTCCCACAGGTTGCGGCAACTGGCGATGGCCCTGACATTATCT
TCTGGGCACACGACCGCTTTGGTGGCTACGCTCAATCTGGCCTGTTGGCTGAAATCACCCC
GGACAAAGCGTTCCAGGACAAGCTGTATCCGTTTACCTGGGATGCCGTACGTTACAACGGC
AAGCTGATTGCTTACCCGATCGCTGTTGAAGCGTTATCGCTGATTTATAACAAAGATCTGCTG
CCGAACCCGCCAAAAACCTGGGAAGAGATCCCGGCGCTGGATAAAGAACTGAAAGCGAAA
GGTAAGAGCGCGCTGATGTTCAACCTGCAAGAACCGTACTTCACCTGGCCGCTGATTGCTG
CTGACGGGGGTTATGCGTTCAAGTATGAAAACGGCAAGTACGACATTAAAGACGTGGGCGT
GGATAACGCTGGCGCGAAAGCGGGTCTGACCTTCCTGGTTGACCTGATTAAAAACAAACAC
ATGAATGCAGACACCGATTACTCCATCGCAGAAGCTGCCTTTAATAAAGGCGAAACAGCGAT
GACCATCAACGGCCCGTGGGCATGGTCCAACATCGACACCAGCAAAGTGAATTATGGTGTA
ACGGTACTGCCGACCTTCAAGGGTCAACCATCCAAACCGTTCGTTGGCGTGCTGAGCGCA
GGTATTAACGCCGCCAGTCCGAACAAAGAGCTGGCAAAAGAGTTCCTCGAAAACTATCTGC
TGACTGATGAAGGTCTGGAAGCGGTTAATAAAGACAAACCGCTGGGTGCCGTAGCGCTGAA
GTCTTACGAGGAAGAGTTGGTGAAAGATCCGCGTATTGCCGCCACTATGGAAAACGCCCAG
AAAGGTGAAATCATGCCGAACATCCCGCAGATGTCCGCTTTCTGGTATGCCGTGCGTACTG
CGGTGATCAACGCCGCCAGCGGTCGTCAGACTGTCGATGAAGCCCTGAAAGACGCGCAGA
CTAATTCGAGCTCGAACAACAACAACAATAACAATAACAACAACCTCGGGATCGAGGGAAGG
ATTTCACATATGGCTGCCATCCGGAAGAAACTGGTGATTGTTGGTGATGGAGCCTGTGGAAA
GAACTGCTTGCTCATAGTCTTCAGCAAGGACCAGTTCCCAGAGGTGTATGTGCCCACAGTG
TTTGAGAACTATGTGGCAGATATCGAGGTGGATGGAAAGCAGGTAGAGTTGGCTTTGTGGG
ACACAGCTGGGCAGGAAGATTATGATCGCCTGAGGCCCCTCTCCTACCCAGATACCGATGT
TATACTGATGTGTTTTTCCATCGACAGCCCTGATAGTTTAGAAAACATCCCAGAAAAGTGGAC
CCCAGAAGTCAAGCATTTCTGTCCCAACGTGCCCATCATCCTGGTTGGGAATAAGAAGGAT
CTTCGGAATGATGAGCACACAAGGCGGGAGCTAGCCAAGATGAAGCAGGAGCCGGTGAAA
CCTGAAGAAGGCAGAGATATGGCAAACAGGATTGGCGCTTTTGGGTACATGGAGTGTTCAG
CAAAGACCAAAGATGGAGTGAGAGAGGTTTTTGAAATGGCTACGAGAGCTGCTCTGCAAGC
TTAA

11） **pMAL-MBP-FRB-RhoA/N12**

>Amino acid sequence

MKIEEGKLVIWINGDKGYNGLAEVGKKFEKDTGIKVTVEHPDKLEEKFPQVAATGDGPDIIFWAH
DRFGGYAQSGLLAEITPDKAFQDKLYPFTWDAVRYNGKLIAYPIAVEALSLIYNKDLLPNPPKTW
EEIPALDKELKAKGKSALMFNLQEPYFTWPLIAADGGYAFKYENGKYDIKDVGVDNAGAKAGLT
FLVDLIKNKHMNADTDYSIAEAAFNKGETAMTINGPWAWSNIDTSKVNYGVTVLPTFKGQPSKP
FVGVLSAGINAASPNKELAKEFLENYLLTDEGLEAVNKDKPLGAVALKSYEEELVKDPRIAATME
NAQKGEIMPNIPQMSAFWYAVRTAVINAASGRQTVDEALKDAQTNSSSNNNNNNNNNNLGIEG
RISHMSMGGREMWHEGLEEASRLYFGERNVKGMFEVLEPLHAMMERGPQTLKETSFNQAYG
RDLMEAQEWCRKYMKSGNVKDLTQAWDLYYHVFRRISKQQISYASRGGGSSGGGELAAIRKK
LVIVGDGA*

>DNA sequence

ATGAAAATCGAAGAAGGTAAACTGGTAATCTGGATTAACGGCGATAAAGGCTATAACGGTCT
CGCTGAAGTCGGTAAGAAATTCGAGAAAGATACCGGAATTAAAGTCACCGTTGAGCATCCG
GATAAACTGGAAGAGAAATTCCCACAGGTTGCGGCAACTGGCGATGGCCCTGACATTATCT
TCTGGGCACACGACCGCTTTGGTGGCTACGCTCAATCTGGCCTGTTGGCTGAAATCACCCC
GGACAAAGCGTTCCAGGACAAGCTGTATCCGTTTACCTGGGATGCCGTACGTTACAACGGC
AAGCTGATTGCTTACCCGATCGCTGTTGAAGCGTTATCGCTGATTTATAACAAAGATCTGCTG
CCGAACCCGCCAAAAACCTGGGAAGAGATCCCGGCGCTGGATAAAGAACTGAAAGCGAAA
GGTAAGAGCGCGCTGATGTTCAACCTGCAAGAACCGTACTTCACCTGGCCGCTGATTGCTG
CTGACGGGGGTTATGCGTTCAAGTATGAAAACGGCAAGTACGACATTAAAGACGTGGGCGT
GGATAACGCTGGCGCGAAAGCGGGTCTGACCTTCCTGGTTGACCTGATTAAAAACAAACAC
ATGAATGCAGACACCGATTACTCCATCGCAGAAGCTGCCTTTAATAAAGGCGAAACAGCGAT
GACCATCAACGGCCCGTGGGCATGGTCCAACATCGACACCAGCAAAGTGAATTATGGTGTA
ACGGTACTGCCGACCTTCAAGGGTCAACCATCCAAACCGTTCGTTGGCGTGCTGAGCGCA
GGTATTAACGCCGCCAGTCCGAACAAAGAGCTGGCAAAAGAGTTCCTCGAAAACTATCTGC
TGACTGATGAAGGTCTGGAAGCGGTTAATAAAGACAAACCGCTGGGTGCCGTAGCGCTGAA
GTCTTACGAGGAAGAGTTGGTGAAAGATCCGCGTATTGCCGCCACTATGGAAAACGCCCAG
AAAGGTGAAATCATGCCGAACATCCCGCAGATGTCCGCTTTCTGGTATGCCGTGCGTACTG
CGGTGATCAACGCCGCCAGCGGTCGTCAGACTGTCGATGAAGCCCTGAAAGACGCGCAGA
CTAATTCGAGCTCGAACAACAACAACAATAACAATAACAACAACCTCGGGATCGAGGGAAGG
ATTTCACATATGTCCATGGGCGGCCGCGAGATGTGGCATGAAGGCCTGGAAGAGGCATCTC
GTTTGTACTTTGGGGAAAGGAACGTGAAAGGCATGTTTGAGGTGCTGGAGCCCTTGCATGC
TATGATGGAACGGGGCCCCCAGACTCTGAAGGAAACATCCTTTAATCAGGCCTATGGTCGA
GATTTAATGGAGGCCCAAGAGTGGTGCAGGAAGTACATGAAATCAGGGAATGTCAAGGACC
TCACCCAAGCCTGGGACCTCTATTATCATGTGTTCCGACGAATCTCAAAGCAGCAGATCTCG
TACGCGTCCCGGGGCGGTGGCTCATCTGGCGGAGGTGAGCTCGCTGCCATCCGGAAGAA
ACTGGTGATTGTTGGTGATGGAGCCTAA

12）**pET21b-RhoA/13C-FKBP-MBP**

>Amino acid sequence

MCGKTCLLIVFSKDQFPEVYVPTVFENYVADIEVDGKQVELALWDTAGLEDYDRLRPLSYPDTD
VILMCFSIDSPDSLENIPEKWTPEVKHFCPNVPIILVGNKKDLRNDEHTRRELAKMKQEPVKPEE
GRDMANRIGAFGYMECSAKTKDGVREVFEMATRAALQAGDPNWELVYTARLQGGGSSGGGQ
ISYASRGGVQVETISPGDGRTFPKRGQTCVVHYTGMLEDGKKFDSSRDRNKPFKFMLGKQEVI
RGWEEGVAQMSVGQRAKLTISPDYAYGATGHPGIIPPHATLVFDVELLKLEKLENLYFQGEEGKL
VIWINGDKGYNGLAEVGKKFEKDTGIKVTVEHPDKLEEKFPQVAATGDGPDIIFWAHDRFGGYA
QSGLLAEITPDKAFQDKLYPFTWDAVRYNGKLIAYPIAVEALSLIYNKDLLPNPPKTWEEIPALDK
ELKAKGKSALMFNLQEPYFTWPLIAADGGYAFKYENGKYDIKDVGVDNAGAKAGLTFLVDLIKN
KHMNADTDYSIAEAAFNKGETAMTINGPWAWSNIDTSKVNYGVTVLPTFKGQPSKPFVGVLSA
GINAASPNKELAKEFLENYLLTDEGLEAVNKDKPLGAVALKSYEEELAKDPRIAATMENAQKGEI
MPNIPQMSAFWYAVRTAVINAASGRQTVDEALKDAQTNSSS*

>DNA sequence

ATGTGTGGAAAGACATGCTTGCTCATAGTCTTCAGCAAGGATCAGTTCCCAGAGGTGTATGT
GCCCACAGTGTTTGAGAACTATGTGGCAGATATCGAGGTGGATGGAAAGCAGGTAGAGTTG
GCTTTGTGGGACACAGCTGGGCTGGAAGATTATGATCGCCTGAGGCCCCTCTCCTACCCAG
ATACCGATGTTATACTGATGTGTTTTTCCATCGACAGCCCTGATAGTTTAGAAAACATCCCAG
AAAAGTGGACCCCAGAAGTCAAGCATTTCTGTCCCAACGTGCCCATCATCCTGGTTGGGAA
TAAGAAGGATCTTCGGAATGATGAGCACACAAGGCGGGAGCTAGCCAAGATGAAGCAGGA
GCCGGTGAAACCTGAAGAAGGCAGAGATATGGCAAACAGGATTGGCGCTTTTGGGTACATG
GAGTGTTCAGCAAAGACCAAAGATGGAGTGAGAGAGGTTTTTGAAATGGCTACGAGAGCTG
CTCTGCAAGCTGGGGATCCCAATTGGGAGCTCGTGTACACGGCGCGCCTGCAGGGAGGT
GGCTCATCTGGCGGAGGTCAGATCTCGTACGCGTCCCGGGGCGGAGTGCAGGTGGAAAC
CATCTCCCCAGGAGACGGGCGCACCTTCCCCAAGCGCGGCCAGACCTGCGTGGTGCACT
ACACCGGGATGCTTGAAGATGGAAAGAAATTTGATTCCTCCCGGGACAGAAACAAGCCCTT
TAAGTTTATGCTAGGCAAGCAGGAGGTGATCCGAGGCTGGGAAGAAGGGGTTGCCCAGAT
GAGTGTGGGTCAGAGAGCCAAACTGACTATATCTCCAGATTATGCCTATGGTGCCACTGGG
CACCCAGGCATCATCCCACCACATGCCACTCTCGTCTTCGATGTGGAGCTTCTAAAACTGG
AAAAGCTTGAGAACCTGTACTTCCAGGGCGAAGAAGGTAAACTGGTAATCTGGATTAACGG
CGATAAAGGCTATAACGGTCTCGCTGAAGTCGGTAAGAAATTCGAGAAAGATACCGGAATTA
AAGTCACCGTTGAGCATCCGGATAAACTGGAAGAGAAATTCCCACAGGTTGCGGCAACTGG
CGATGGCCCTGACATTATCTTCTGGGCACACGACCGCTTTGGTGGCTACGCTCAATCTGGC
CTGTTGGCTGAAATCACCCCGGACAAAGCGTTCCAGGACAAGCTGTATCCGTTTACCTGGG
ATGCCGTACGTTACAACGGCAAGCTGATTGCTTACCCGATCGCTGTTGAAGCGTTATCGCTG
ATTTATAACAAAGACCTGCTGCCGAACCCGCCAAAAACCTGGGAAGAGATCCCGGCGCTGG
ATAAAGAACTGAAAGCGAAAGGTAAGAGCGCGCTGATGTTCAACCTGCAAGAACCGTACTT
CACCTGGCCGCTGATTGCTGCTGACGGGGGTTATGCGTTCAAGTATGAAAACGGCAAGTAC
GACATTAAAGACGTGGGCGTGGATAACGCTGGCGCGAAAGCGGGTCTGACCTTCCTGGTT
GACCTGATTAAAAACAAACACATGAATGCAGACACCGATTACTCCATCGCAGAAGCTGCCTT
TAATAAAGGCGAAACAGCGATGACCATCAACGGCCCGTGGGCATGGTCCAACATCGACACC
AGCAAAGTGAATTATGGTGTAACGGTACTGCCGACCTTCAAGGGTCAACCATCCAAACCGTT
CGTTGGCGTGCTGAGCGCAGGTATTAACGCCGCCAGTCCGAACAAAGAGCTGGCAAAAGA
GTTCCTCGAAAACTATCTGCTGACTGATGAAGGTCTGGAAGCGGTTAATAAAGACAAACCGC

TGGGTGCCGTAGCGCTGAAGTCTTACGAGGAAGAGTTGGCGAAAGATCCACGTATTGCCG
CCACTATGGAAAACGCCCAGAAAGGTGAAATCATGCCGAACATCCCGCAGATGTCCGCTTT
CTGGTATGCCGTGCGTACTGCGGTGATCAACGCCGCCAGCGGTCGTCAGACTGTCGATGA
AGCCCTGAAAGACGCGCAGACTAATTCCAGCTCGTAG

13）**pMAL-MBP-KRas-Q61L**

>Amino acid sequence

MKIEEGKLVIWINGDKGYNGLAEVGKKFEKDTGIKVTVEHPDKLEEKFPQVAATGDGPDIIFWAH
DRFGGYAQSGLLAEITPDKAFQDKLYPFTWDAVRYNGKLIAYPIAVEALSLIYNKDLLPNPPKTW
EEIPALDKELKAKGKSALMFNLQEPYFTWPLIAADGGYAFKYENGKYDIKDVGVDNAGAKAGLT
FLVDLIKNKHMNADTDYSIAEAAFNKGETAMTINGPWAWSNIDTSKVNYGVTVLPTFKGQPSKP
FVGVLSAGINAASPNKELAKEFLENYLLTDEGLEAVNKDKPLGAVALKSYEEELVKDPRIAATME
NAQKGEIMPNIPQMSAFWYAVRTAVINAASGRQTVDEALKDAQTNSSSNNNNNNNNNNLGIEG
RISHMMTEYKLVVVGAGGVGKSALTIQLIQNHFVDEYDPTIEDSYRKQVVIDGETCLLDILDTAGL
EEYSAMRDQYMRTGEGFLCVFAINNTKSFEDIHHYREQIKRVKDSEDVPMVLVGNKCDLPSRT
VDTKQAQDLARSYGIPFIETSAKTRQGVDDAFYTLVREIRKHKEKMSKDGKKKKKKSKTKCVIM*

>DNA sequence

ATGAAAATCGAAGAAGGTAAACTGGTAATCTGGATTAACGGCGATAAAGGCTATAACGGTCT
CGCTGAAGTCGGTAAGAAATTCGAGAAAGATACCGGAATTAAAGTCACCGTTGAGCATCCG
GATAAACTGGAAGAGAAATTCCCACAGGTTGCGGCAACTGGCGATGGCCCTGACATTATCT
TCTGGGCACACGACCGCTTTGGTGGCTACGCTCAATCTGGCCTGTTGGCTGAAATCACCCC
GGACAAAGCGTTCCAGGACAAGCTGTATCCGTTTACCTGGGATGCCGTACGTTACAACGGC
AAGCTGATTGCTTACCCGATCGCTGTTGAAGCGTTATCGCTGATTTATAACAAAGATCTGCTG
CCGAACCCGCCAAAAACCTGGGAAGAGATCCCGGCGCTGGATAAAGAACTGAAAGCGAAA
GGTAAGAGCGCGCTGATGTTCAACCTGCAAGAACCGTACTTCACCTGGCCGCTGATTGCTG
CTGACGGGGGTTATGCGTTCAAGTATGAAAACGGCAAGTACGACATTAAAGACGTGGGCGT
GGATAACGCTGGCGCGAAAGCGGGTCTGACCTTCCTGGTTGACCTGATTAAAAACAAACAC
ATGAATGCAGACACCGATTACTCCATCGCAGAAGCTGCCTTTAATAAAGGCGAAACAGCGAT
GACCATCAACGGCCCGTGGGCATGGTCCAACATCGACACCAGCAAAGTGAATTATGGTGTA
ACGGTACTGCCGACCTTCAAGGGTCAACCATCCAAACCGTTCGTTGGCGTGCTGAGCGCA
GGTATTAACGCCGCCAGTCCGAACAAAGAGCTGGCAAAAGAGTTCCTCGAAAACTATCTGC
TGACTGATGAAGGTCTGGAAGCGGTTAATAAAGACAAACCGCTGGGTGCCGTAGCGCTGAA
GTCTTACGAGGAAGAGTTGGTGAAAGATCCGCGTATTGCCGCCACTATGGAAAACGCCCAG
AAAGGTGAAATCATGCCGAACATCCCGCAGATGTCCGCTTTCTGGTATGCCGTGCGTACTG
CGGTGATCAACGCCGCCAGCGGTCGTCAGACTGTCGATGAAGCCCTGAAAGACGCGCAGA
CTAATTCGAGCTCGAACAACAACAACAATAACAATAACAACAACCTCGGGATCGAGGGAAGG
ATTTCACATATGATGACTGAATATAAACTTGTGGTAGTTGGAGCTGGTGGCGTAGGCAAGAG
TGCCTTGACGATACAGCTAATTCAGAATCATTTTGTGGACGAATATGATCCAACAATAGAGGA
TTCCTACAGGAAGCAAGTAGTAATTGATGGAGAAACCTGTCTCTTGGATATTCTCGACACAG
CAGGTCTGGAGGAGTACAGTGCAATGAGGGACCAGTACATGAGGACTGGGGAGGGCTTTC
TTTGTGTATTTGCCATAAATAATACTAAATCATTTGAAGATATTCACCATTATAGAGAACAAATTA
AAAGAGTTAAGGACTCTGAAGATGTACCTATGGTCCTAGTAGGAAATAAATGTGATTTGCCTT
CCAGAACAGTAGACACAAAACAGGCTCAGGACTTAGCAAGAAGTTATGGAATTCCTTTTATT
GAAACATCAGCAAAGACAAGACAGGGTGTTGATGATGCCTTCTATACATTAGTTCGAGAAAT
TCGAAAACATAAAGAAAAGATGAGCAAAGATGGTAAAAAGAAGAAAAAGAAGTCAAAGACAA
AGTGTGTAATTATGTAA

14）**pMAL-MBP-KRas-S17N**

>Amino acid sequence

MKIEEGKLVIWINGDKGYNGLAEVGKKFEKDTGIKVTVEHPDKLEEKFPQVAATGDGPDIIFWAH
DRFGGYAQSGLLAEITPDKAFQDKLYPFTWDAVRYNGKLIAYPIAVEALSLIYNKDLLPNPPKTW
EEIPALDKELKAKGKSALMFNLQEPYFTWPLIAADGGYAFKYENGKYDIKDVGVDNAGAKAGLT
FLVDLIKNKHMNADTDYSIAEAAFNKGETAMTINGPWAWSNIDTSKVNYGVTVLPTFKGQPSKP
FVGVLSAGINAASPNKELAKEFLENYLLTDEGLEAVNKDKPLGAVALKSYEEELVKDPRIAATME
NAQKGEIMPNIPQMSAFWYAVRTAVINAASGRQTVDEALKDAQTNSSSNNNNNNNNNNLGIEG
RISHMMTEYKLVVVGAGGVGKNALTIQLIQNHFVDEYDPTIEDSYRKQVVIDGETCLLDILDTAG
QEEYSAMRDQYMRTGEGFLCVFAINNTKSFEDIHHYREQIKRVKDSEDVPMVLVGNKCDLPSR
TVDTKQAQDLARSYGIPFIETSAKTRQGVDDAFYTLVREIRKHKEKMSKDGKKKKKKSKTKCVI
M*

>DNA sequence

ATGAAAATCGAAGAAGGTAAACTGGTAATCTGGATTAACGGCGATAAAGGCTATAACGGTCT
CGCTGAAGTCGGTAAGAAATTCGAGAAAGATACCGGAATTAAAGTCACCGTTGAGCATCCG
GATAAACTGGAAGAGAAATTCCCACAGGTTGCGGCAACTGGCGATGGCCCTGACATTATCT
TCTGGGCACACGACCGCTTTGGTGGCTACGCTCAATCTGGCCTGTTGGCTGAAATCACCCC
GGACAAAGCGTTCCAGGACAAGCTGTATCCGTTTACCTGGGATGCCGTACGTTACAACGGC
AAGCTGATTGCTTACCCGATCGCTGTTGAAGCGTTATCGCTGATTTATAACAAAGATCTGCTG
CCGAACCCGCCAAAAACCTGGGAAGAGATCCCGGCGCTGGATAAAGAACTGAAAGCGAAA
GGTAAGAGCGCGCTGATGTTCAACCTGCAAGAACCGTACTTCACCTGGCCGCTGATTGCTG
CTGACGGGGGTTATGCGTTCAAGTATGAAAACGGCAAGTACGACATTAAAGACGTGGGCGT
GGATAACGCTGGCGCGAAAGCGGGTCTGACCTTCCTGGTTGACCTGATTAAAAACAAACAC
ATGAATGCAGACACCGATTACTCCATCGCAGAAGCTGCCTTTAATAAAGGCGAAACAGCGAT
GACCATCAACGGCCCGTGGGCATGGTCCAACATCGACACCAGCAAAGTGAATTATGGTGTA
ACGGTACTGCCGACCTTCAAGGGTCAACCATCCAAACCGTTCGTTGGCGTGCTGAGCGCA
GGTATTAACGCCGCCAGTCCGAACAAAGAGCTGGCAAAAGAGTTCCTCGAAAACTATCTGC
TGACTGATGAAGGTCTGGAAGCGGTTAATAAAGACAAACCGCTGGGTGCCGTAGCGCTGAA
GTCTTACGAGGAAGAGTTGGTGAAAGATCCGCGTATTGCCGCCACTATGGAAAACGCCCAG
AAAGGTGAAATCATGCCGAACATCCCGCAGATGTCCGCTTTCTGGTATGCCGTGCGTACTG
CGGTGATCAACGCCGCCAGCGGTCGTCAGACTGTCGATGAAGCCCTGAAAGACGCGCAGA
CTAATTCGAGCTCGAACAACAACAACAATAACAATAACAACAACCTCGGGATCGAGGGAAGG
ATTTCACATATGATGACTGAATATAAACTTGTGGTAGTTGGAGCTGGTGGCGTAGGCAAGAAT
GCCTTGACGATACAGCTAATTCAGAATCATTTTGTGGACGAATATGATCCAACAATAGAGGAT
TCCTACAGGAAGCAAGTAGTAATTGATGGAGAAACCTGTCTCTTGGATATTCTCGACACAGC
AGGTCAAGAGGAGTACAGTGCAATGAGGGACCAGTACATGAGGACTGGGGAGGGCTTTCT
TTGTGTATTTGCCATAAATAATACTAAATCATTTGAAGATATTCACCATTATAGAGAACAAATTAA
AAGAGTTAAGGACTCTGAAGATGTACCTATGGTCCTAGTAGGAAATAAATGTGATTTGCCTTC
TAGAACAGTAGACACAAAACAGGCTCAGGACTTAGCAAGAAGTTATGGAATTCCTTTTATTGA
AACATCAGCAAAGACAAGACAGGGTGTTGATGATGCCTTCTATACATTAGTTCGAGAAATTC
GAAAACATAAAGAAAAGATGAGCAAAGATGGTAAAAAGAAGAAAAAGAAGTCAAAGACAAAG
TGTGTAATTATGTAA

15）**pMAL-MBP-FRB-KRas/N12**

>Amino acid sequence

MKIEEGKLVIWINGDKGYNGLAEVGKKFEKDTGIKVTVEHPDKLEEKFPQVAATGDGPDIIFWAH
DRFGGYAQSGLLAEITPDKAFQDKLYPFTWDAVRYNGKLIAYPIAVEALSLIYNKDLLPNPPKTW
EEIPALDKELKAKGKSALMFNLQEPYFTWPLIAADGGYAFKYENGKYDIKDVGVDNAGAKAGLT
FLVDLIKNKHMNADTDYSIAEAAFNKGETAMTINGPWAWSNIDTSKVNYGVTVLPTFKGQPSKP
FVGVLSAGINAASPNKELAKEFLENYLLTDEGLEAVNKDKPLGAVALKSYEEELVKDPRIAATME
NAQKGEIMPNIPQMSAFWYAVRTAVINAASGRQTVDEALKDAQTNSSSNNNNNNNNNNLGIEG
RISHMEMWHEGLEEASRLYFGERNVKGMFEVLEPLHAMMERGPQTLKETSFNQAYGRDLMEA
QEWCRKYMKSGNVKDLTQAWDLYYHVFRRISKQQISYASRGGGSSGGGELMTEYKLVVVGAG
*

>DNA sequence

ATGAAAATCGAAGAAGGTAAACTGGTAATCTGGATTAACGGCGATAAAGGCTATAACGGTCT
CGCTGAAGTCGGTAAGAAATTCGAGAAAGATACCGGAATTAAAGTCACCGTTGAGCATCCG
GATAAACTGGAAGAGAAATTCCCACAGGTTGCGGCAACTGGCGATGGCCCTGACATTATCT
TCTGGGCACACGACCGCTTTGGTGGCTACGCTCAATCTGGCCTGTTGGCTGAAATCACCCC
GGACAAAGCGTTCCAGGACAAGCTGTATCCGTTTACCTGGGATGCCGTACGTTACAACGGC
AAGCTGATTGCTTACCCGATCGCTGTTGAAGCGTTATCGCTGATTTATAACAAAGATCTGCTG
CCGAACCCGCCAAAAACCTGGGAAGAGATCCCGGCGCTGGATAAAGAACTGAAAGCGAAA
GGTAAGAGCGCGCTGATGTTCAACCTGCAAGAACCGTACTTCACCTGGCCGCTGATTGCTG
CTGACGGGGGTTATGCGTTCAAGTATGAAAACGGCAAGTACGACATTAAAGACGTGGGCGT
GGATAACGCTGGCGCGAAAGCGGGTCTGACCTTCCTGGTTGACCTGATTAAAAACAAACAC
ATGAATGCAGACACCGATTACTCCATCGCAGAAGCTGCCTTTAATAAAGGCGAAACAGCGAT
GACCATCAACGGCCCGTGGGCATGGTCCAACATCGACACCAGCAAAGTGAATTATGGTGTA
ACGGTACTGCCGACCTTCAAGGGTCAACCATCCAAACCGTTCGTTGGCGTGCTGAGCGCA
GGTATTAACGCCGCCAGTCCGAACAAAGAGCTGGCAAAAGAGTTCCTCGAAAACTATCTGC
TGACTGATGAAGGTCTGGAAGCGGTTAATAAAGACAAACCGCTGGGTGCCGTAGCGCTGAA
GTCTTACGAGGAAGAGTTGGTGAAAGATCCGCGTATTGCCGCCACTATGGAAAACGCCCAG
AAAGGTGAAATCATGCCGAACATCCCGCAGATGTCCGCTTTCTGGTATGCCGTGCGTACTG
CGGTGATCAACGCCGCCAGCGGTCGTCAGACTGTCGATGAAGCCCTGAAAGACGCGCAGA
CTAATTCGAGCTCGAACAACAACAACAATAACAATAACAACAACCTCGGGATCGAGGGAAGG
ATTTCACATATGGAGATGTGGCATGAAGGCCTGGAAGAGGCATCTCGTTTGTACTTTGGGGA
AAGGAACGTGAAAGGCATGTTTGAGGTGCTGGAGCCCTTGCATGCTATGATGGAACGGGG
CCCCCAGACTCTGAAGGAAACATCCTTTAATCAGGCCTATGGTCGAGATTTAATGGAGGCCC
AAGAGTGGTGCAGGAAGTACATGAAATCAGGGAATGTCAAGGACCTCACCCAAGCCTGGG
ACCTCTATTATCATGTGTTCCGACGAATCTCAAAGCAGCAGATCTCGTACGCGTCCCGGGGC
GGTGGCTCATCTGGCGGAGGTGAGCTCATGACTGAATATAAACTTGTGGTAGTTGGAGCTG
GTTAA

16） **pET21b-KRas/13C-FKBP-MBP**

>Amino acid sequence

MGVGKSALTIQLIQNHFVDEYDPTIEDSYRKQVVIDGETCLLDILDTAGLEEYSAMRDQYMRTGE
GFLCVFAINNTKSFEDIHHYREQIKRVKDSEDVPMVLVGNKCDLPSRTVDTKQAQDLARSYGIP
FIETSAKTRQGVDDAFYTLVREIRKHKEKMSKDGGDPNWELVYTARLQGGGSSGGGQISYASR
GGVQVETISPGDGRTFPKRGQTCVVHYTGMLEDGKKFDSSRDRNKPFKFMLGKQEVIRGWEE
GVAQMSVGQRAKLTISPDYAYGATGHPGIIPPHATLVFDVELLKLEKLENLYFQGEEGKLVIWING
DKGYNGLAEVGKKFEKDTGIKVTVEHPDKLEEKFPQVAATGDGPDIIFWAHDRFGGYAQSGLLA
EITPDKAFQDKLYPFTWDAVRYNGKLIAYPIAVEALSLIYNKDLLPNPPKTWEEIPALDKELKAKG
KSALMFNLQEPYFTWPLIAADGGYAFKYENGKYDIKDVGVDNAGAKAGLTFLVDLIKNKHMNAD
TDYSIAEAAFNKGETAMTINGPWAWSNIDTSKVNYGVTVLPTFKGQPSKPFVGVLSAGINAASP
NKELAKEFLENYLLTDEGLEAVNKDKPLGAVALKSYEEELAKDPRIAATMENAQKGEIMPNIPQM
SAFWYAVRTAVINAASGRQTVDEALKDAQTNSSS*

>DNA sequence

ATGGGCGTAGGCAAGAGTGCCTTGACGATACAGCTAATTCAGAATCATTTTGTGGACGAATA
TGATCCAACAATAGAGGATTCCTACAGGAAGCAAGTAGTAATTGATGGAGAAACCTGTCTCT
TGGATATTCTCGACACAGCAGGTCTGGAGGAGTACAGTGCAATGAGGGACCAGTACATGAG
GACTGGGGAGGGCTTTCTTTGTGTATTTGCCATAAATAATACTAAATCATTTGAAGATATTCAC
CATTATAGAGAACAAATTAAAAGAGTTAAGGACTCTGAAGATGTACCTATGGTCCTAGTAGGA
AATAAATGTGATTTGCCTTCCAGAACAGTAGACACAAAACAGGCTCAGGACTTAGCAAGAAG
TTATGGAATTCCTTTTATTGAAACATCAGCAAAGACAAGACAGGGTGTTGATGATGCCTTCTA
TACATTAGTTCGAGAAATTCGAAAACATAAAGAAAAGATGAGCAAAGATGGTGGGGATCCCA
ATTGGGAGCTCGTGTACACGGCGCGCCTGCAGGGAGGTGGCTCATCTGGCGGAGGTCAG
ATCTCGTACGCGTCCCGGGGCGGAGTGCAGGTGGAAACCATCTCCCCAGGAGACGGGCG
CACCTTCCCCAAGCGCGGCCAGACCTGCGTGGTGCACTACACCGGGATGCTTGAAGATGG
AAAGAAATTTGATTCCTCCCGGGACAGAAACAAGCCCTTTAAGTTTATGCTAGGCAAGCAGG
AGGTGATCCGAGGCTGGGAAGAAGGGGTTGCCCAGATGAGTGTGGGTCAGAGAGCCAAA
CTGACTATATCTCCAGATTATGCCTATGGTGCCACTGGGCACCCAGGCATCATCCCACCACA
TGCCACTCTCGTCTTCGATGTGGAGCTTCTAAAACTGGAAAAGCTTGAGAACCTGTACTTCC
AGGGCGAAGAAGGTAAACTGGTAATCTGGATTAACGGCGATAAAGGCTATAACGGTCTCGCT
GAAGTCGGTAAGAAATTCGAGAAAGATACCGGAATTAAAGTCACCGTTGAGCATCCGGATAA
ACTGGAAGAGAAATTCCCACAGGTTGCGGCAACTGGCGATGGCCCTGACATTATCTTCTGG
GCACACGACCGCTTTGGTGGCTACGCTCAATCTGGCCTGTTGGCTGAAATCACCCCGGAC
AAAGCGTTCCAGGACAAGCTGTATCCGTTTACCTGGGATGCCGTACGTTACAACGGCAAGC
TGATTGCTTACCCGATCGCTGTTGAAGCGTTATCGCTGATTTATAACAAAGACCTGCTGCCG
AACCCGCCAAAAACCTGGGAAGAGATCCCGGCGCTGGATAAAGAACTGAAAGCGAAAGGT
AAGAGCGCGCTGATGTTCAACCTGCAAGAACCGTACTTCACCTGGCCGCTGATTGCTGCTG
ACGGGGGTTATGCGTTCAAGTATGAAAACGGCAAGTACGACATTAAAGACGTGGGCGTGGA
TAACGCTGGCGCGAAAGCGGGTCTGACCTTCCTGGTTGACCTGATTAAAAACAAACACATG
AATGCAGACACCGATTACTCCATCGCAGAAGCTGCCTTTAATAAAGGCGAAACAGCGATGAC
CATCAACGGCCCGTGGGCATGGTCCAACATCGACACCAGCAAAGTGAATTATGGTGTAACG
GTACTGCCGACCTTCAAGGGTCAACCATCCAAACCGTTCGTTGGCGTGCTGAGCGCAGGT
ATTAACGCCGCCAGTCCGAACAAAGAGCTGGCAAAAGAGTTCCTCGAAAACTATCTGCTGA
CTGATGAAGGTCTGGAAGCGGTTAATAAAGACAAACCGCTGGGTGCCGTAGCGCTGAAGTC

TTACGAGGAAGAGTTGGCGAAAGATCCACGTATTGCCGCCACTATGGAAAACGCCCAGAAA
GGTGAAATCATGCCGAACATCCCGCAGATGTCCGCTTTCTGGTATGCCGTGCGTACTGCGG
TGATCAACGCCGCCAGCGGTCGTCAGACTGTCGATGAAGCCCTGAAAGACGCGCAGACTA
ATTCCAGCTCGTAG

**Table S4. Molecular weights and extinction coefficients of recombinant proteins.**

| Protein Name | Molecular Weight (kDa) | Extinction Coefficient ($M^{-1}\cdot cm^{-1}$) |
|---|---|---|
| MBP-Cdc42-Q61L | 63.2 | 82,530 |
| MBP-Cdc42-T17N | 63.3 | 82,530 |
| MBP-FRB-Cdc42/N12 | 56.5 | 91,915 |
| Cdc42/13C-FKBP-MBP | 75.5 | 102,470 |
| MBP-Rac1-Q61L | 62.9 | 89,645 |
| MBP-Rac1-T17N | 63.0 | 89,645 |
| MBP-FRB-Rac1/N12 | 57.0 | 91,915 |
| Rac1/13C-FKBP-MBP | 75.3 | 109,585 |
| MBP-RhoA-Q63L | 63.3 | 85,050 |
| MBP-RhoA-T19N | 63.3 | 85,050 |
| MBP-FRB-RhoA/N12 | 57.3 | 91,790 |
| RhoA/13C-FKBP-MBP | 75.3 | 105,115 |
| MBP-KRas-Q61L | 64.4 | 78,520 |
| MBP-KRas-S17N | 64.4 | 78,520 |
| MBP-FRB-KRas/N12 | 56.6 | 93,280 |
| KRas/13C-FKBP-MBP | 74.9 | 96,970 |
